# Transferring CRISPRi knockdown responses to new cell lines requires screening part of each line

**DOI:** 10.64898/2026.11.15.766815

**Authors:** Md Kawsar, Sajib Acharjee Dip, Bikash Kumar Paul, Abdullah Al Noman, Liqing Zhang, Mohammad Ali Moni

## Abstract

Perturb-seq screens measure knockdown responses in one cell line at a time. Models that transfer responses to new lines rarely beat simple averages, and standard metrics often ignore noise. It is unclear how much of a response is shared and what a new line must contribute. We studied CRISPR interference Perturb-seq of essential genes in four human cell lines, using noise ceilings and calibrated metrics. We also introduce CASTER, a set-based model that adapts the transferred mean to the target line. After noise correction, responses correlated at a median of 0.52, highest between the *TP53* wild-type lines. With 30% of a line screened, a linear model and CASTER scored 0.84 and 0.80 versus 0.44 for the transferred mean (1 equals a technical duplicate). CASTER led only at 2.5% screened (0.62 versus 0.47) and fell below the transferred mean without a screen. Response size tracked general essentiality more than line-specific dependency.

**Highlights:**

- Knockdown responses correlate at about 0.5 between lines after noise correction
- Knockdowns that activate p53 in HepG2 and RPE1 have more similar responses
- A partial screen lets a linear model adapt responses as well as a set-based model
- Response size tracks general gene essentiality more than line-specific dependency

**In brief:** Kawsar et al. analyze CRISPRi Perturb-seq of essential genes in four human cell lines and find that knockdown responses correlate at about 0.5 between lines after noise correction. Screening part of a new line lets a linear model adapt transferred responses, whereas control cells alone do not.

## Introduction

Pooled CRISPR screens with single-cell transcriptomic readouts (Perturb-seq) measure the transcriptome-wide effect of a genetic perturbation in thousands of cells at once^1–4^. CRISPR interference (CRISPRi)^5,6^ has made such screens scalable to every expressed or essential gene^7,8^. They are now used to map gene function, genetic interactions and regulatory programs^8–11^. In cancer they have found regulators of immune evasion, drug response and oncogenic variants^12–16^. Each screen, however, is run in one cellular context. Cancers differ in lineage, drivers, ploidy and baseline state, which shapes how a cell responds to losing a gene^17–20^. Screening every perturbation in every relevant context is not feasible, so models that transfer perturbation responses across contexts are needed^21,22^.

Several classes of models predict single-cell perturbation responses. Autoencoder models encode cells in a latent space and model perturbations there^23–27^. Optimal-transport and flow-matching models map control cells to perturbed cells^28,29^. Graph-based models use gene networks to predict unseen perturbations^30,31^. Single-cell foundation models are pretrained on many cells and adapted to perturbation tasks^32–35^. Sensitivity analysis of inferred cell-fate trajectories identifies the genes whose perturbation most changes fate^36^. State, a transformer trained on sets of cells with a distributional loss, transferred effects to new contexts better than these models^37^. But benchmarks show that simple baselines, such as a perturbation’s mean response across training contexts or a linear model, are hard to beat^38–40^. Responses shared by all perturbations in a context can also inflate standard metrics^41^. A community challenge now asks for virtual-cell models that generalize to held-out contexts^42^. It therefore remains unclear how much of a knockdown’s response is specific to a line, and what a new line must contribute before a learned model beats a transferred average.

Here we ask which parts of a knockdown response carry over between cell lines. We use one essential-gene CRISPRi library profiled in K562 (chronic myeloid leukemia), HepG2 (hepatoblastoma), Jurkat (T-cell acute lymphoblastic leukemia) and non-transformed hTERT RPE-1 cells (RPE1)^8,43^. We also introduce CASTER (Context-Adaptive Set Transformer for pErturbation Responses), a model that starts from a knockdown’s mean response in the other lines and corrects it for the target line (Figure 1E).

**Figure 1.**
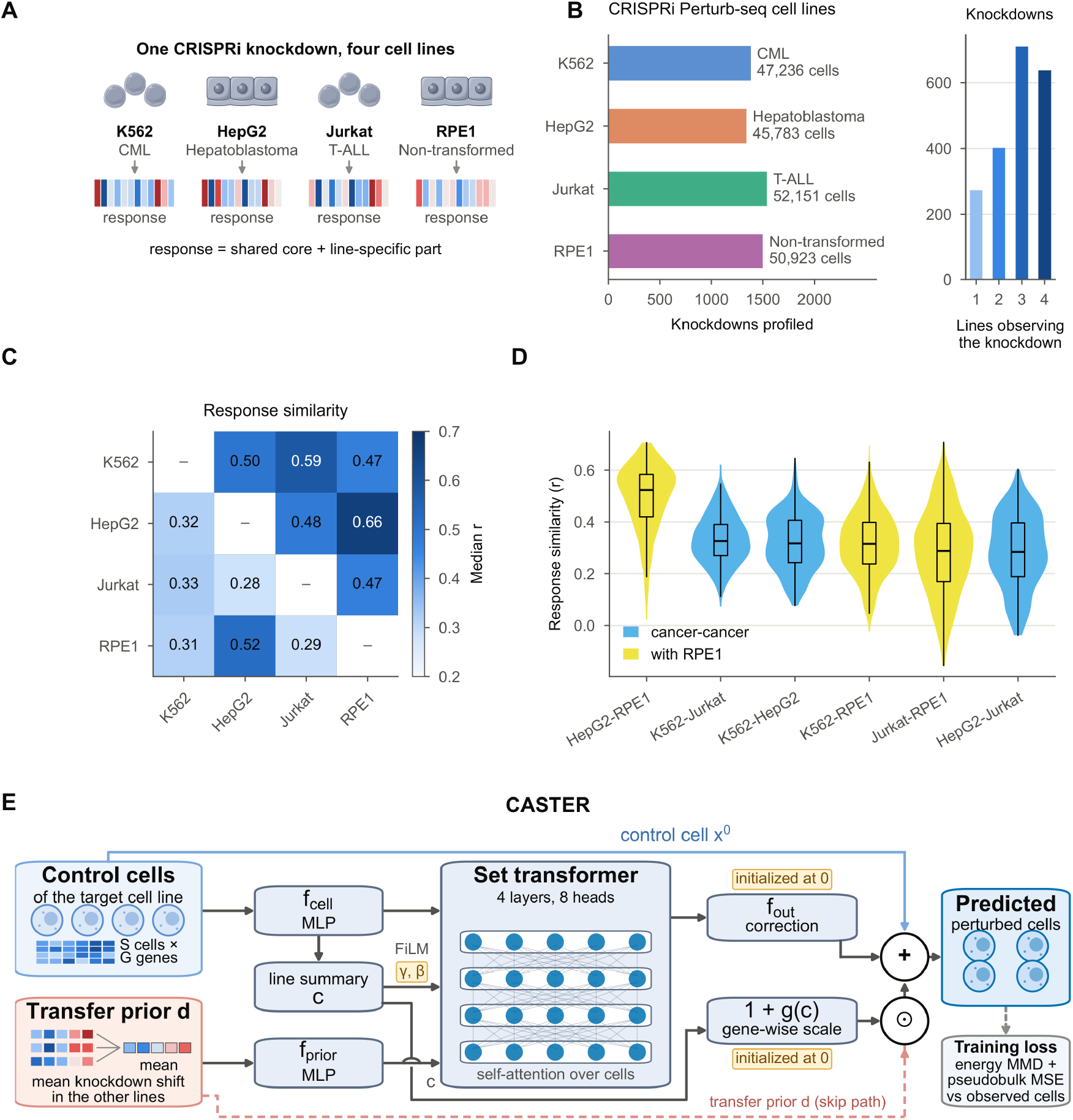
Knockdown responses differ between cell lines and the CASTER model. (A) The same CRISPRi knockdown, profiled in three cancer lines and a non-transformed line, produces a shared core response plus a line-specific component. (B) Knockdowns retained per line, with cell numbers (left), and lines retaining each knockdown (right). (C) Response similarity: median Pearson correlation of observed response shifts between lines (knockdowns with ≥ 10 DEGs in both; n per pair in Table S15); raw values below the diagonal, values corrected for measurement noise above it. (D) Per-knockdown response similarity for each pair of lines (same knockdowns as in C); cancer-cancer pairs in sky blue, pairs involving RPE1 in yellow. (E) CASTER. The transfer prior d is scaled gene-wise by 1 + g(c) (⊙, element-wise product) and added to each control cell with a correction from a FiLM-conditioned set transformer; both start at zero (STAR Methods). Cell icons: Servier Medical Art (smart.servier.com), CC BY 3.0, recolored. See also Figures S1, S2, S3 and S4 and Tables S1, S15 and S3.

We first measure how similar responses to the same knockdown are between lines and correct this similarity for noise using split-half noise ceilings (Figures 1C and 1D). To score predictions, we use the calibrated metrics of Miller et al.^44^. These compare a prediction with a technical duplicate and with a prediction that carries no knockdown-specific information.

CASTER reads a knockdown’s mean shift in the other lines and a set of the target line’s unperturbed cells. Unlike a single offset per line, it can rescale each gene and correct each cell. We test it with part of the new line screened and with only its control cells, against the transferred mean, a per-line offset and a linear model.

After noise correction, responses were moderately shared and most similar between the two TP53 wild-type lines. With part of the new line screened, CASTER and the linear model both beat the transferred mean, and the linear model did as well as CASTER. Control cells alone did not adapt responses. Response size tracked general essentiality in DepMap Chronos scores^45,46^ more than line-specific dependency. Finally, a small screen of the new line, not a richer model of its control cells, is what lets knockdown responses be carried to a new cell line.

## Results

### Knockdown responses correlate at about 0.5 between cell lines after noise correction

We assembled CRISPRi Perturb-seq profiles of essential-gene knockdowns in four human cell lines (Figures 1A and 1B). They are K562 (chronic myeloid leukemia; *TP53* null)^8^, HepG2 (hepatoblastoma; *TP53* wild type)^43^, Jurkat (T-cell acute lymphoblastic leukemia; *TP53* mutant)^43^ and RPE1 (non-transformed; *TP53* wild type)^8,47^. We used the knockdown-filtered, uniformly processed release of these data^37^ and analyzed 196,093 cells: up to 32 cells per knockdown per line and 3,000 controls per line (STAR Methods; Table S1). After filtering, 1,383 (K562), 1,340 (HepG2), 1,537 (Jurkat) and 1,499 (RPE1) knockdowns remained (2,023 distinct target genes); 638 were retained in all four lines and 782 in all three cancer lines. Because on-target efficacy was filtered per line, the set of informative knockdowns is itself line dependent (Figure 1B). We modeled 2,000 genes selected on control cells only, with all knockdown targets excluded (STAR Methods).

The number of differentially expressed genes (DEGs; Wilcoxon rank-sum test against held-out controls, false discovery rate [FDR] < 0.05) varied widely between knockdowns and between lines (median 2 to 29; Table S1; Figures S1A and S1B) and were largest in the two *TP53* wild-type lines, RPE1 and HepG2. For 706 genes whose knockdown produced ≥ 10 DEGs in at least two lines, we compared response profiles. They were only moderately correlated: median Pearson r was 0.31 between cancer lines and 0.38 between a cancer line and RPE1 (Figures 1C and 1D), and DEG sets overlapped little (median Jaccard index 0.08 and 0.10; Figure S1C). Before correction, the most similar pair was HepG2-RPE1 (median raw r = 0.52); the other five pairs ranged from 0.28 to 0.33. Nadig et al. reported noise-corrected cross-line correlations of perturbation profiles in these lines with pair medians of 0.40 (K562-RPE1) to 0.75 (HepG2-RPE1), HepG2-RPE1 being the most similar pair^43^. Our raw values are lower because they include measurement noise at 32 cells per knockdown. We corrected each knockdown’s cross-line r for noise by dividing it by the square root of the two within-line split-half reliabilities (Spearman’s correction for attenuation; knockdowns with ≥ 10 DEGs in both lines; STAR Methods). The median rose from 0.31 to 0.52 between cancer lines and from 0.38 to 0.56 between a cancer line and RPE1. After correction, the two leukemia lines were the second most similar pair (K562-Jurkat 0.59). Each of the two top pairs joins one line from each source study, so the study of origin alone does not set the ranking. After correction, responses were moderately correlated, but a large line-specific part remained. The next sections ask how much of this part can be predicted.

Similarity rose with response size, from 0.31 for knockdowns with ≥ 10 DEGs to 0.45 for those with ≥ 200 DEGs in both cancer lines (Figure S1D). Part of the apparent line specificity of weak responses is therefore measurement noise. The noise ceiling is the highest Pearson delta (the Pearson correlation between predicted and observed mean shifts) any predictor can reach against the observed response (STAR Methods). With 32 cells per knockdown, its mean was 0.51 to 0.69 across lines (Table S2), so absolute accuracies should be read against these ceilings. Because HepG2 and RPE1 are the two *TP53* wild-type lines, we scored the activity of 14 consensus p53 target genes^48^ for every knockdown (Figure S2). p53 targets were induced by 28.4% of knockdowns in HepG2 and 47.1% in RPE1, and by almost none in K562 and Jurkat (0.1% or less). The p53 response to single knockdowns was shared most strongly between HepG2 and RPE1 (Spearman rho = 0.63, versus 0.06 to 0.21 for the other pairs). Knockdowns inducing p53 in both lines had more similar HepG2 and RPE1 responses (median r = 0.57 versus 0.45; Mann-Whitney p = 1.9 × 10⁻²²). These results link shared p53 activation with response similarity. They do not show that p53 causes the similarity.

### CASTER starts from the transferred mean response and adapts it to the target line

These observations motivated CASTER (Figure 1E). Like State^37^, CASTER operates on *sets* of cells and is trained with a distributional (energy-distance maximum mean discrepancy [MMD]) loss^49,50^, so it models a perturbed population rather than a single average cell. A perturbation is represented by its *transfer prior*: the mean shift it produced in the other lines in which it was observed. The target line is never part of this prior. The target line is represented by a set of its unperturbed cells, read by a transformer encoder; feature-wise linear modulation (FiLM)^51^ lets the same prior act differently in each line. The output adds a line-dependent, gene-wise rescaling of the prior and a cell-resolved correction. Both start at zero, so an untrained CASTER equals PerturbMean, and training moves it away from this starting point (STAR Methods; hyperparameters in Table S3). In simulations, CASTER learned line-specific structure only where it existed, as did a linear transfer model (Figure S3). Ablations on the K562 run showed that the transfer prior mattered most and that attention and FiLM mattered little (Figure S4; Table S4; Note S1).

### A partial screen of the target line lets learned models beat mean baselines

We first evaluated the setting in which a new line has been screened with only part of a library (Figure 2A)^37^. For each of the four lines in turn, 25% of its eligible knockdowns were used for training, 5% for model selection and 70% for testing; the other three lines were fully observed. We compared CASTER with five baselines. PerturbMean is a knockdown’s mean response in the other lines. ContextMean is the target line’s mean response over its screened knockdowns. LineOffset is PerturbMean plus the target line’s mean deviation from it. Linear is a linear transfer model fitted on the screened knockdowns. NoChange predicts zero shift. Our main metric is the scaled score of Miller et al.^44^ (0 = NoChange, 1 = technical duplicate) on top-100 Pearson delta (Pearson delta on the 100 genes with the largest absolute t statistic, relative to the perturbed mean), scored against all cells of each knockdown (Figure 2B); other metrics follow Cell-Eval^37^ (STAR Methods; training curves in Figure S5). We repeated the task with three random seeds, each with its own split and initialization.

**Figure 2.**
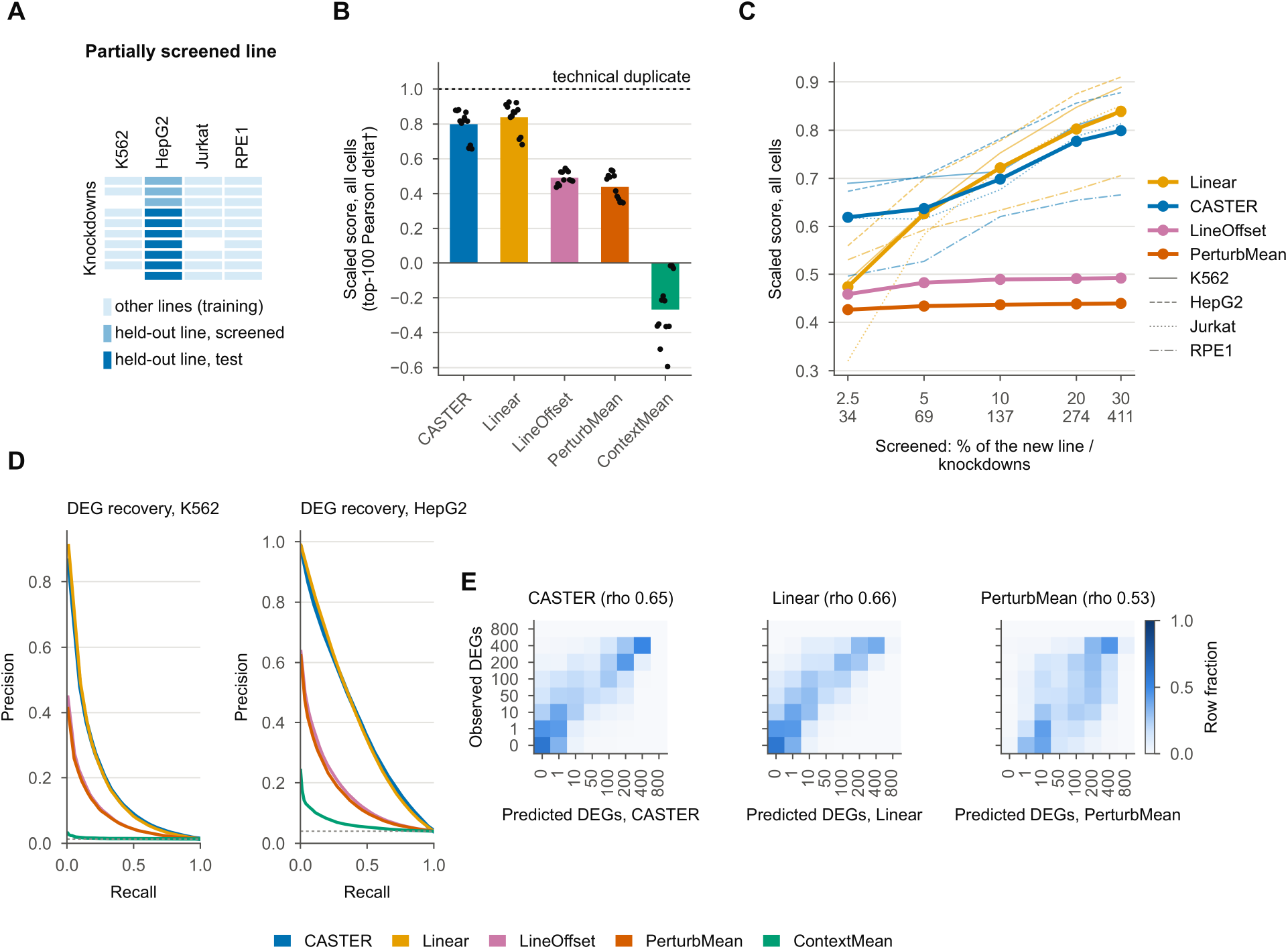
A partial screen lets learned models beat mean baselines. (A) Task: for each target line, 25% of its knockdowns train, 5% validate and 70% test; the other lines are fully observed. (B) Scaled top-100 Pearson delta (relative to the perturbed mean) against all cells of each held-out knockdown, from NoChange (0) to the technical duplicate (1; dashed). Noise-free predictions; bars, mean over four lines and three seeds; dots, line-seed runs (n = 12). (C) The same score for nested screened sets of 2.5% to 30% of the target line (fixed 70% test set); thick lines, mean over lines and seeds; thin lines, per line. Values in Table S5. (D) Precision-recall curves for recovering observed DEGs (32 cells, first seed; test knockdowns of K562, n = 917, and HepG2, n = 894); dashed line, base rate. (E) Observed versus predicted number of DEGs per knockdown (row-normalized) for CASTER, Linear and PerturbMean (all lines, first seed, n = 3,841 knockdowns). Method colors as in the key below. See also Figures S5, S6, S7, S16 and S8 and Tables S16, S4, S17 and S5.

Averaged over three seeds and four lines, Pearson delta was 0.28 for Linear, 0.28 for CASTER, 0.25 for LineOffset, 0.21 for ContextMean and 0.20 for PerturbMean (standard deviation [SD] over seeds below 0.005). Per-line values are in Figure S6 and secondary metrics in Figure S7. CASTER therefore improved on the mean baselines in every seed but not on Linear, and much of its average gain over PerturbMean came from RPE1, where PerturbMean was weak.

We next asked where CASTER gains over the transfer prior. The gain grew with response size in every line (Spearman rho 0.22 to 0.64; Figure S8): the median gain in Pearson delta was 0.09 to 0.33 for knockdowns with ≥ 50 DEGs and 0.00 to 0.03 for those with < 10 DEGs. No such gradient appeared in the unseen-line task.

We then measured how much of a new line must be screened. We kept the same 70% of knockdowns as the test set and trained on nested screened sets of 2.5%, 5%, 10%, 20% and 30% of the library (three seeds; STAR Methods), scoring top-100 Pearson delta against all cells (Figure 2C). Linear’s scaled score rose from 0.47 at 2.5% to 0.84 at 30% (0.63, 0.72 and 0.80 at 5%, 10% and 20%). CASTER’s rose from 0.62 to 0.80 (0.64, 0.70 and 0.78). PerturbMean, which uses no data from the line, stayed at 0.43 to 0.44 (Table S5). On average, Linear exceeded PerturbMean from 2.5% screened on (not in Jurkat at 2.5%: 0.32 versus 0.52), and reached 91% of its 30% gain over PerturbMean at 20%. The gain was still rising at 30%, so these data do not give a smallest useful screen size. CASTER was ahead of Linear when the screen was smallest: at 2.5% (about 34 screened knockdowns) it scored higher in the three cancer lines, by 0.11 to 0.30, but not in RPE1 (−0.03; 11 of 12 line-seed runs). Its lead shrank as more knockdowns were screened (ahead in 7 of 12 runs at 5% and 3 of 12 runs at 10%) and was gone from 20% on. Because CASTER starts from PerturbMean and is trained on all source-line knockdowns, whereas Linear is fitted from scratch on the screened ones, we ran four controls at small screen sizes (2.5% shown here; 5% and 10% in Table S5). A ridge on the residual from PerturbMean reached 0.56, PerturbMean with a gene-wise scale 0.50, Linear with 8 or 16 components 0.43 and 0.46, and the mean-profile CASTER 0.63 (CASTER 0.62, Linear 0.47; Table S5). Starting from the transferred mean therefore explained most of CASTER’s lead over Linear. CASTER stayed above the anchored ridge in the three cancer lines but not in RPE1. The mean-profile CASTER did as well as full CASTER, so the lead did not need cell-to-cell variation among control cells. We did not test whether the remaining gap comes from training on source-line knockdowns or from CASTER’s non-linear form.

The learned models also recovered the observed DEGs better than PerturbMean (area under the precision-recall curve for differential expression [DE AUPRC] 0.19 for CASTER, 0.18 for Linear and 0.14 for PerturbMean, mean over seeds and lines; Figure 2D) and ranked knockdowns by their number of DEGs more accurately (Spearman rho pooled over lines 0.65, 0.66 and 0.53; Figure 2E).

### Adaptation goes beyond a single offset per cell line

Per-line Pearson delta was stable across seeds and splits (Figure 3A). ContextMean alone captures much of the response in RPE1, so CASTER’s gain could be little more than one constant shift per line. We tested this with four controls (Figure 3B shows the first three). First, LineOffset adds the target line’s mean deviation from PerturbMean to every prediction. Its Pearson delta was 0.251, versus 0.278 for CASTER (first seed; CASTER better for 56% of knockdowns, two-sided paired Wilcoxon p = 3.1 × 10⁻⁸⁴). Second, the mean-profile CASTER receives the line’s mean control profile instead of a set of cells. It reached 0.271 versus 0.278 for full CASTER (first seed, same split; p = 1.1 × 10⁻²⁸). Third, the trained CASTER given another line’s control cells reached 0.151 versus 0.275 with the correct cells (mean over two seeds; paired Wilcoxon p < 10⁻³⁰⁰). Fourth, we trained CASTER with only a line label: the mean control profile with its genes shuffled in a fixed order per line. It reached 0.274, against 0.271 with the mean profile and 0.278 for full CASTER (first seed, same splits). On the scaled top-100 Pearson delta against all cells (first seed, noise-free predictions; Table S6), full CASTER scored 0.80, the mean-profile CASTER 0.84, the line-label CASTER 0.81, LineOffset 0.50 and Linear 0.83; the wrong-line CASTER scored 0.36 versus 0.80 with the correct cells (two seeds). On this score, the mean-profile and line-label models were not below full CASTER. Control cells, and even the mean control profile, therefore added nothing we could measure, and a line label was enough. Linear sees no control cells at all and does as well. This suggests that the adaptation comes mainly from the screened knockdowns of the line.

**Figure 3.**
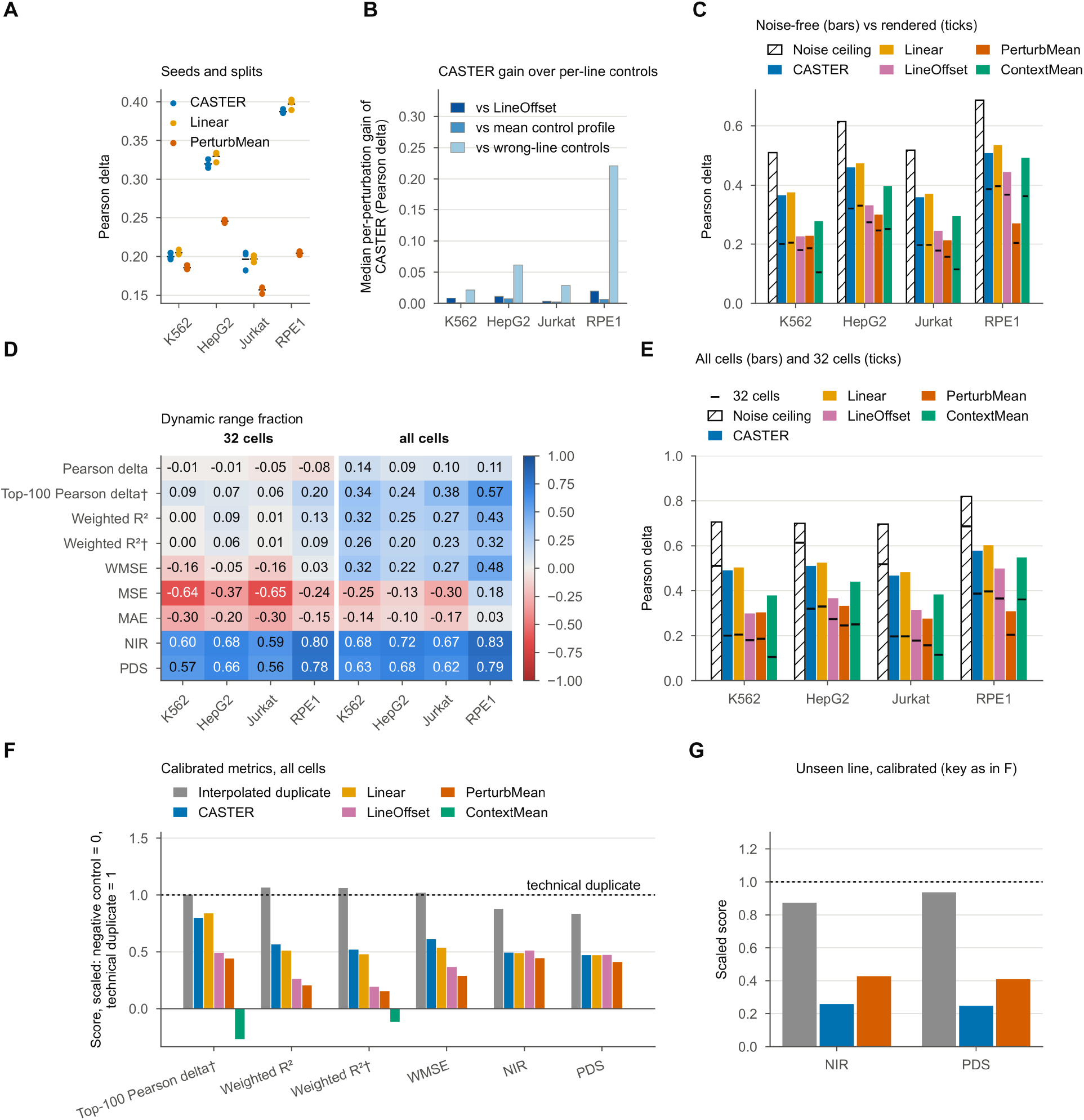
Controls and calibrated metrics for the partial-screen task. (A) Pearson delta per target line and seed (dots; tick, mean) for CASTER, Linear and PerturbMean (32 cells; three seeds). (B) Median per-knockdown gain of CASTER over LineOffset, over the mean-profile CASTER, and over CASTER given another line’s control cells (32-cell Pearson delta, as rendered; on the calibrated score the mean-profile model is not below CASTER, Table S6; first seed; n = test knockdowns per line, 894 to 1,027). (C) Pearson delta (32 cells) with noise-free predictions (bars; mean over seeds), as rendered (ticks) and the noise ceiling (hatched). (D) Dynamic range fraction of each metric per line at 32 cells and with all cells: how well the metric separates the technical duplicate from the negative control (−1 to 1). † relative to the perturbed mean. (E) Pearson delta against all cells (bars; noise-free, mean over seeds) and against 32 cells (ticks), with the noise ceiling (hatched; tick, 32-cell ceiling). (F) Calibrated metrics against all cells on a scale from the negative control (0) to the technical duplicate (1; dashed): 0 = NoChange for the metrics relative to the perturbed mean (†), 0 = the perturbed mean for NIR, PDS, WMSE and weighted R². Metrics with DRF ≥ 0.2 with all cells; not shown: Pearson delta, MSE and MAE. Mean over lines and seeds (Table S9). (G) Calibrated metrics in the unseen-line task (32 cells, one seed; scale as in F). See also Tables S7, S8, S9, S2, S18 and S6.

Every baseline predicts one mean shift per knockdown. To score all methods the same way, this shift was added to sampled control cells, as many as the knockdown has observed cells (at most 32; we call this rendering), so each prediction inherits the sampling noise of those cells. CASTER sees these cells and could learn to remove that noise. We therefore rescored every method after removing this noise (STAR Methods). Removing it raised CASTER, Linear and ContextMean by about 0.15 in Pearson delta and PerturbMean and LineOffset by about 0.06 (Figure 3C). CASTER therefore does not gain by canceling input noise; rendering on sampled control cells lowered the absolute accuracy of every method, most for those that use the target line’s own data. Further robustness checks, including similarity on all 6,546 measured genes, are in Table S7.

### Scoring against all cells raises noise ceilings and shows Linear and CASTER close

Miller et al. showed that several standard metrics barely separate a perfect from a useless prediction on these four datasets, and proposed calibrating metrics with positive and negative controls^44,52^. We therefore rescored every held-out knockdown against one random half of its cells. The other half served as a technical duplicate (positive control), and an interpolated duplicate as a second positive control. The negative control was the perturbed mean (the mean response of the line’s screened knockdowns, which is also the ContextMean prediction), or a zero shift (NoChange) for metrics that are already relative to the perturbed mean (STAR Methods). The dynamic range fraction (DRF; mean of clipped per-knockdown values) is 0 when a metric cannot separate the two controls and 1 when it separates them fully. Averaged over lines, it was 0.67 for the normalized inverse rank (NIR) and 0.64 for the perturbation discrimination score (PDS). PDS differs from the perturbation discrimination score of Equation 13, which ranks pseudobulk profiles by L1 distance among all test knockdowns. The DRF was near zero or negative for Pearson delta, top-100 Pearson delta, weighted R², weighted R² relative to the perturbed mean, weighted mean squared error (WMSE), mean squared error (MSE) and mean absolute error (MAE) (Figure 3D). With only 16 cells per half, the technical duplicate is no closer to the truth than the perturbed mean on these metrics, so they cannot validate a predictor at this cell count. We call a metric calibrated when its DRF is 0.2 or more, and report scaled scores (0 = negative control, 1 = technical duplicate) only for those; a score above 1 is possible because a noise-free prediction can beat a noisy 16-cell duplicate. On the two calibrated metrics (NIR and PDS), CASTER scored 0.49 and 0.48. Linear scored 0.48 and 0.47, LineOffset 0.50 and 0.48 and PerturbMean 0.43 and 0.42 (32 cells as rendered; Tables S8 and S9). With 32 cells, the reference response is as noisy as the training data. We therefore rescored the trained models, without retraining, against all available cells of each held-out knockdown (median 57 cells; STAR Methods). The mean noise ceiling of Pearson delta rose by 0.09 to 0.20 in the four lines (to 0.70 to 0.82; Figure 3E; Table S2). On the scaled top-100 Pearson delta against all cells, Linear scored 0.84, CASTER 0.80, LineOffset 0.49 and PerturbMean 0.44 (Figures 2B and 3F; Table S9). CASTER scored above Linear on WMSE (0.61 versus 0.54), and the two were level on NIR (0.49 versus 0.49) and PDS (0.47 versus 0.47; Figure 3F). On every calibrated score, LineOffset was above PerturbMean (Tables S8 and S9).

### Control cells alone did not let CASTER adapt responses to an unseen line

We next asked whether responses could be transferred into a line with no perturbation data at all, using only its unperturbed cells (Figure 4A). With this model, they could not. As rendered at 32 cells, CASTER was worse than PerturbMean in 21 of 24 line-by-metric panels (Pearson delta, discrimination, DE AUPRC, DE overlap, fold-change Spearman and effect-size Spearman, four lines; one seed; Figures 4B and S9). PerturbMean was significantly better on all three main metrics (Table S10): Pearson delta 0.197 versus 0.171 (p = 2.4 × 10⁻¹⁰⁵), discrimination 0.363 versus 0.214 (p = 2.3 × 10⁻²⁶⁹) and DE AUPRC 0.136 versus 0.107 (p = 2.9 × 10⁻⁴⁵). After removing rendering noise, Pearson delta was level (0.26 for CASTER and 0.25 for PerturbMean, mean over lines), but discrimination stayed far lower (0.22 versus 0.38; Table S10). CASTER therefore lost the ability to tell knockdowns apart and gained nothing clear in return. One possible reason is that corrections learned between source lines pull predictions toward a response shared by all knockdowns. We did not test this. On the calibrated metrics, CASTER also scored below PerturbMean (NIR 0.26 versus 0.43 and PDS 0.25 versus 0.41; Figure 3G).

**Figure 4.**
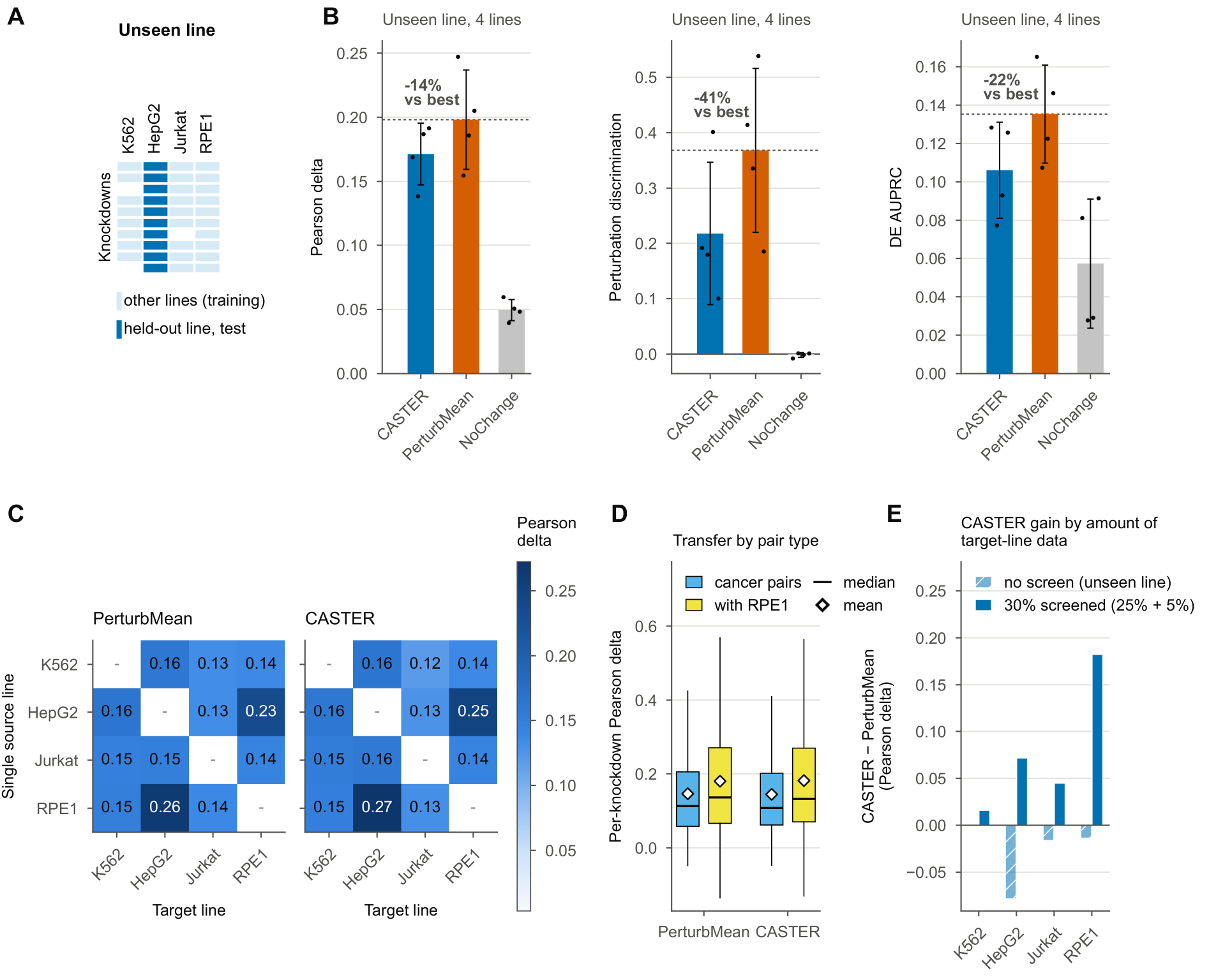
Control cells alone do not adapt responses to an unseen line. (A) Unseen-line task: no perturbed cells of the target line are used. (B) Main metrics in the unseen-line task, as rendered at 32 cells (noise-free values in Table S10; one seed; bars, mean over lines; error bars, SD over lines; dots, lines, n = 4). Dashed line, best baseline; percentage, relative difference of CASTER from it. (C) Single-source transfer: Pearson delta for every source (rows) and target (columns) line; shared color scale. (D) Per-knockdown transfer accuracy averaged over cancer-to-cancer directions versus directions involving RPE1 (n = 1,204 knockdowns observed in RPE1 and in at least two cancer lines). Boxes, interquartile range (IQR); line, median; diamond, mean; whiskers, 1.5 × IQR; two-sided paired Wilcoxon test (PerturbMean), p value in the text. (E) CASTER minus PerturbMean (Pearson delta, as rendered at 32 cells) per line, with no screen (unseen line) or with 30% of the target line screened (25% training, 5% validation). See also Figures S2, S5, S7, S9 and S10 and Tables S10 and S11.

To test whether transfer depends on how related two lines are, we transferred every knockdown from a single source line to each other line (12 directions; Figure 4C). With the PerturbMean transfer, per-knockdown accuracy (Pearson delta, mean over knockdowns) averaged over cancer-to-cancer directions was 0.14, versus 0.17 for directions involving RPE1 (n = 1,204 knockdowns observed in RPE1 and in at least two cancer lines, paired Wilcoxon p = 9.5 × 10⁻⁵¹; Figure 4D). Contrary to our expectation, transfer involving the non-transformed line was more accurate. The difference came mostly from the HepG2-RPE1 pair (Pearson delta 0.23 from HepG2 to RPE1 and 0.26 from RPE1 to HepG2, the two highest entries of the matrix). This matches the higher response similarity of these two adherent, epithelial-derived lines (Figure 1C) and the larger, better-resolved responses of RPE1 (Figure S1A). In single-source transfer CASTER was close to PerturbMean on Pearson delta (worse in 6 of 12 directions, 1 tie) but worse in 12 of 12 on discrimination and 12 of 12 on DE overlap (Figures 4C and S10; Table S11). CASTER gained over PerturbMean only when part of the target line was screened (Figure 4E).

### Gains concentrate on strong responses such as ribosome biogenesis knockdowns

Per knockdown (first seed, four lines pooled), CASTER had a higher DE overlap than PerturbMean for 31% of knockdowns, the same for 36% and a lower one for 33%; its fold-change agreement was higher than PerturbMean’s for 60% and than Linear’s for 45% (Figures 5A and 5B). Figure 5C shows the knockdown with the largest CASTER gain over PerturbMean in a cancer line; Figure S11 shows the best and the worst case in each cancer line, with Linear for comparison. Predicted shifts of S-phase and G2/M signature scores^53^ had mean absolute errors of 0.06 (S) and 0.08 (G2/M) for CASTER, against 0.09 and 0.14 for PerturbMean and 0.06 and 0.08 for Linear (Figure 5D). When knockdowns were clustered by the similarity of their responses, predicted similarity maps recovered the observed structure with mean adjusted Rand indices (over 2 to 20 clusters and four lines) of 0.26 (CASTER), 0.23 (Linear), 0.10 (PerturbMean) and 0.02 (ContextMean) (Figure 5E shows K562).

**Figure 5.**
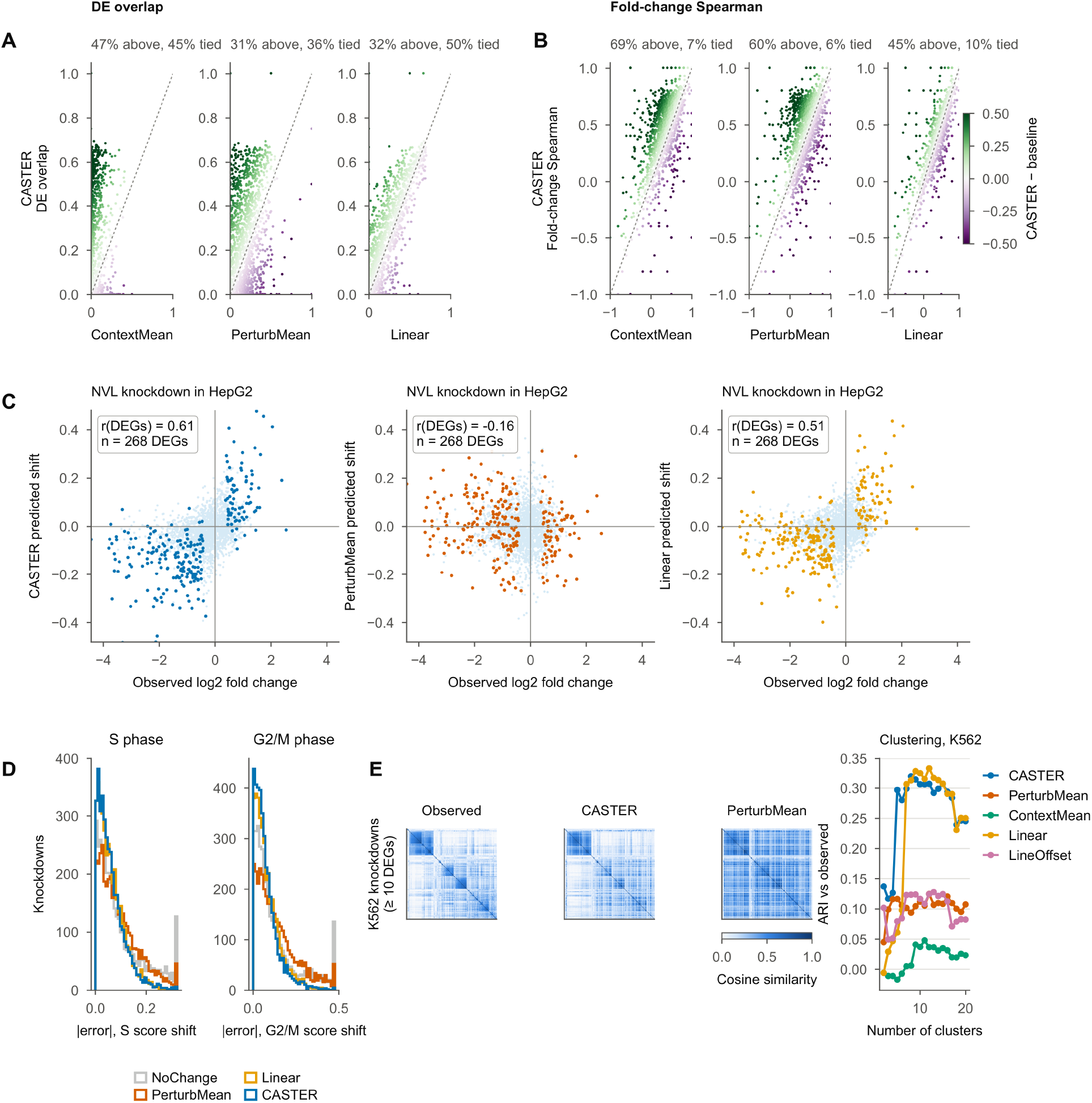
Gains concentrate on strong responses. (A and B) Per held-out knockdown (four lines pooled, first seed), CASTER (y) versus each baseline (x) for DE overlap (A; n = 2,666 knockdowns with observed DEGs) and fold-change Spearman (B; n = 2,094); color, CASTER minus baseline (green, CASTER better). Panel titles, share of knockdowns above the diagonal and tied. (C) Observed log₂ fold change versus predicted shift for the knockdown with the largest CASTER gain over PerturbMean in a cancer line (colored, observed DEGs; r over DEGs). (D) Absolute error of predicted S-phase and G2/M signature shifts (n = 3,841 knockdowns, first seed); values above the 99.5th percentile are in the last bin. (E) Knockdown-similarity (cosine) maps for K562 knockdowns with ≥ 10 DEGs and adjusted Rand index between predicted and observed clusterings (first seed). See also Figures S11 and S12 and Table S12.

We next asked which knockdowns CASTER adapts most. In each cancer line, the knockdown with the largest gain over PerturbMean (among those with ≥ 30 DEGs) was a regulator of ribosome biogenesis or a ribosomal protein: *NVL* in HepG2, *RPL26* in Jurkat and *RPL6* in K562 (r over DEGs for CASTER versus PerturbMean: 0.61 versus −0.16, 0.71 versus 0.32 and 0.60 versus 0.36; Figures 5C and S11). Ribosome knockdowns cause strong, line-dependent stress responses^54–56^. Because CASTER’s gain on them was similar in the p53-deficient lines (K562 and Jurkat) and in HepG2, p53 alone does not explain it. To test this beyond single examples, we used all held-out knockdowns. After adjusting for response size, the gain over PerturbMean was larger for ribosome-biogenesis and ribosomal-protein knockdowns in every line (first seed; Figure S12A). The regression coefficient was 0.149 in HepG2, 0.130 in Jurkat, 0.125 in K562 and 0.077 in RPE1 (all p ≤ 2.5 × 10⁻⁴). Linear gained similarly on these knockdowns (coefficients 0.06 to 0.13), whereas LineOffset (−0.03 to 0.02) and ContextMean (−0.03 to 0.05) did not, so the models that learn from screened knockdowns gain most on these strong, line-dependent responses. Holding out whole complexes also tests how much the scores depend on related genes being screened. Under complex-grouped splits (three seeds, scaled top-100 Pearson delta against all cells), CASTER and Linear scored 0.73 and 0.73, LineOffset 0.45 and PerturbMean 0.41 (random splits: 0.80, 0.84, 0.49 and 0.44). Because whole groups moved between training and test, the test sets differ; on the knockdowns held out under both splits (about 650 per run), CASTER and Linear scored 0.74 and 0.73 with grouped and 0.79 and 0.82 with random splits. Linear’s lead on random splits disappeared under grouped splits (CASTER ahead in 6 of 12 line-seed runs): Linear lost 0.09 and CASTER 0.05, so part of Linear’s advantage on random splits may come from related genes in training. Because the perturbed mean depends on which knockdowns are screened, PerturbMean’s scores also moved although it uses no target data. On the raw top-100 Pearson delta, CASTER minus Linear rose by 0.02 (bootstrap 95% interval 0.02 to 0.03; scaled: 0.04, 0.03 to 0.05; Table S12). This shift is small.

### Response size reflects shared gene essentiality rather than line-specific dependency

Finally, we asked whether response size relates to how much a line depends on the gene, measured independently by genome-scale CRISPR fitness screens in DepMap as the Chronos gene effect (Chronos score)^45,46,57^. In each cancer line, knockdowns of genes on which the line depends more strongly (more negative Chronos score) were expected to produce larger transcriptional responses. Response size correlated with the Chronos score in all three lines: rho = −0.45 in K562 (n = 1,325), −0.54 in HepG2 (n = 1,275) and −0.39 in Jurkat (n = 1,465) (all Holm p ≤ 6.8 × 10⁻⁵⁴; all knockdowns of each line with a Chronos score; Figure 6A). We then asked whether the line-specific part of the response tracks the line-specific part of the dependency. For genes profiled in all three cancer lines, each part is the line’s value minus the mean of the other two. The correlation was weak: rho = −0.12 in K562 (Holm p = 0.004), −0.05 in HepG2 (Holm p = 0.318) and −0.05 in Jurkat (Holm p = 0.318) (Figure 6B). The coupling was not line-specific. A line’s response size correlated about as strongly with another line’s Chronos scores as with its own (own: −0.45 in K562, −0.54 in HepG2 and −0.39 in Jurkat; other lines, mean: −0.40, −0.49 and −0.38; Figure S13). The mean Chronos score of the other two lines predicted each line’s own score with Pearson r = 0.82, 0.85 and 0.83. The response-dependency link therefore reflects general gene essentiality: knockdowns of broadly essential genes cause large transcriptional responses in every line tested.

**Figure 6.**
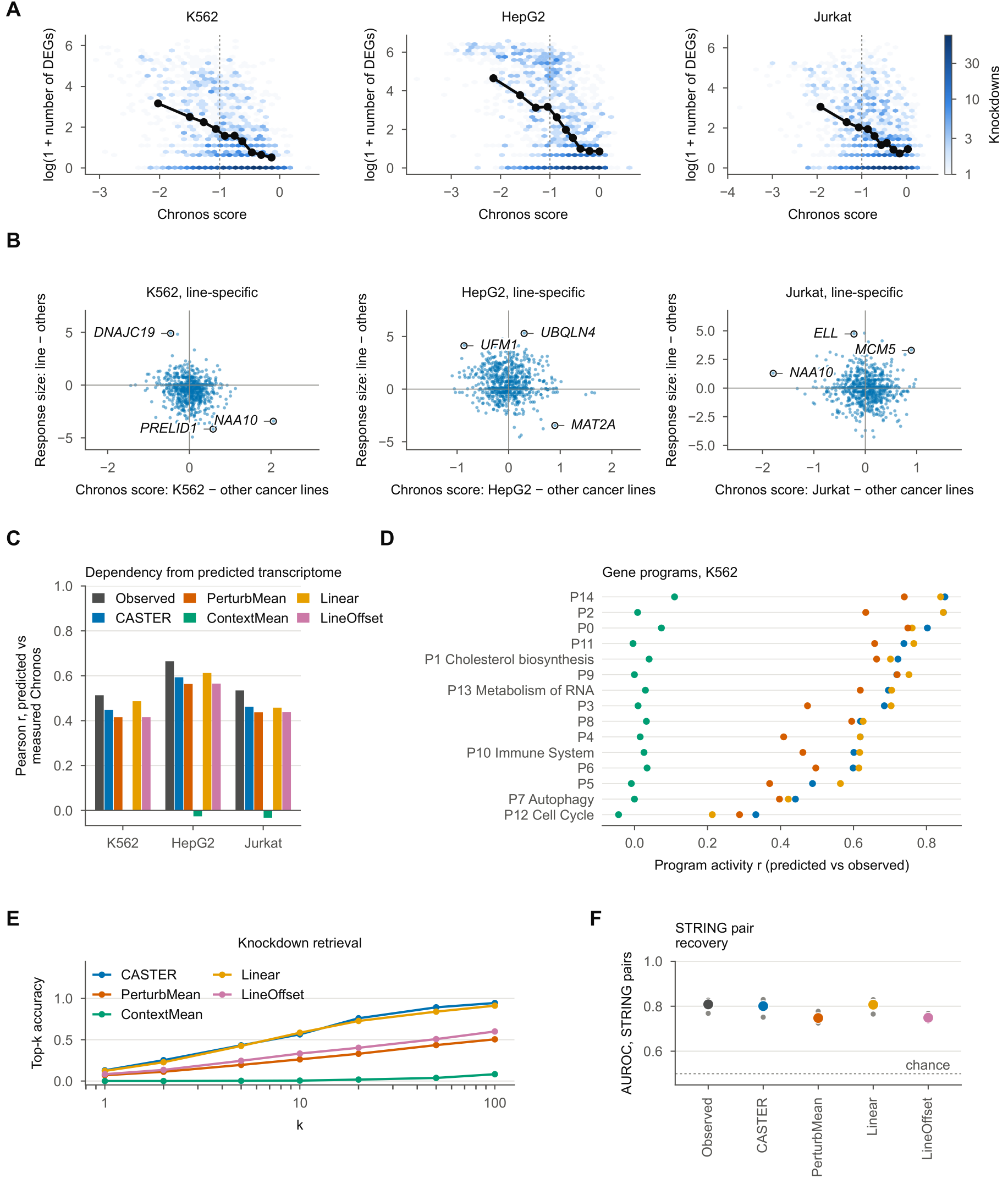
Response size reflects shared gene essentiality. (A) Response size (log(1 + number of DEGs)) versus Chronos score of the knocked-down gene in each cancer line (all knockdowns with a Chronos score; n, Spearman rho and Holm-adjusted p in the text); black line, decile means; dashed line, Chronos score −1 (typical essential gene). (B) Line-specific response versus line-specific dependency (line minus mean of the other two cancer lines; genes profiled in all three). Labeled, most extreme genes. (C) Pearson r between measured Chronos scores and Chronos scores predicted by a ridge model (trained on observed shifts) from observed or predicted shifts (held-out knockdowns with a Chronos score). (D) Concordance of predicted and observed gene-program activity across knockdowns (K562; programs as in Figure S15, top Reactome term where FDR < 0.05). (E) Knockdown retrieval: top-k accuracy of identifying the knockdown that produced an observed response among all held-out candidates of a line (about 960), for the 100 strongest knockdowns per line. (F) AUROC for separating STRING-linked (combined score ≥ 700) knockdown pairs by response similarity (knockdowns with ≥ 10 DEGs); large dots, mean over lines; small dots, lines; dashed line, chance. Method colors in (E) and (F) as in (C); observed responses in black outline. See also Figures S13, S17, S14 and S15 and Table S13.

A ridge model trained to predict Chronos scores from *observed* transcriptional shifts (10-fold cross-validation over held-out knockdowns) reached a mean Pearson r of 0.57 across the three cancer lines. Applied unchanged to *predicted* shifts, it reached 0.50 (CASTER), 0.52 (Linear), 0.47 (PerturbMean) and −0.02 (ContextMean) (Figure 6C; Figure S14). Transferred responses therefore retain part of the essentiality information carried by measured responses. Because Chronos scores are largely shared between lines (Figure S13), this analysis speaks to general essentiality, not to line-specific dependency.

Predicted responses also preserved the functional organization of the knockdowns. We identified 15 co-regulated gene programs per line from the observed responses and annotated them with Reactome^58^. Predicted program activities correlated with observed ones at mean r (over four lines) = 0.63 (CASTER), 0.62 (Linear), 0.51 (PerturbMean) and 0.00 (ContextMean) (Figure 6D shows K562; Figure S15; Table S13). In a retrieval task (identifying which knockdown produced an observed response among all candidates in a line), the correct knockdown was within the top 10 for 56% (CASTER), 58% (Linear), 26% (PerturbMean) and 0% (ContextMean) of the 100 strongest knockdowns (Figure 6E). Response similarity between knockdowns recovered high-confidence STRING associations^59^ with an area under the receiver operating characteristic curve (AUROC) of 0.81 for observed responses, against 0.80 for CASTER and 0.75 for PerturbMean (Figure 6F).

## Discussion

This study asked which parts of a knockdown response carry over between cell lines and what a transfer model needs from a new line (Table S14). Metrics had to be calibrated before methods could be ranked, and even among calibrated metrics the order of close methods changed with the metric.

Perturbation biology increasingly relies on models that aim to predict an experiment before it is run^21,22,42^. We asked a focused version of this question with one CRISPRi library profiled in three cancer lines and a non-transformed line. Responses to the same knockdown were moderately correlated across lines, more so for strong responses and for the two *TP53* wild-type lines, HepG2 and RPE1, as Nadig et al. also found^43^.

When 30% of the target line was screened, Linear and CASTER performed alike and both beat the mean baselines. Linear was slightly ahead on top-100 Pearson delta with random splits, CASTER on weighted error metrics, and the two were level when whole complexes were held out. With only 2.5% screened, CASTER did better, mainly because it starts from the transferred mean. Because Linear is simpler and needs no control cells, a partial screen and a linear model is the practical choice for a new line. This agrees with evidence that well-specified linear models are strong competitors for perturbation prediction^38,39^ and with the warning that average-shift effects can dominate standard metrics^41^. Controls for a per-line offset, line identity and the control cells themselves, together with calibrated metrics, describe what the model adds beyond such effects. Benchmarks that render predictions on sampled control cells should remove that noise before comparing absolute accuracies. CASTER’s gains concentrated on strong responses, in particular knockdowns of ribosome genes, as did Linear’s. Because this gain was similar in the p53-deficient lines, it is not explained by p53 alone^54,56^.

In lines with no screening data, CASTER was worse than PerturbMean. This model, trained on three lines, did not find information in unperturbed transcriptomes that predicts how a new line reshapes a response; perturbation data from that line were what made adaptation work. Models trained on many more lines might learn such a mapping, but on current data a partial screen of each new line was more useful than control profiles alone. Across four lines, Linear’s calibrated score reached 91% of its 30% gain over PerturbMean when 20% of the library was screened (Figure 2C). CASTER led when about 34 knockdowns of the new line were screened; from 20% (about 274 knockdowns) on, Linear was slightly ahead with random splits. A ridge anchored on the transferred mean recovered most of this lead, and the mean-profile CASTER led as well, so the lead came mainly from the anchor, not from cell-to-cell variation among control cells.

We had expected transfer to be more accurate among cancer lines than to or from the non-transformed line. The opposite held, mostly because of the HepG2-RPE1 pair. Shared p53 status and adherent epithelial growth went with better transfer than transformation status did (HepG2 and RPE1 do not share a lineage). This suggests, as a hypothesis to test with more lines, that p53 status and growth mode may help when choosing reference lines for perturbation atlases.

Response size tracked Chronos scores in all three cancer lines, but the link was not line-specific: other lines’ scores predicted it about as well, consistent with earlier work in K562 cells^8^. Libraries designed around differential dependencies, such as synthetic-lethal partners of cancer drivers^60,61^, would be needed to test line-specific dependency.

Methodologically, these conclusions rest on selecting genes on control cells only, scoring every method with the same pipeline and a valid predicted-side DE test, and scoring against all available cells.

### Limitations of the study

This study has several limitations. First, it covers four lines, three of them cancer lines from different lineages; with so few lines, line generalization cannot be separated from lineage or p53 status, and conclusions about cancer in general require many more lines. Second, we trained on at most 32 cells per knockdown, which caps the achievable accuracy (noise ceilings), and used compact models trained on CPUs. Third, CASTER represents a perturbation by its observed effect elsewhere, so it cannot predict perturbations never observed in any line. Fourth, the dependency analyses rely on CRISPR knockout fitness, whereas the screens use CRISPRi knockdown, and the four screens differ in effector construct, time point and sequencing depth. Fifth, we did not compare with a large pretrained perturbation model. Finally, this work used public data only and did not test any prediction experimentally.

## Resource availability

### Lead contact

Requests for further information and resources should be directed to and will be fulfilled by the lead contact, Mohammad Ali Moni.

### Materials availability

This study did not generate new unique reagents.

### Data and code availability

- This paper analyzes existing, publicly available data. Accession numbers for the datasets are listed in the key resources table.
- All original code has been deposited at Zenodo and is publicly available at https://doi.org/10.5281/zenodo.24239384 as of the date of publication.
- Any additional information required to reanalyze the data reported in this paper is available from the lead contact upon request.

## Acknowledgments

We thank the authors of the Replogle et al. and Nadig et al. Perturb-seq studies, the Arc Institute for releasing the processed data, and the DepMap, STRING and Reactome teams. Icons in Figure 1A and the graphical abstract are from Servier Medical Art (smart.servier.com; CC BY 3.0) and from Marcel Tisch via Bioicons (CC0), recolored. This work was supported by the US National Science Foundation (award 2319522 to L.Z.).

## Author contributions

Conceptualization, methodology, software, formal analysis and writing – original draft, M.K. and S.A.D.; validation, S.A.D., B.K.P. and A.A.N.; writing – review and editing, all authors; supervision, L.Z. and M.A.M.

## Declaration of interests

The authors declare no competing interests.

## Declaration of generative AI and AI-assisted technologies in the writing process

During the preparation of this work the authors used Claude (Anthropic) to draft and edit text. After using this tool, the authors reviewed and edited the content as needed and take full responsibility for the content of the publication. Claude was also used in the analysis, as described in STAR Methods.

## Supplemental information

Document S1. Methods S1, Note S1, Figures S1–S17 and Tables S1–S20.

Data S1. Tables S1 to S20 in Excel format, including the per-seed learning-curve values (Table S5, part B), related to STAR Methods.

## STAR★Methods

### Key resources table

| REAGENT or RESOURCE | SOURCE | IDENTIFIER |
|---|---|---|
| Deposited data |  |  |
| K562 and RPE1 CRISPRi Perturb-seq | Replogle et al.<sup>8</sup> | Figshare+: https://doi.org/10.25452/figshare.plus.20029387 |
| HepG2 and Jurkat CRISPRi Perturb-seq | Nadig et al.<sup>43</sup> | GEO: GSE264667 |
| Knockdown-filtered four-line release (file replogle_concat.h5ad) | Adduri et al.<sup>37</sup> | https://huggingface.co/datasets/arcinstitute/State-Replogle-Filtered (downloaded October 2026) |
| DepMap 24Q4 Public, CRISPR gene effect (Chronos; file CRISPRGeneEffect.csv) | DepMap<sup>62</sup> | Figshare+: https://doi.org/10.25452/figshare.plus.27993248 |
| DepMap 24Q4 mutation calls (*TP53* status) | DepMap<sup>62</sup> | Figshare+: https://doi.org/10.25452/figshare.plus.27993248 |
| STRING v12.0 human protein links (file 9606.protein.links.v12.0) | Szklarczyk et al.<sup>59</sup> | https://string-db.org; RRID:SCR_005223 |
| Reactome pathways (GMT file) | Milacic et al.<sup>58</sup> | https://reactome.org/download-data; RRID:SCR_003485 |
| Consensus p53 target genes | Fischer<sup>48</sup> | N/A |
| Cell-cycle signature genes | Tirosh et al.<sup>53</sup> | N/A |
| Experimental models: Cell lines |  |  |
| K562 | Lozzio and Lozzio<sup>63</sup> | RRID:CVCL_0004 |
| HepG2 | Aden et al.<sup>64</sup> | RRID:CVCL_0027 |
| Jurkat | Schneider et al.<sup>65</sup> | RRID:CVCL_0065 |
| hTERT RPE-1 | Bodnar et al.<sup>47</sup> | RRID:CVCL_4388 |
| Software and algorithms |  |  |
| CASTER and all analysis code | This paper | Zenodo: https://doi.org/10.5281/zenodo.24239384 |
| Python 3.12.13 | Python Software Foundation | https://www.python.org; RRID:SCR_008394 |
| PyTorch 2.13.0 (CPU build) | Paszke et al.<sup>66</sup> | https://pytorch.org; RRID:SCR_018536 |
| NumPy 2.5.3 | Harris et al.<sup>67</sup> | https://numpy.org; RRID:SCR_008633 |
| SciPy 1.18.1 | Virtanen et al.<sup>68</sup> | https://scipy.org; RRID:SCR_008058 |
| scikit-learn 1.9.1 | Pedregosa et al.<sup>69</sup> | https://scikit-learn.org; RRID:SCR_002577 |
| pandas 3.0.6 | McKinney<sup>70</sup> | https://pandas.pydata.org; RRID:SCR_018214 |
| h5py 3.16.0 | Collette<sup>71</sup> | https://www.h5py.org |
| NetworkX 3.7 (Louvain communities) | Hagberg et al.<sup>72</sup> | https://networkx.org; RRID:SCR_016864 |
| Matplotlib 3.11.2 | Hunter<sup>73</sup> | https://matplotlib.org; RRID:SCR_008624 |

### Experimental model and study participant details

This study generated no new experimental data. All analyses use published CRISPR interference (CRISPRi) Perturb-seq profiles of four human cell lines: K562 (chronic myeloid leukemia [CML], *BCR-ABL1*-positive; female)^63^, HepG2 (hepatoblastoma-derived, historically described as hepatocellular carcinoma; male)^64,74^, Jurkat (T-cell acute lymphoblastic leukemia [T-ALL]; male)^65,75^ and hTERT RPE-1 (RPE1; telomerase-immortalized, non-transformed retinal pigment epithelium; female)^47^. No cell lines were cultured in this study; authentication was done by the original studies. K562 and RPE1 were profiled by Replogle et al.^8^ and HepG2 and Jurkat by Nadig et al.^43^ with the same essential-gene CRISPRi single-guide RNA (sgRNA) library. The four screens differ in CRISPRi effector construct, time point after transduction and sequencing depth, which adds technical differences between lines to the biological ones.

### Method details

#### Data acquisition

We used the four-line release of these data published with the State model^37^ (key resources table). It contains library-size-normalized, log(1 + x)-transformed expression of 6,546 genes measured in all four lines. Knockdowns and cells were already filtered for on-target knockdown, as described by Adduri et al.^37^: median residual target expression of the knockdown ≤ 0.30, residual of the cell ≤ 0.50, and ≥ 30 cells. We used this release rather than the per-line original files for uniform processing. Models were trained on a random subsample of up to 32 cells per knockdown and line and 3,000 control cells per line, drawn from the cell labels only, without use of expression values. For scoring against all cells, every remaining knockdown cell of the release was added (all-cells data; download steps are given with the code).

#### Gene space

We modeled 2,000 genes. They were chosen as the top genes by bin-normalized dispersion (20 mean bins; our implementation of the procedure of Satija et al.^76^) computed on the **pooled control cells only**. Of the 2,023 knockdown targets, 1,850 are among the 6,546 measured genes and were excluded before selection; the other 173 are not measured. Because no knockdown cell enters gene selection, held-out effects cannot leak through the choice of genes, and the on-target signal cannot inflate any metric.

#### Problem formulation

Let *χ*_*ℓ*_^0^ be the distribution of unperturbed cells in context (cell line) *ℓ*, and *χ*_*ℓ*_^*p*^ the distribution after knockdown *p*. With *G* the number of modeled genes, the pseudobulk shift of *p* in *ℓ* is the vector in ℝ^*G*^

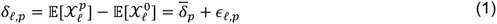

Here *δ̄*_*p*_ is the mean of *δ* over the lines that observed *p*, and *∈*_*ℓ*,*p*_ is what is left for line *ℓ*. Transfer to a target line *t* asks for *χ*_*t*_^*p*^, given control cells of *t*, observations of *p* in other lines and, optionally, a set of other knockdowns screened in *t*. PerturbMean estimates only *δ̄*_*p*_; any method that beats it must recover part of *∈*_*t*,*p*_ (Methods S1, Proposition 4).

#### CASTER architecture

CASTER maps a set of *S* control cells *X*^0^ ∈ ℝ^*S*×*G*^ from the target line, together with a knockdown, to a set of *S* predicted perturbed cells. It has four parts.

##### Transfer prior

The knockdown is represented by its observed shift in the *other* lines. For knockdown *p* in line *ℓ*, with *C*_*p*_ the lines that observed *p*:

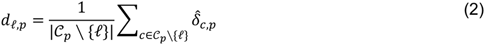

Here *δ̂*_*c*,*p*_ is the empirical shift relative to the control mean of line *c*. The prior *d* equals the PerturbMean prediction (Equation 6). The target line never contributes to any prior. When no other source line observed *p* (always the case in single-source transfer), the prior is computed from a random half of *p*’s cells, and the model is trained against the disjoint other half.

##### Encoders

A 4-layer multilayer perceptron (MLP) with Gaussian error linear units (GELU)^77^, *f*_cell_, embeds each control cell into ℝ^ℎ^ (ℎ = 256). A 2-layer MLP *f*_prior_ embeds the prior. A line summary *c* is the mean of the cell embeddings, passed through a 2-layer MLP that outputs feature-wise linear modulation (FiLM)^51^ parameters (*γ*, *β*); ⊙ is the element-wise product:

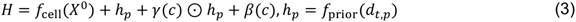

##### Set transformer

*H* passes through a 4-layer, 8-head, pre-norm, bidirectional transformer encoder *f*_ST_^78,79^ without positional encodings. The model is therefore permutation-equivariant over cells^80^, which we checked numerically (Methods S1, Proposition 2).

##### Output

The prediction adds a line-dependent, gene-wise rescaling of the transferred shift and a cell-resolved correction (**1** is the vector of *S* ones):

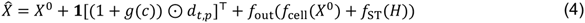

Here *g* is linear in *c*, and *f*_out_ is a LayerNorm^81^ followed by a linear map. Both *g* and the output layer of *f*_out_ are initialized at zero, so training starts exactly at the PerturbMean solution (Equation 4; Methods S1, Proposition 1).

##### Ablations

On the K562 partial-screen split, we compared the full model (*S* = 32) with set sizes *S* = 1 (single cells) and *S* = 8, and with versions without the transformer (a per-cell MLP), without FiLM, without the prior (a learned one-hot knockdown embedding instead) and with a one-hot knockdown embedding added to the prior, each trained for 20 epochs with one or two seeds (two for the main variants). Each variant was compared with the full model by paired Wilcoxon signed-rank tests over held-out knockdowns on Pearson delta, discrimination and DE AUPRC, with Holm correction over the 18 tests (six variants × three metrics).

#### Training objective

For a predicted set *X̂* with cells *x̂*_*i*_ and an observed set *X* with cells *x_j_* of the same knockdown and line, the loss is the squared maximum mean discrepancy (MMD)^49^ with the energy-distance kernel^50,82^, plus a pseudobulk term on the set means *x̂̄* and *x̂*:

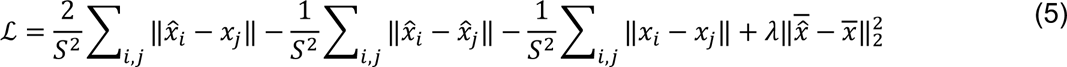

We used *λ* = 1 and *S* = 32. Training used AdamW^83^ (learning rate 3 × 10⁻⁴, weight decay 10⁻⁴, batch 32 sets), gradient-norm clipping at 1.0, a 5% linear warm-up and cosine decay. The number of epochs was 20 for the partial-screen, unseen-line and learning-curve tasks and the ablations, and 10 for single-source transfer. After each epoch the validation loss was computed, and the checkpoint with the lowest validation loss was kept. In the partial-screen task, validation uses the 5% validation knockdowns of the target line. In the unseen-line and transfer tasks, it uses 5% of source-line items, so no perturbed cell of the target line is ever used. All runs used CPUs only.

#### Baselines

Every baseline predicts one mean shift *δ̂* per knockdown. So that single-cell DE metrics are computed identically for all methods, this shift is added to the same control cells given to CASTER (rendering).

- **NoChange**: a prediction of zero shift, *δ̂* = 0.
- **PerturbMean**: *δ̂*_*t*,*p*_ = *d*_*t*,*p*_, the mean shift of *p* over the source lines that observed it.
- **ContextMean**: the mean shift of the target line’s screened knockdowns (training plus validation), the same for every knockdown^39^. It equals the perturbed mean (below).
- **LineOffset**: PerturbMean plus one constant shift per target line, the mean residual 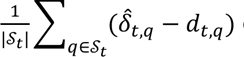 over the screened knockdowns *S*_*t*_. It tests whether a model’s gain is a single per-line offset.
- **Linear** (a linear transfer model): ridge regression^84^ (penalty chosen by efficient leave-one-out cross-validation over 13 values) from a 64-component principal component analysis (PCA) of the concatenated source shifts of *p* to its target shift (Equation 8). It is fitted on the target line’s screened knockdowns; a missing source is imputed with the mean over the observing sources. This linear model adapts to the target line and is of the kind shown to match deep models^38,39^.

#### Small-screen controls

At 2.5%, 5% and 10% screened, four further models were fitted on the same nested splits as the learning curve and scored in the same way. (i) Anchored ridge: PerturbMean plus a ridge regression of the residual (target shift minus PerturbMean) on the same PCA features as Linear (with an intercept; *min*(64, |*S*_*t*_| − 1) components, so 33 at 2.5%; PCA fitted on the source-line features of all knockdowns of the split, which contain no target data; penalty chosen per split by efficient leave-one-out cross-validation from 13 values log-spaced in [10^-2^, 10^4^]). (ii) Scaled PerturbMean: PerturbMean with a gene-wise scale *b*_g_ fitted on the screened knockdowns by least squares through the origin and shrunk toward 1, 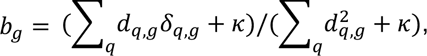 with *κ* the median of 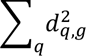 over genes. (iii, iv) Linear with 8 or 16 PCA components instead of 64. In addition, the mean-profile CASTER (below) was trained at 2.5% (three seeds).

#### Tasks and splits

##### Partial-screen task

For each target line *t*, the eligible knockdowns are those observed in *t* and in at least one other line. They were shuffled (one ordering per seed) and split into 25% training, 5% validation and 70% test. All knockdowns of the three other lines are training data. Table S19 lists the information available to each method.

##### Learning curve

With the same ordering, the test set was kept fixed (the last 70%) and the screened set was the first 2.5%, 5%, 10%, 20% or 30% of the ordering, split 5:1 into training and validation, so screened sets are nested; at 30% this is the partial-screen split. Baselines and CASTER were run for three seeds at every fraction (scores and screened-set sizes in Table S5; sizes at 30% in Table S20).

##### Unseen-line task

CASTER is trained on all knockdowns of the other three lines, with 5% of their items for validation, and tested on every eligible knockdown of the target line; no perturbed cell of the target line is used. The task was run with one seed per line.

##### Single-source transfer

For each ordered pair (source *s*, target *t*), training uses *s* only and the test set is every knockdown observed in both (one seed). Unperturbed cells of every line, including *t*, are available to all methods, as in Adduri et al.^37^. Control cells of each line were split 50/50 into model-input control cells and disjoint reference control cells for DE tests.

##### Seeds

The partial-screen task was run with three seeds (numbered 0, 1 and 2); each seed changes the knockdown split, the model initialization and the minibatch order. "First seed" refers to seed 0.

##### Controls for per-line effects

(i) LineOffset (above). (ii) Mean-profile CASTER: identical architecture and training, but every control set is replaced by the line’s mean control profile repeated *S* times, so the model knows which line it is in but sees no cell-to-cell variation. (iii) Wrong-line control cells: the trained CASTER is given the control cells of another line (the next line in the order K562, HepG2, Jurkat, RPE1); its predicted shift (prediction minus its input) is added to the target line’s own control cells, so only the model’s use of the control set changes. (iv) Line-label CASTER: every control set is replaced by the line’s mean control profile with its genes shuffled by a fixed, line-specific permutation, which identifies the line on the scale of real profiles but carries no gene-wise information about it. Because the outputs of the mean-profile and line-label models are in expression units, their predicted response is the output minus the line’s mean control profile; for the mean-profile model this equals the output minus its input.

##### Complex-grouped splits

Knockdowns were grouped by Louvain communities^85^ (NetworkX^72^; resolution 1, seed 0) of the STRING (v12)^59^ graph with combined score ≥ 950 among the 2,023 targets (500 groups, the largest 157 genes). All 73 ribosomal-protein targets fall in one 131-gene group, together with 58 translation-initiation, signal-recognition and ribosome-biogenesis factors. Whole groups were assigned to training, validation or test (three seeds), so no held-out gene had a same-group sibling in training.

#### Gene programs and cell-cycle analyses

Gene programs were defined per line by K-means^86^ (scikit-learn; 15 programs, 10 restarts) on the z-scored observed shift matrix (knockdowns × genes), clustering genes by their response profiles. Programs were annotated with their most enriched Reactome pathway^58^ (hypergeometric test, background = modeled genes). Program activity is the mean shift over program genes. Knockdown-similarity structure used average-linkage hierarchical clustering of cosine distances between shifts of knockdowns with ≥ 10 observed DEGs, compared with the adjusted Rand index^87^. Recovery of functional relationships used STRING (v12)^59^ pairs with combined score ≥ 700, scored by the cosine similarity of the shifts (AUROC). The cell-cycle shift of a knockdown is the mean shift over the S-phase and G2/M signature genes of Tirosh et al.^53^ present in the gene space; the cell-cycle error (Figure 5D) is the mean absolute difference between predicted and observed cell-cycle shifts over held-out knockdowns.

#### p53 activity

p53 activity was scored with 14 consensus p53 target genes^48^ measured in all four lines (*MDM2*, *BAX*, *TP53I3*, *RRM2B*, *ZMAT3*, *FDXR*, *AEN*, *TRIAP1*, *XPC*, *RPS27L*, *CCNG1*, *GADD45A*, *PPM1D* and *APAF1*; *CDKN1A* and *GDF15* are not among the shared measured genes). Each gene was z-scored against the line’s control cells and the score was the mean z over the 14 genes. A knockdown was called p53-inducing when its cells’ score exceeded the controls’ (Welch t test, Benjamini-Hochberg [BH] FDR < 0.05, positive shift). *TP53* status: K562 *TP53* null and Jurkat *TP53* mutant (DepMap damaging-mutation calls), HepG2 *TP53* wild type (DepMap) and RPE1 *TP53* wild type^8,47^. The rank correlation of p53 scores between two lines (Figure S2B) uses the knockdowns observed in both.

#### Simulation (positive control)

Four synthetic lines shared 160 knockdowns and 400 genes. Knockdown effects were sparse combinations of 12 gene programs (program loadings ∼ N(0, 0.5²) with 30% nonzero; program weights ∼ N(0, 1) with 15% nonzero). Each line observed a random 80% of the knockdowns, with 24 cells per knockdown and 600 controls drawn from N(baseline + effect, 0.4²), clipped at 0. In the “unlearnable” scenario each line added independent noise to the loadings (N(0, 0.25²) on 20% of entries). In the “learnable” scenario a gene’s effect in a line was the shared effect scaled by its baseline expression in that line (baselines ∼ Gamma(1.2, 0.9) with 25% of genes silent), which is observable from control cells. All methods were run on the HepG2-like line with the partial-screen split (25% plus 5% screened) and 60 epochs.

#### Dependency analyses

Chronos gene effects (Chronos scores)^46^ (DepMap^45,57,62^, release 24Q4) were taken for K562 (ACH-000551), HepG2 (ACH-000739) and Jurkat (ACH-000995); RPE1 is not profiled. Response size is *m* = log(1 + *n*_DEG_), with *n*_DEG_ the number of DEGs. The main correlation (Figure 6A) uses all knockdowns of a line with a Chronos score; the decile means in Figure 6A are the mean response size within deciles of the Chronos score. The line-specific response of a knockdown in line *ℓ* is its response size minus the mean over the other two cancer lines, and the line-specific dependency is defined the same way on Chronos scores (Equation 11). For dependency prediction, a ridge model was trained, within 10-fold cross-validation over the held-out knockdowns of each cancer line, to map *observed* shifts to Chronos scores. It was then applied unchanged to the shifts predicted by each method for the held-out fold. As a control for line specificity, each line’s response size was also correlated with the other lines’ Chronos scores, and each line’s Chronos scores were predicted by the mean Chronos score of the other two lines.

#### Formal definition of baseline models

Let 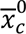 be the mean of the model-input control cells of line *c*, and 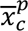 the mean of the observed cells of knockdown *p* in *c*. The observed shift is 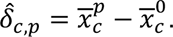 For the target line *t*, let *S*_*t*_ be its screened knockdowns (training plus validation) and *C*_*p*_ the source lines that observed *p*.

##### PerturbMean

averages the shift of *p* over the source lines that observed it:

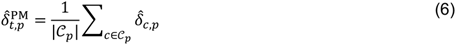

##### ContextMean

predicts the same shift for every knockdown of *t*, the mean shift of its screened knockdowns (the perturbed mean):

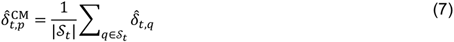

##### Linear

Each knockdown is described by the concatenation of its shifts in the *K* source lines, *u*_*p*_ = [*δ̂*_*c*_1_,*p*_; … ; *δ̂*_*cK*,*p*_] ∈ ℝ^*KG*^. A source that did not observe *p* contributes 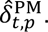 The features are reduced by PCA, fitted on source data only, to *k* = *min*(64, |*S*_*t*_| − 1) components *v*_*p*_ = *W*^*T*^(*u*_*p*_ − *u*), where *W* holds the component loadings and *u* is the feature mean. A ridge map *B* ∈ ℝ^*k*×*G*^ with intercept *b* is then fitted on the screened knockdowns:

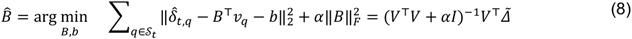

Here *V* stacks the centered *v*_*q*_ and Δ̃ the centered target shifts. The penalty *α* is chosen from 13 log-spaced values in [10^-2^, 10^4^] by efficient leave-one-out cross-validation. The prediction is 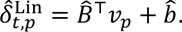 NoChange predicts *δ̂* = 0. Every pseudobulk prediction is rendered as cells by adding it to the same control cells, 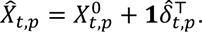

#### Formal definition of downstream analyses

##### Knockdown retrieval

For the 100 knockdowns *j* with the most observed DEGs in a line, the number of predictions that are closer to the observed response than the correct one (rank 0 is best) is

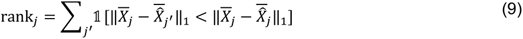

and the top-*k* accuracy is

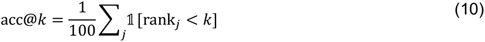

Here the observed response is compared with all predictions; in Equation 13 the prediction is compared with all observed responses.

##### STRING-pair recovery

For knockdowns with ≥ 10 observed DEGs, each unordered pair (*p*, *q*) is labeled positive if STRING v12 reports a combined score ≥ 700 between their products. It is scored by the cosine similarity of the predicted shifts, *s*_*pq*_ = cos(*δ̂*_*p*_, *δ̂*_*q*_), and summarized by the AUROC.

##### Gene programs

Let *Z* be the observed shift matrix (knockdowns × genes), z-scored per gene. K-means with 15 clusters (10 restarts, seed 0) on the rows of *Z*^*T*^ assigns each gene to a program *k* with gene set *G*_*k*_. The activity of program *k* under knockdown *p* is 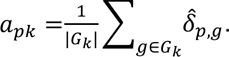 Concordance is the Pearson correlation across knockdowns of predicted and observed *a*_·*k*_. Programs are annotated by the Reactome pathway with the smallest hypergeometric p value, with the modeled genes as the background and at least 3 overlapping genes.

##### Knockdown-similarity structure

Cosine-distance matrices of predicted and observed shifts (knockdowns with ≥ 10 DEGs) were clustered by average linkage into 2 to 20 clusters, and the clusterings were compared by the adjusted Rand index.

##### Dependency prediction

Within each cancer line, held-out knockdowns with a measured Chronos score were split into 10 folds. A ridge model *f* (penalty chosen by leave-one-out cross-validation over 13 values in [10^-1^, 10^T^]) was fitted on the *observed* shifts of 9 folds and applied to the observed and to each method’s predicted shifts of the remaining fold. The score is the Pearson correlation between the out-of-fold predictions *f*(*δ̂*_*p*_) and the measured Chronos scores. Because the regressor never sees a predicted shift during training, any loss of accuracy reflects information missing from the prediction.

##### Line-specific dependency

For gene *p* observed in all three cancer lines, with response size 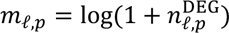 and Chronos score *e*_*ℓ*,*p*_, the line-specific components are

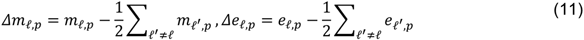

The line-specific test is the Spearman correlation of *Δm*_*ℓ*,·_ and *Δe*_*ℓ*,·_ per line.

##### Response similarity between lines

For each pair of lines and each knockdown with ≥ 10 DEGs in both, similarity is the Pearson correlation of the two observed shift vectors over the modeled genes. DEG-set similarity between lines is the Jaccard index of the two DEG sets. The stratified analysis repeats this for minimum response sizes of 10, 25, 50, 100 and 200 DEGs.

#### Software and computation

All analyses were run on a single workstation (Windows 10, 4 CPU cores, 8 GB RAM, no GPU) with Python 3.12.13, PyTorch 2.13.0 (CPU build)^66^, NumPy (v2.5.3)^67^, SciPy (v1.18.1)^68^, scikit-learn (v1.9.1)^69^, pandas (v3.0.6)^70^, h5py (v3.16.0)^71^, NetworkX (v3.7)^72^ and Matplotlib (v3.11.2)^73^. Partial-screen and unseen-line runs took 19 to 62 min each to train and evaluate (Table S20); single-source transfer models about 10 min each. All randomness (cell subsampling, control-pool splits, knockdown splits, model initialization, minibatch sampling and the control cells given to each method) is seeded. Control input sets are derived deterministically from a hash of the knockdown, seed and line, so that every method receives identical inputs. The analysis runs as a fixed sequence of scripts. The tables behind every figure panel are provided with the code.

The code includes tests for the multiple-testing correction, the DE test, the discrimination metric, the loss, permutation equivariance and the separation of splits and control pools. The code checks these rules at run time.

#### Use of generative AI

Claude (Anthropic) wrote the analysis and plotting code, ran the computational experiments and drafted text under the authors’ direction. All figures are plots made by this code; icons are from the credited sources, and no image was made by a generative model. The authors chose the analyses, reviewed the code and its outputs, and checked the reported numbers against the stored results.

### Quantification and statistical analysis

#### Evaluation suite

For test knockdown *j* with observed cells *X*_j_, predicted cells *X̂*_j_ (same number of cells) and reference control cells *R* (500 cells, disjoint from model inputs); bars denote means over cells:

##### Pearson delta

The correlation between predicted and observed pseudobulk shifts:

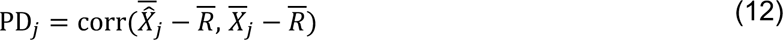

##### Perturbation discrimination

Following Wu et al.^40^ and Adduri et al.^37^, with *r_j_* the number of other test knockdowns whose observed pseudobulk is closer in L1 distance to the prediction than the true one, and *T* the number of test knockdowns:

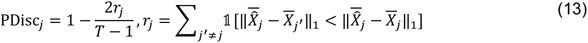

A perfect model scores 1 and chance scores 0. **Observed differential expression (ground truth).** Observed cells of each knockdown are tested against *R* gene by gene with the Wilcoxon rank-sum test^88,89^ (asymptotic, tie-corrected), with BH correction^90^ at FDR < 0.05. Observed log₂ fold changes are log₂ ratios of mean expression back-transformed to counts, with pseudocount 0.01. The limitations of cell-level DE tests are well known^91,92^.

##### Predicted differential expression

A Wilcoxon test is not valid for predicted cells that are made by adding a shift to control cells. Adding a shift to a control cell turns each of its exact zeros into a small positive value, and against zero-inflated reference cells the rank test then calls most genes significant whatever the shift. In the HepG2 run, every shifted method produced 1,283 to 1,494 “DEGs” per knockdown, against 0.1 for unshifted control cells. We therefore score predictions with a moment-matched z test (Figure S16) on the predicted pseudobulk shift *δ̂*_g_, using the reference-control standard deviation *s*_g_ and the sample sizes of the observed experiment; *Φ* is the standard normal distribution function:

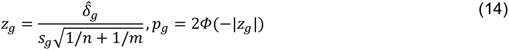

Here *n* is the number of perturbed cells and *m* = 500, with BH adjustment. Predicted fold-change ranks and signs use *δ̂*_g_. Observed ground truth is unchanged. Wilcoxon-based results are given in Table S17 for comparison. **DE AUPRC** is the area under the precision-recall curve (average precision^93,94^) for recovering observed DEGs from predicted −log₁₀ adjusted p values. **DE overlap** (predicted versus observed DEGs) is

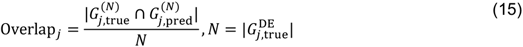

Here both sets are ranked by the absolute log₂ fold change among significant genes. **Fold-change Spearman** is computed over observed DEGs. **Direction agreement** is the fraction of observed DEGs with the correct predicted sign. **Effect-size Spearman** is the rank correlation, across knockdowns within a run, between the predicted and observed numbers of DEGs.

**Mean absolute error (MAE) and mean squared error (MSE)** are the mean absolute and mean squared differences between predicted and observed pseudobulk shifts over the *G* = 2,000 modeled genes, 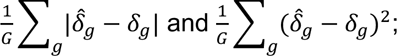 lower is better, and paired tests are oriented accordingly.

##### Calibrated metrics

Following Miller et al.^44^ and Mejia et al.^52^, each held-out knockdown’s cells were split into two random halves (seeded per knockdown). Half A defines the truth; half B, with shift *δ*^*B*^, is the technical duplicate (positive control). A Welch t test of half A against all other cells of the line gives per-gene statistics *t*_g_ and BH-adjusted p values *q*_g_. DEG weights are *w_g_* = ((|*t*_g_| − *min*|*t*|)/(*max*|*t*| − *min*|*t*|))^2^, the interpolated duplicate is 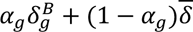 with *α* = 1 − *q*_g_, and *δ* is the perturbed mean: the mean shift of the line’s screened knockdowns, which is also the ContextMean prediction. Metrics: Pearson delta; top-100 Pearson delta (Pearson delta relative to the perturbed mean on the 100 genes with the largest absolute *t*_g_); MSE; MAE; weighted MSE (WMSE), 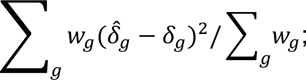 weighted *R*^2^ relative to control and to the perturbed mean (clipped at −1); the normalized inverse rank (NIR; the fraction of other held-out knockdowns whose truth is farther, in Euclidean distance, from the prediction than the correct truth) and the perturbation discrimination score (PDS; the same with Manhattan distance), with the names used by Miller et al.^44^. The negative control is the perturbed mean, or a zero shift (NoChange) for the two metrics relative to the perturbed mean. The dynamic range fraction (DRF) of a metric is (technical duplicate − negative control) / (perfect − negative control), where the perfect score is 1 for correlations, *R*^2^, NIR and PDS, or (negative control − technical duplicate) / negative control for lower-is-better metrics; it is computed per knockdown, clipped to [−1, 1] and averaged. The ratio-of-means version is also reported (Table S9). A method’s scaled score is (method − negative control) / (technical duplicate − negative control), computed on the per-line, per-seed means and then averaged. A metric is called calibrated when its DRF is 0.2 or more; scaled scores are shown only for those. Unless stated otherwise, scores use noise-free predictions; the 32-cell scores as rendered are given for comparison.

##### Noise-free rescoring

Every method’s prediction is rendered on *n* control cells sampled from the model-input control cells (*n* = number of observed cells of the knockdown, at most 32), so its pseudobulk shift contains the sampling noise 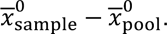 We subtracted this term from every method’s predicted shift (for CASTER, this treats its output relative to its own input as the predicted effect) and recomputed all pseudobulk metrics. These noise-free scores are reported alongside the rendered ones.

##### Noise ceiling

The Pearson correlation *r*_ℎ_ between the shifts of the two halves estimates the split-half reliability; by Spearman-Brown, the reliability of the full-cell shift is *ρ* = 2*r*_ℎ_/(1 + *r*_ℎ_), and the highest Pearson delta any predictor can reach against the full-cell truth is 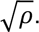 The mean noise ceiling of a line is the mean over its held-out knockdowns.

##### All-cells rescoring

The trained models were not retrained. For each held-out knockdown, the truth shift, the observed DEGs (Wilcoxon against the same 500 reference controls) and the split-half calibration were recomputed from all cells of that knockdown in the release (median 57 cells per held-out knockdown), and each method’s noise-free predicted shift was scored against them; predicted-side DE tests used the knockdown’s full cell count. The reference-control mean was checked to equal the one the predictions were stored against.

##### Noise-corrected similarity between lines

For each knockdown with at least 10 DEGs in two lines, the raw similarity *r* is the Pearson correlation of its two mean shifts. Its split-half reliability in each line is the mean, over 20 random splits of its cells into halves, of the correlation between the half shifts, stepped up with Spearman-Brown to the full sample (*ρ*_*a*_, *ρ*_*b*_). The corrected similarity is 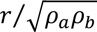 (correction for attenuation^95^). We report medians over knockdowns with bootstrap intervals (1,000 resamples of knockdowns) and a sensitivity analysis restricted to knockdowns with reliability ≥ 0.5 in both lines. Similarity on all 6,546 measured genes (Table S7) repeats this analysis without restricting to the modeled genes.

##### Comparison of CASTER and Linear

For each line, the per-knockdown difference (CASTER minus Linear) in the scaled top-100 Pearson delta against all cells was averaged over seeds and the mean over knockdowns was bootstrapped over knockdowns (2,000 resamples, percentile intervals). We report 95% intervals and, as a practical margin, 5% of the line’s dynamic range (technical duplicate minus zero shift).

##### Learning-curve scoring

Each split of the learning curve was scored against all cells of its test knockdowns on the scaled top-100 Pearson delta. The two halves of each knockdown’s cells, the Welch statistics that define its top 100 genes and the technical duplicate do not depend on the screened fraction, so they were computed once per knockdown and reused; the perturbed mean was recomputed from each split’s screened knockdowns.

##### Grouped versus random splits

The two splits were compared on the scaled top-100 Pearson delta against all cells, on the full test sets and on the knockdowns held out under both splits (same line and seed). Because the perturbed mean depends on the screened set, the raw score is reported too. The change in CASTER minus Linear (grouped minus random, matched knockdowns) was bootstrapped over knockdowns within each line-seed run (2,000 resamples, percentile intervals; Table S12).

##### Response size and gain

For the gain over PerturbMean (Figure S8), held-out knockdowns were binned by their number of observed DEGs (< 10, 10 to 49, ≥ 50), and the gain was correlated with log(1 + *n*_DEG_) by Spearman correlation per line.

##### p53 analyses

The data-defined p53 module (101 genes) contains the 14 targets and every gene whose shift correlates with the p53 score across knockdowns at r > 0.3 in both HepG2 and RPE1. Knockdowns inducing p53 in both lines were compared with the others by a two-sided Mann-Whitney test on HepG2-RPE1 similarity, overall and within quartiles of response size. For the size-matched comparison, each knockdown inducing p53 in both lines was paired one to one (without replacement) with the other knockdown of nearest response size (log(1 + the smaller DEG count of the two lines); 200 pairs) and the pairs were compared by a two-sided Wilcoxon signed-rank test. The regression is an ordinary least-squares regression of HepG2-RPE1 similarity on a p53 indicator and response size (two-sided t test on the coefficient).

##### Ribosome analysis

The ribosome gene set comprises ribosomal proteins (gene symbols starting with RPL or RPS, excluding *RPS6K* and *RPL3L* genes) and ribosome-biogenesis genes (Reactome pathways “Major pathway of rRNA processing in the nucleolus and cytosol”, R-HSA-6791226, and “rRNA processing in the nucleus and cytosol”, R-HSA-8868773). Per line and seed, the gain over PerturbMean was regressed by ordinary least squares on a ribosome indicator and log(1 + *n*_DEG_) (two-sided t test on the ribosome coefficient).

#### Statistics

In the partial-screen task, CASTER was compared with each baseline by one-sided paired Wilcoxon signed-rank tests (CASTER better) over held-out knockdowns pooled across the four lines (first seed; n = number of test knockdowns), with Holm correction^96^ over nine tests (three baselines × three main metrics). In the unseen-line task, CASTER and PerturbMean were compared by two-sided paired Wilcoxon signed-rank tests (Holm over three metrics). Transfer among cancer lines was compared with transfer involving RPE1 by a two-sided paired Wilcoxon test on per-knockdown accuracy. Response size and Chronos scores were related by Spearman correlations with Holm correction over the three cancer lines. Lower-is-better metrics (MAE, MSE) were tested in that direction. In Table S17, each Holm family is one task, comparison and metric group, as given in its caption.

Other tests: Mann-Whitney tests (p53 similarity; n = knockdowns), Wilcoxon signed-rank tests on matched pairs (p53; n = pairs), Welch t tests with BH correction (p53 calls and calibrated metrics; n = cells), Wilcoxon rank-sum tests with BH correction (observed DEGs; n = cells), hypergeometric tests (program annotation; n = genes), ordinary least-squares regressions (p53 and ribosome terms; n = knockdowns) and percentile bootstrap intervals (1,000 resamples for similarity medians, 2,000 for the CASTER-Linear comparisons). Tests over knockdowns and seeds pool the same knockdown from up to three seeds; these p values are therefore approximate.

Analyses beyond the main comparisons are exploratory and reported without multiplicity claims: the offset, identity and line-label controls, calibrated metrics, all-cells rescoring, the learning curve, the small-screen controls, noise-corrected similarity, the p53 analyses, complex-grouped splits and the grouped-versus-random comparison on matched knockdowns. Counts of line-seed runs (for example 11 of 12) treat seeds as separate runs; where the line is the unit, we say so. Statistical details of each analysis, including the test used, the value of n and what n represents (knockdowns, lines or seeds), and the meaning of centers and error bars, are given in the Results, the figure legends and Tables S17 and S9. Tests were two-sided unless stated. p or adjusted p below 0.05 was called significant. p values below 10⁻³⁰⁰ are below machine precision and are reported as p < 10⁻³⁰⁰. No data were excluded beyond the filters described above.

## Methods S1. Formal properties of CASTER and of context transfer, related to STAR Methods

We state four properties that frame the benchmark results. Notation follows the problem formulation in STAR Methods; *d* = *d*_*t*,*p*_ is the transfer prior of knockdown *p* in target line *t*.

### Proposition 1 (CASTER contains PerturbMean)

At initialization (*g* ≡ 0, and the final linear layer of *f*_out_ equal to zero), for any control set *X*^0^ and knockdown *p*, the CASTER prediction is

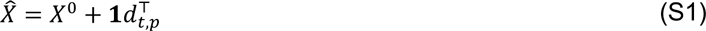

Its pseudobulk shift therefore equals the PerturbMean prediction exactly. *Proof.* The output is *X*^0^ + **1**[(1 + *g*(*c*)) ⊙ *d*]^*T*^ + *f*_out_(·). With *g* = 0 the middle term is **1***d*^*T*^, and *f*_out_(*z*) = *Wz* + *b* with *W* = 0 and *b* = 0 is identically zero. ∎ Checkpoints are compared after each training epoch, so this property describes the starting point of training; it does not bound the validation loss of the selected model.

### Proposition 2 (permutation equivariance)

For any permutation matrix *Π* ∈ {0,1}^*S*×*S*^, CASTER(*ΠX*^0^, *p*) = *Π*CASTER(*X*^0^, *p*). *Proof. f*_cell_ acts row-wise and commutes with *Π*. The line summary *c* is a mean over rows and is invariant. The FiLM and prior terms are broadcast identically to every row. Self-attention without positional encodings is permutation-equivariant^S1,S2^. *f*_out_ and the residual act row-wise. A composition of equivariant and row-broadcast invariant maps is equivariant. ∎ We checked this property numerically. It means that predictions depend on the *distribution* of control cells, not on their order.

### Proposition 3 (the training loss identifies distributions)

For distributions *P*, *Q* on ℝ^*G*^ with finite first moments, the energy distance^S3^

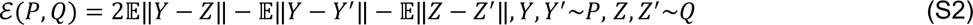

is nonnegative and equals zero if and only if *P* = *Q*. It is the squared MMD with the (conditionally negative definite) energy kernel^S4,S5^. The empirical V-statistic in the loss converges almost surely to ℰ as *S* → ∞. Minimizing the population loss therefore matches the full predicted and observed perturbed distributions, not only their means. The added pseudobulk term is minimized by the conditional mean, so the combined objective is minimized by any predictor whose distribution equals the observed one, and both terms vanish there. ∎

### Proposition 4 (when can context transfer beat PerturbMean?)

Decompose the target shift as *δ*_*t*,*p*_ = *d*_*t*,*p*_ + *ɛ*^*r*^_*t*,*p*_, where *ɛ*^*r*^_*t*,*p*_ is the residual of the transfer prior. Let ℱ_*t*_ be the information available to a predictor at test time: the prior, the control cells of *t* and, in the partial-screen task, the screened knockdowns of *t*. Under squared error, the optimal prediction of *δ*_*t*,*p*_ is

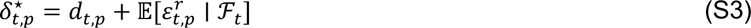

Its risk improves on that of PerturbMean by exactly 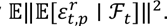 *Proof.* By the orthogonality of conditional expectation, 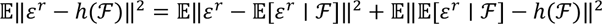 for any measurable ℎ. PerturbMean is ℎ ≡ 0. ∎ Consequently, *no* method, however expressive, can outperform PerturbMean in expectation unless the residual of the transfer prior is predictable from the target line’s control cells and screened knockdowns. The comparison between CASTER (and Linear) and PerturbMean is therefore an empirical estimate of 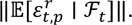 A null result means that this predictable part is small relative to estimation noise at the available data scale, not that context does not matter. D‖*ɛ*^*r*^ ‖^2^ (the line-specific variance quantified in Figure 1) can be large while its predictable part is small. In the positive-control simulation the residual is constructed to be a function of the control profile, so a gain over PerturbMean is possible.

## Note S1. Additional results, related to Figures 1 to 5

### Response sizes

Responses were largest in RPE1 and HepG2 (Figure S1A; median DEGs 2 in K562, 5 in HepG2, 2 in Jurkat and 29 in RPE1). This agrees with Nadig et al., who found the transcriptome-wide impact of these knockdowns to be largest in RPE1 and correlated across the four lines (average r = 0.62 for impact size)^S6^.

### Similarity and p53

Corrected medians per pair ranged from 0.47 to 0.66 and were highest for HepG2-RPE1. With all cells per knockdown, raw medians rose to 0.37 to 0.57, and corrected medians stayed within 0.01 (Table S15). No corrected value exceeded 1, and reliability was below 0.3 for only 0.2% of knockdown pairs. The bootstrap 95% interval of the cancer-line median was 0.51 to 0.54, and restricting to pairs with reliability ≥ 0.5 in both lines gave 0.55. Among knockdowns that induced p53 in neither HepG2 nor RPE1, HepG2-RPE1 remained the most similar pair (median r = 0.42), so shared p53 status goes with part, but not all, of their similarity; lineage and growth mode (both adherent and epithelial-derived) may contribute the rest. Nadig et al. noted the same pairing and the shared p53 wild-type, adherent state^S6^; our p53 target score tests this link per knockdown. The difference was not carried by p53 target genes themselves: after removing the 101 genes of a data-defined p53 module, HepG2-RPE1 similarity was 0.53 for knockdowns inducing p53 in both lines versus 0.44 for the others (with these genes: 0.57 versus 0.45). Knockdowns inducing p53 also had larger responses. When matched one to one on response size, they remained more similar (median difference 0.10, n = 200 pairs, Wilcoxon signed-rank p = 3.4 × 10⁻²¹). In a least-squares regression of similarity on response size and a p53 indicator, the p53 coefficient was 0.047 (p = 2.1 × 10⁻⁶). Within each quartile of response size, knockdowns inducing p53 in both lines were more similar (0.43 versus 0.36, 0.51 versus 0.46, 0.59 versus 0.55, 0.60 versus 0.57; Mann-Whitney p ≤ 0.043 in each quartile).

### Simulation and ablations

In simulations where the line-specific component was pure noise, CASTER stayed at the PerturbMean solution (Pearson delta 0.67 versus 0.68; Linear 0.61). When the line-specific component depended on each gene’s baseline expression, which is observable from control cells, CASTER recovered it (Pearson delta 0.84 versus 0.49), and so did Linear (0.86; Figure S3; simulation design in STAR Methods). The simulation therefore shows that CASTER can learn such structure without inventing it, not that it learns more than a linear model.

We ablated CASTER on the K562 partial-screen run with 20 training epochs, as in the main analysis, and two seeds for the main variants (Figure S4). The full model (S = 32) reached a Pearson delta of 0.201, compared with 0.187 for PerturbMean and 0.206 for Linear on the same split. The variants reached 0.177 (single cells, S = 1), 0.179 (S = 8), 0.189 (no attention), 0.201 (no FiLM), 0.100 (no transfer prior, learned embedding instead) and 0.199 (prior plus ID embedding). On Pearson delta (paired Wilcoxon, Holm), the full model was significantly better than the single-cell (S = 1), S = 8, no-attention, no-prior and prior-plus-ID variants. Removing the transfer prior was by far the most damaging change; the contributions of attention and FiLM were small (Table S4). These ablations come from a single held-out line and should be read as indicative.

### Partial-screen details

Restricted to the three cancer lines, Pearson delta was 0.239 for CASTER, 0.244 for Linear, 0.210 for LineOffset and 0.196 for PerturbMean. On the first seed, CASTER was significantly better than PerturbMean and ContextMean on the main metrics except discrimination against PerturbMean, and better than Linear only on DE AUPRC (one-sided paired Wilcoxon, Holm; Table S17). On the first seed, CASTER versus Linear gave Pearson delta 0.278 versus 0.282 (Holm p = 1.000), discrimination 0.430 versus 0.422 (Holm p = 0.117) and DE AUPRC 0.198 versus 0.181 (Holm p = 1.6 × 10⁻¹⁷). Across seeds, CASTER beat PerturbMean on Pearson delta in every seed (two-sided paired Wilcoxon p = 1.2 × 10⁻¹⁵², 1.3 × 10⁻¹²⁵, 1.5 × 10⁻¹⁵⁷), whereas the comparison with Linear was inconsistent (p = 0.081, 1.9 × 10⁻⁵¹, 0.755; median differences 0.000, −0.007, 0.001). A Wilcoxon test is not valid for predicted cells that are made by adding a shift to control cells. It called a mean of 1,333 DEGs per CASTER prediction, against 61 observed (Figure S16). DE metrics therefore use a moment-matched test on the predicted shift (STAR Methods). Conclusions are the same under both tests (Table S17). To compare CASTER with Linear directly, we bootstrapped knockdowns (seeds averaged) for the difference in the raw top-100 Pearson delta against all cells, with a margin of 5% of each line’s dynamic range: CASTER minus Linear was −0.017 in HepG2 (−0.027 to −0.008), −0.019 in Jurkat (−0.028 to −0.010), −0.024 in K562 (−0.032 to −0.016) and −0.035 in RPE1 (−0.049 to −0.021). In raw units the margin was 0.026 in HepG2, 0.025 in Jurkat, 0.023 in K562 and 0.040 in RPE1, so Linear was ahead in every line by an amount close to the margin. Per line (mean over seeds), Pearson delta for CASTER, PerturbMean and Linear was 0.200, 0.186 and 0.205 in K562; 0.320, 0.245 and 0.330 in HepG2; 0.197, 0.157 and 0.197 in Jurkat; and 0.387, 0.204 and 0.397 in RPE1 (Figure 3A; first-seed values in Table S16; secondary metrics in Figure S7). Among the secondary metrics, MAE, fold-change Spearman and direction agreement favored CASTER over PerturbMean (Table S17). DE overlap favored PerturbMean in three of four lines (Table S16). The tests in Table S17 are one-sided (CASTER better), so they cannot show that a baseline was better. The line-label CASTER’s output is in expression units, so its response was taken relative to the mean control profile and not to the shuffled input.

### Noise-free rendering

With noise-free rendering, CASTER remained above PerturbMean (91% of knockdowns better, p < 10⁻³⁰⁰) and LineOffset (88%, p < 10⁻³⁰⁰), and was close to Linear (39% better, median difference −0.010, p = 9.0 × 10⁻¹³⁶). Without rendering noise, ContextMean moved above PerturbMean and LineOffset in all four lines, and Linear stayed above CASTER in all four.

### Calibrated metrics

At 32 cells per knockdown, the DRF of Pearson delta and MAE was near zero or negative (−0.04 and −0.24; Figure 3D). This supports the finding of Miller et al.^S7^ that these two metrics are poorly calibrated. Pearson delta is the main metric of this and many other benchmarks. On the calibrated metrics at 32 cells (as rendered), CASTER was better than Linear for 50% of knockdowns on NIR (p = 5.4 × 10⁻⁴) and 49% of knockdowns on PDS (p = 0.127) (Table S9), so the two were close.

Against all cells, the DRF rose to 0.11 for Pearson delta (32 cells: −0.04), −0.09 for MAE (32 cells: −0.24), 0.38 for top-100 Pearson delta (32 cells: 0.11) and 0.25 for weighted R² relative to the perturbed mean (32 cells: 0.04). Against all cells, Pearson delta (noise-free predictions) was 0.511 for CASTER, 0.528 for Linear, 0.370 for LineOffset, 0.305 for PerturbMean and 0.438 for ContextMean. All metrics against all cells are in Table S18. With all cells, 5% to 11% of genes were differentially expressed per knockdown; DE AUPRC was highest for Linear and CASTER (0.35 and 0.34, versus 0.09 for NoChange), whereas DE overlap was highest for PerturbMean and LineOffset, which call many genes.

### Single-source transfer

In single-source transfer, CASTER showed the same pattern as PerturbMean: Pearson delta was 0.14 for cancer-to-cancer directions and 0.17 for directions with RPE1 (paired Wilcoxon p = 1.4 × 10⁻⁵⁶). In these four lines, transfer followed shared p53 status and growth mode more than transformation status. With one non-transformed line, these factors cannot be separated further.

### Ribosome examples and related genes in training

The knockdowns where CASTER lost most against PerturbMean were *SKP1* in HepG2, *SUDS3* in Jurkat and *OGFOD1* in K562 (r over DEGs: −0.51 versus 0.15, 0.18 versus 0.46 and 0.43 versus 0.62; Figure S11); none is a ribosome gene. In HepG2, for example, the PerturbMean prediction for *NVL* knockdown was uncorrelated or anti-correlated with the observed DEGs, whereas CASTER recovered their direction and relative magnitude.

When whole STRING-defined complexes were held out together, so that no sibling was in training, the median gain for ribosome knockdowns was 0.165 in K562, 0.183 in HepG2, 0.139 in Jurkat, 0.410 in RPE1, versus −0.004, 0.009, 0.011, 0.118 for other knockdowns (Figure S12B; the RPE1 bar has no ribosomal-protein knockdowns, because that group went to training in RPE1). Because members of the same complex respond similarly, a held-out ribosome gene could benefit from siblings in the training fraction. The gain correlated with the number of same-complex genes in training in Jurkat (rho = 0.38) but not clearly in the other lines (Figure S12C).

In the grouped splits, no test knockdown had a same-group sibling in training in any line or seed, whereas random splits had 677 to 789 such knockdowns per line. The largest translation group (131 genes: all 73 ribosomal-protein targets plus 58 translation-initiation, signal-recognition and ribosome-biogenesis factors) fell entirely into the test set in K562, HepG2 and Jurkat and into training in RPE1 (first seed). In HepG2 (first seed), the grouped split had the same sizes as the random split (320, 64 and 894 knockdowns), because whole groups were added until 25% was reached and the last one added was a single gene; the two training sets shared 80 of 320 knockdowns.

### Essentiality in predicted responses

The dependency-response coupling itself was preserved in predicted responses. On the held-out knockdowns, predicted response size correlated with the Chronos score at rho = −0.41 (K562), −0.52 (HepG2) and −0.33 (Jurkat) for CASTER, against −0.44, −0.54 and −0.43 for observed responses of the same knockdowns and −0.51, −0.46 and −0.33 for PerturbMean. ContextMean, which ignores knockdown identity, lost it (Figure S17).

---

**Figure S1.**
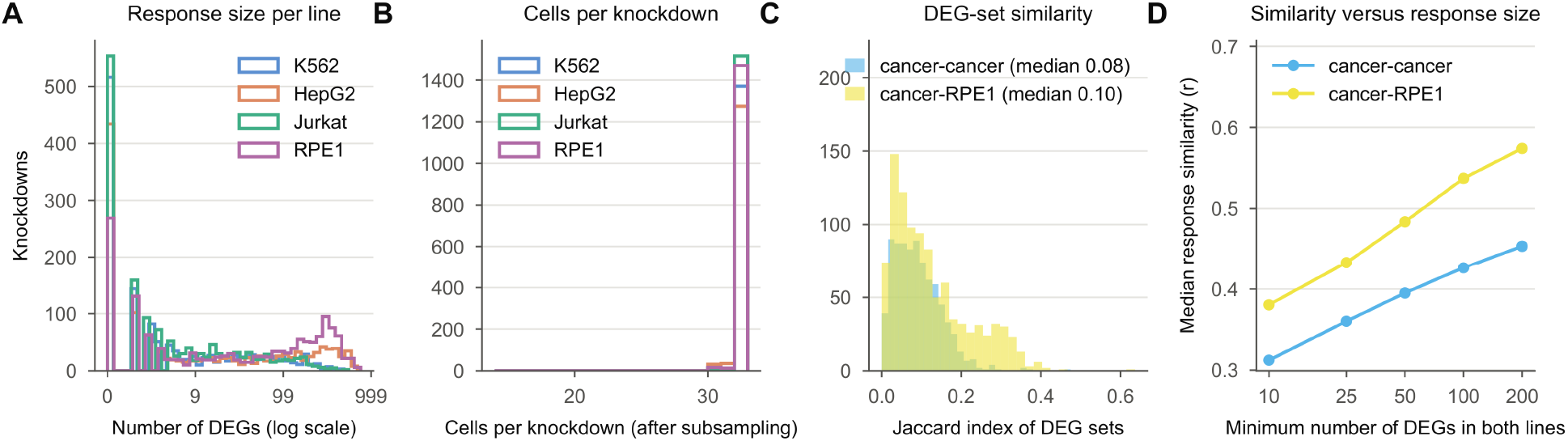
Response sizes and line overlap, related to Figure 1. (A) Distribution of the number of DEGs per knockdown in each line. (B) Cells per knockdown after subsampling. (C) DEG-set similarity (Jaccard index) between lines. (D) Median cross-line response similarity as a function of response size (minimum number of DEGs in both lines).

**Figure S2.**
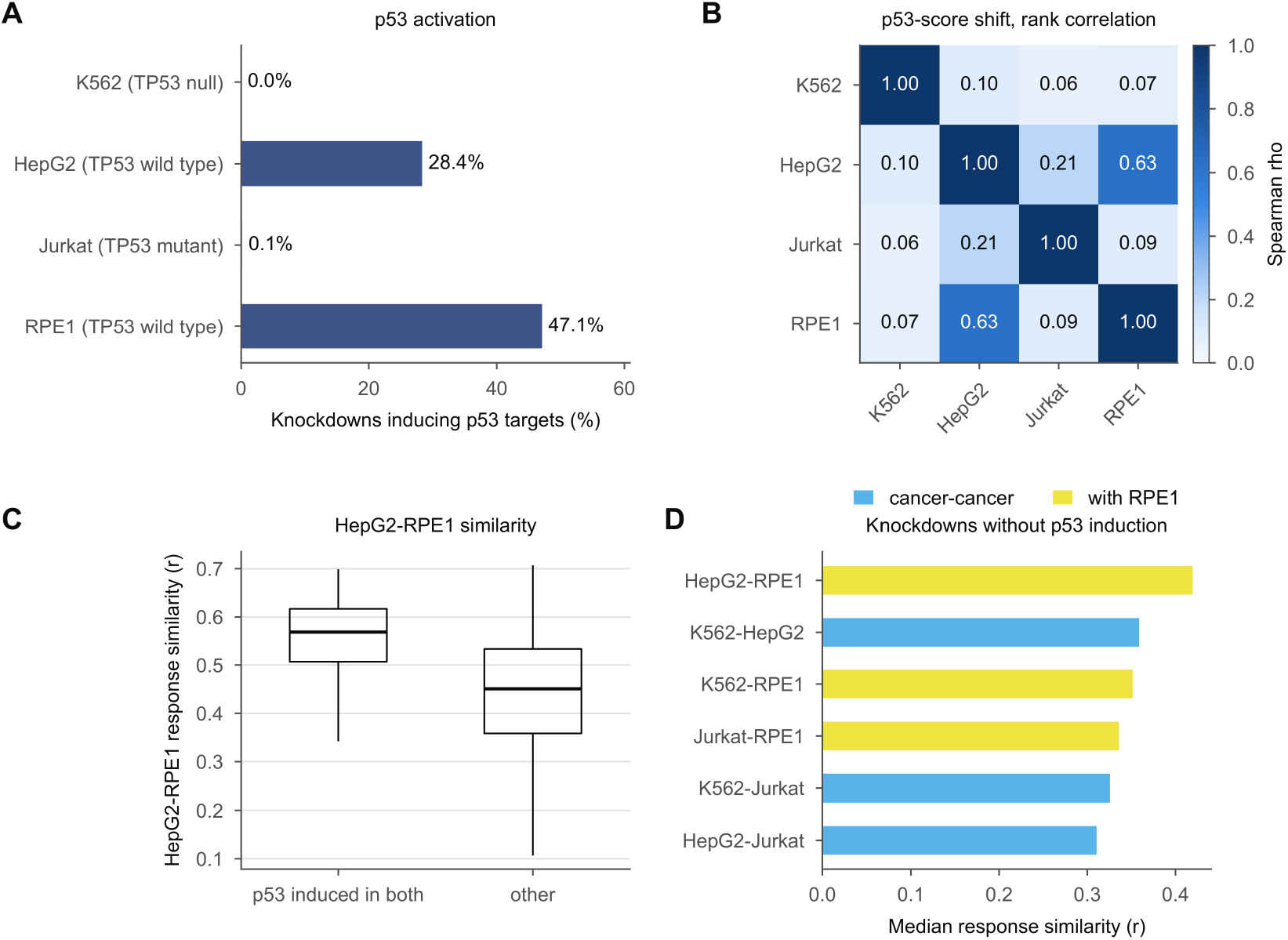
p53 activity across lines, related to Figures 1 and 4. (A) Percentage of knockdowns inducing the 14-gene p53 target score (FDR < 0.05), with *TP53* status. (B) Rank correlation of p53-score shifts between lines (knockdowns observed in both). (C) HepG2-RPE1 response similarity for knockdowns inducing p53 in both lines versus others (two-sided Mann-Whitney test; p value in the text). (D) Median response similarity per line pair among knockdowns inducing p53 in neither line; cancer-cancer pairs in sky blue, pairs involving RPE1 in yellow, as in Figure 1D.

**Figure S3.**
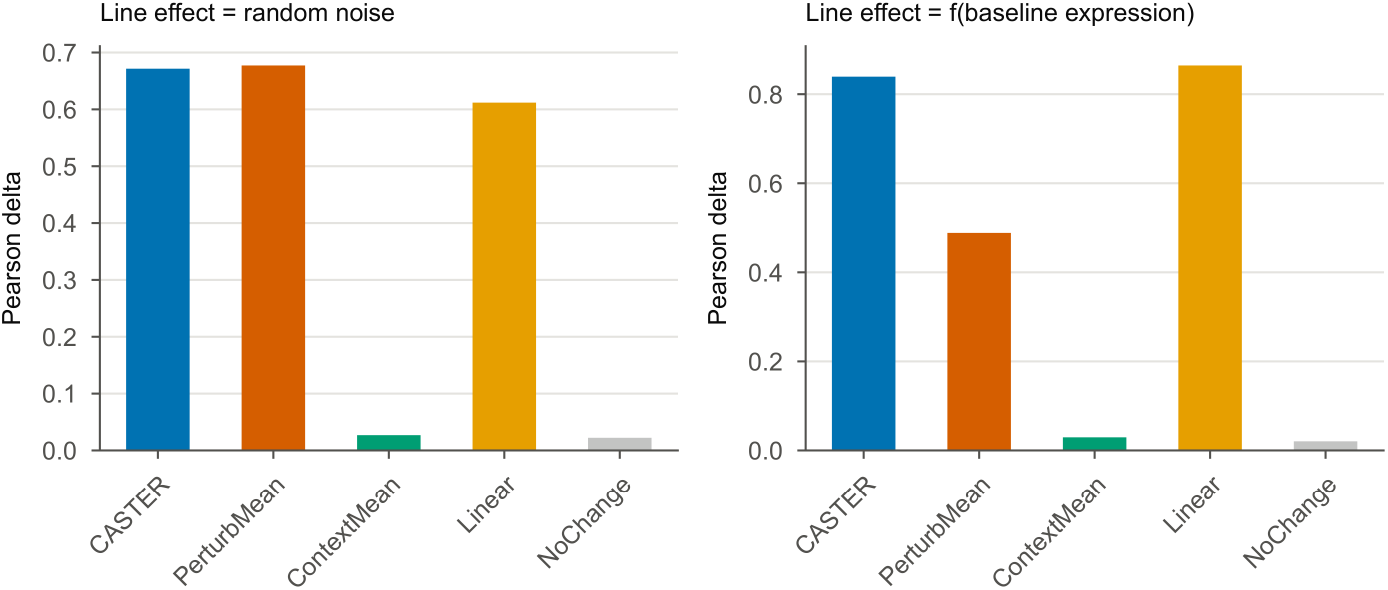
Positive-control simulation, related to Figure 1. Synthetic four-line data in which the line-specific component is random noise (left) or a function of baseline expression observable from control cells (right).

**Figure S4.**
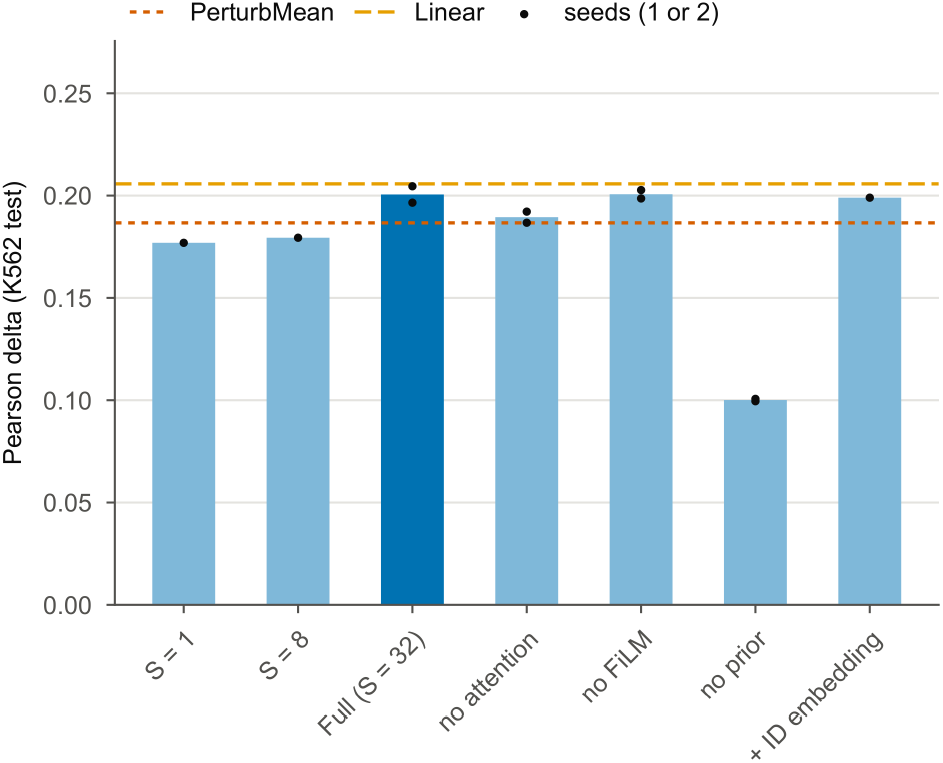
CASTER ablations, related to Figure 1. Held-out Pearson delta for set sizes and architectural ablations on the K562 partial-screen run (20 epochs; bars, mean over seeds; dots, seeds, one or two per variant), with PerturbMean (dotted) and Linear (dashed) on the same split (two seeds). Values as in Table S4.

**Figure S5.**
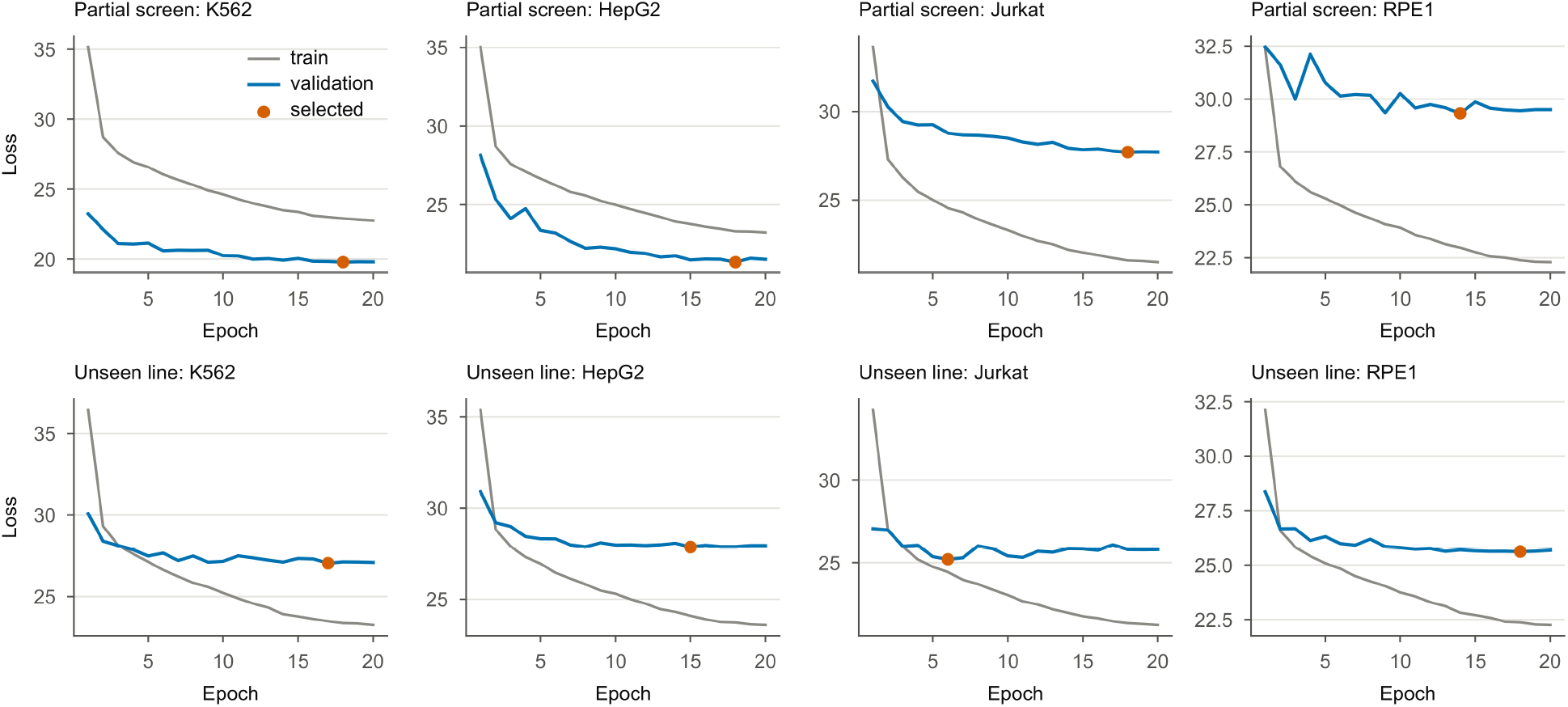
Training curves, related to Figures 2 and 4. Training and validation loss per epoch for every partial-screen (top) and unseen-line (bottom) run; the selected checkpoint is marked.

**Figure S6.**
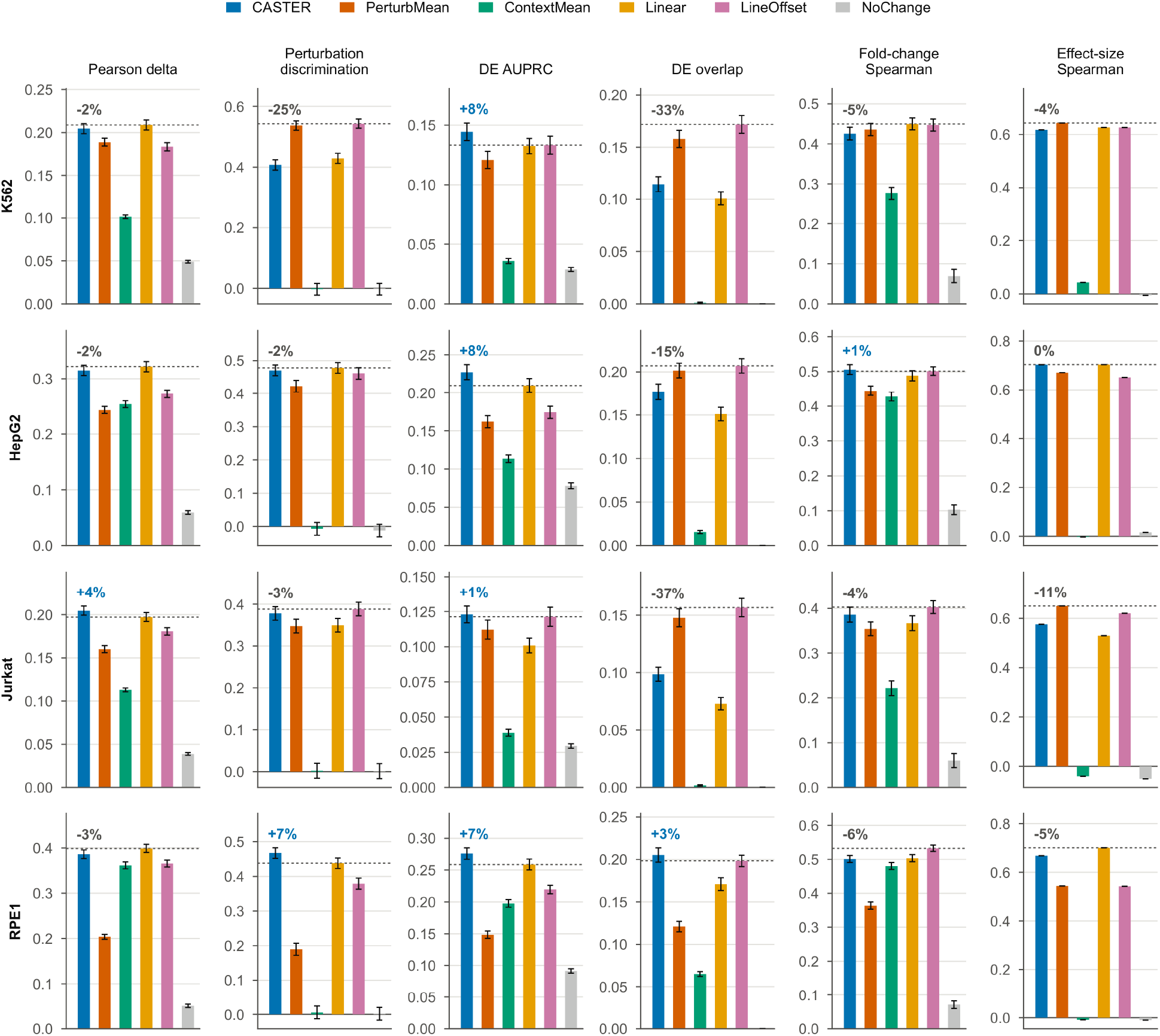
All 32-cell metrics per target line, partial-screen task, related to Figure 2. First seed; error bars, SEM over knockdowns (effect-size Spearman is a single value per line); percentages, relative difference of CASTER from the best other method (dashed line).

**Figure S7.**
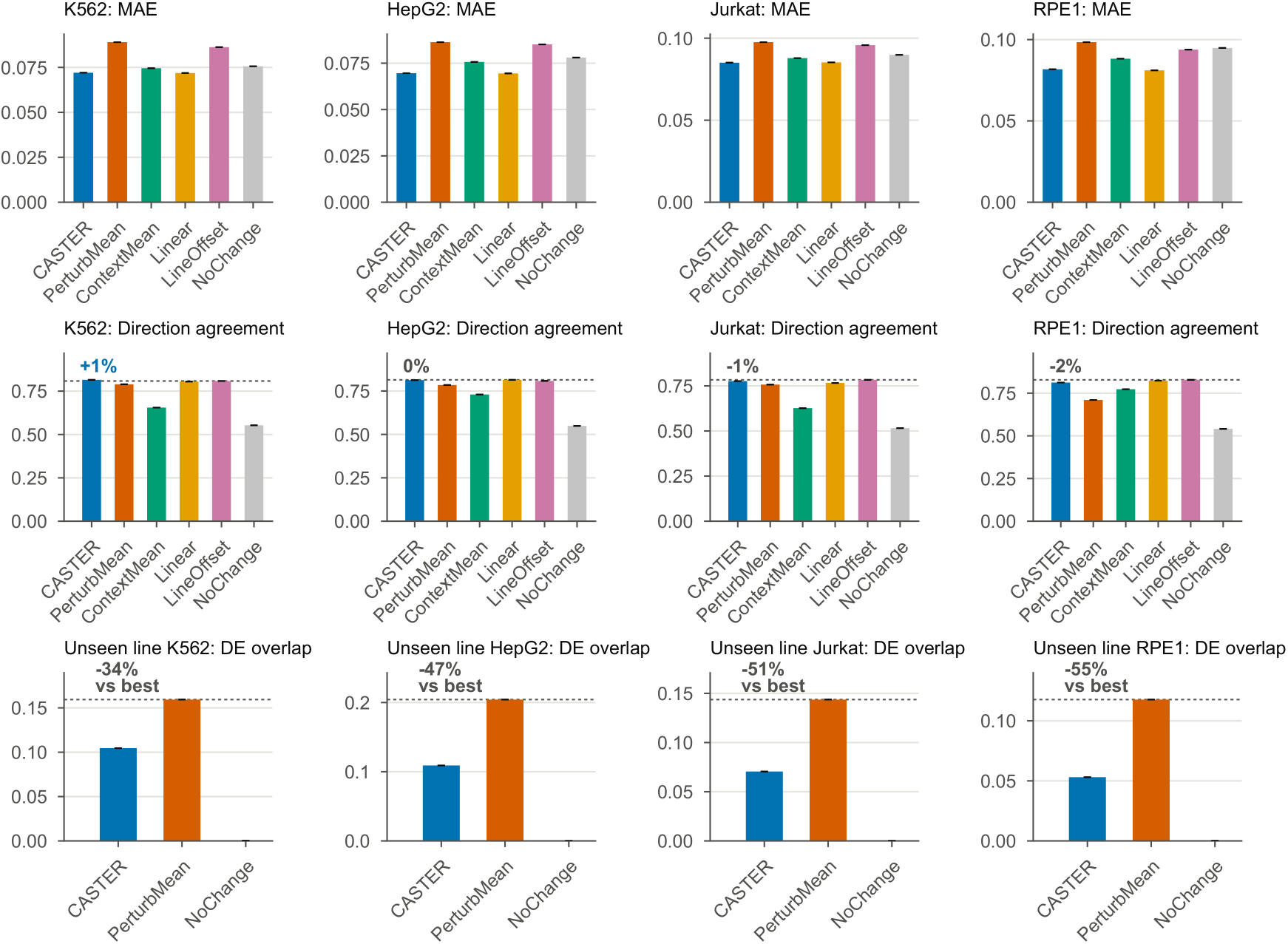
Secondary metrics, related to Figures 2 and 4. Mean absolute error and direction agreement per line (partial-screen task), and DE overlap per line in the unseen-line task.

**Figure S8.**
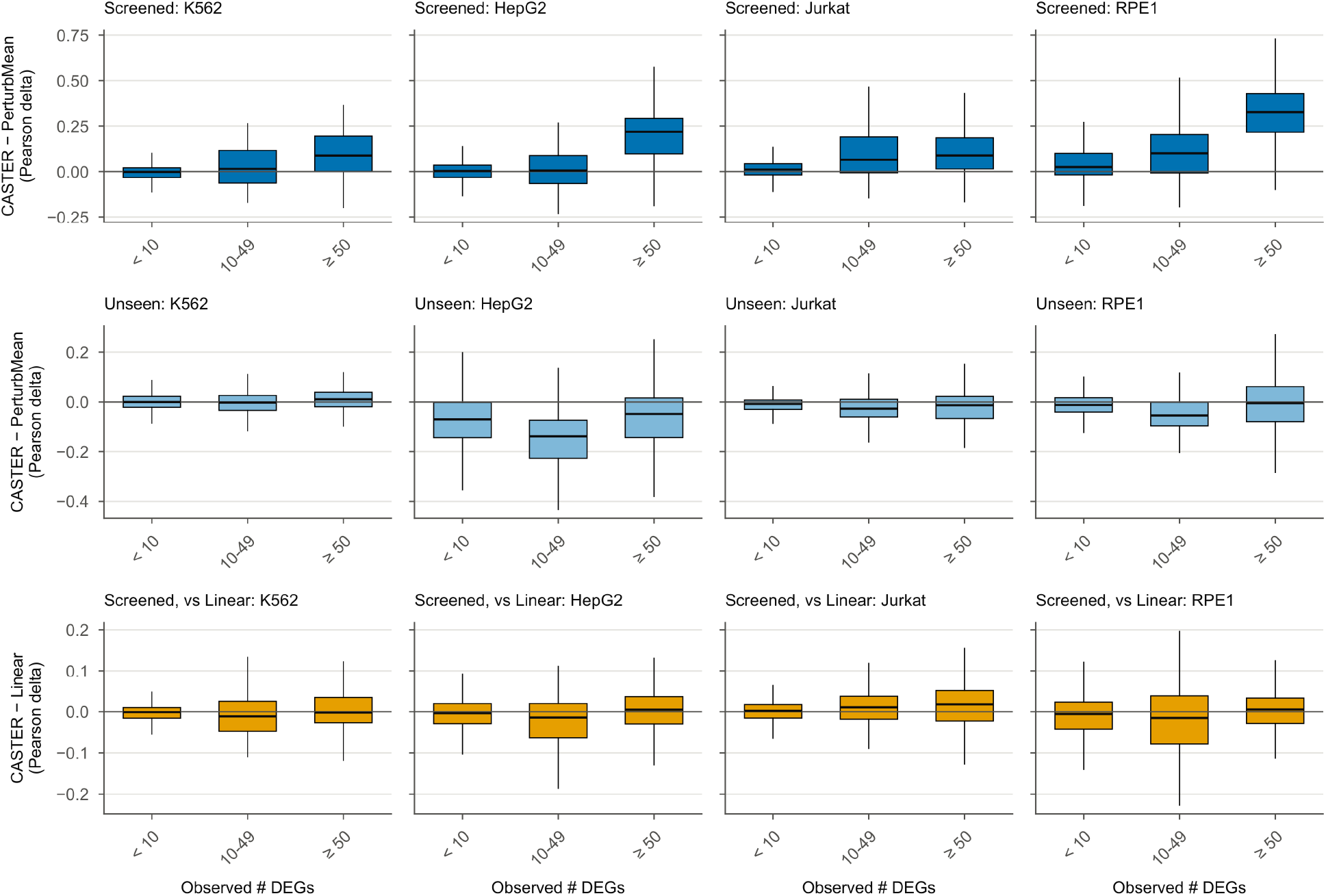
CASTER’s gain as a function of response size, related to Figures 2 and 4. Per-knockdown difference in Pearson delta, binned by the observed number of DEGs (< 10, 10 to 49, ≥ 50): CASTER minus PerturbMean when 30% of the line is screened (top row) and in unseen lines (middle row), and CASTER minus Linear when 30% is screened (bottom row). Boxes, IQR; whiskers, 1.5 × IQR.

**Figure S9.**
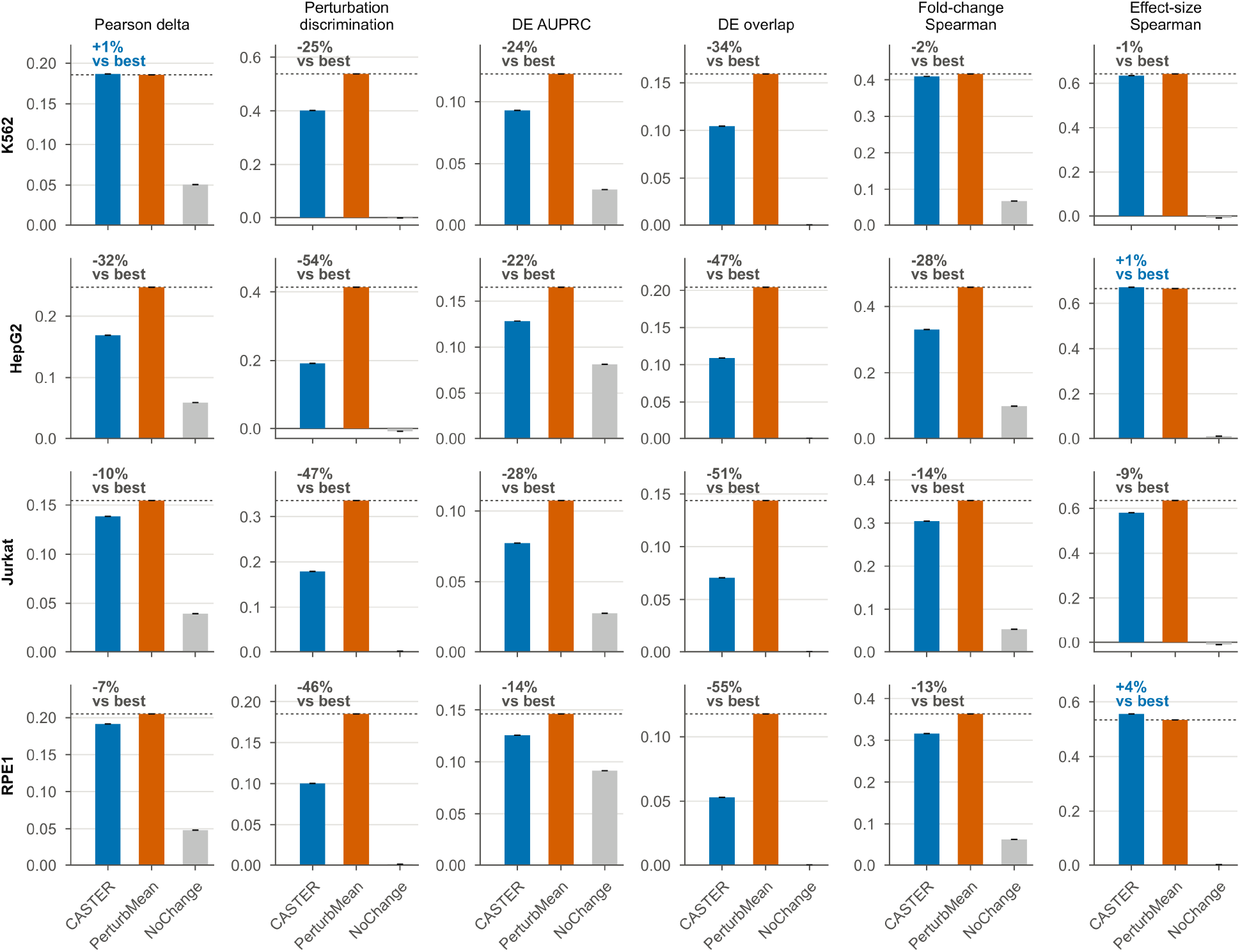
All metrics per target line in the unseen-line task, related to Figure 4. As Figure S6, for the unseen-line task (CASTER, PerturbMean and NoChange; one seed).

**Figure S10.**
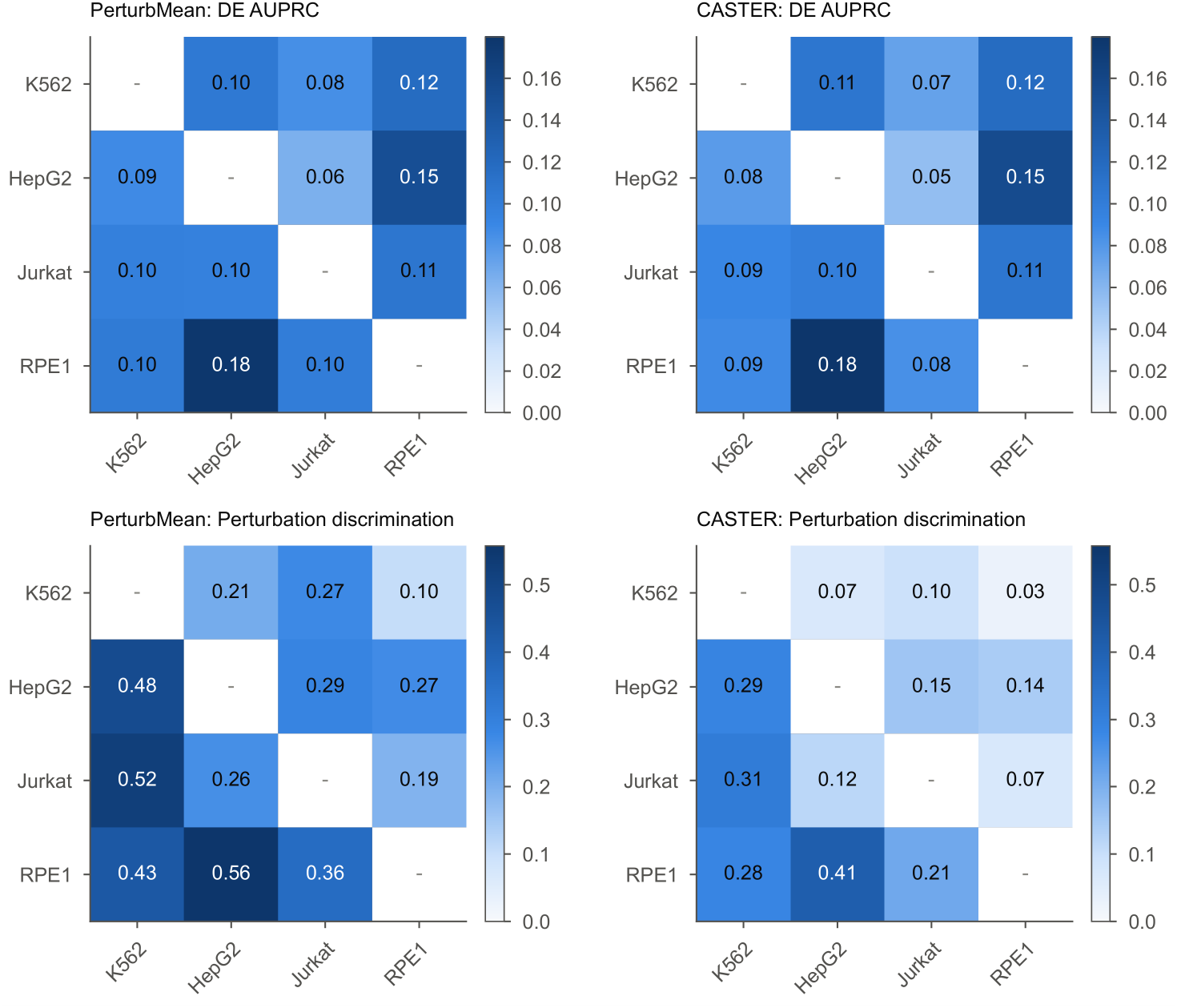
Single-source transfer, all metrics, related to Figure 4. DE AUPRC and perturbation discrimination for every source and target pair.

**Figure S11.**
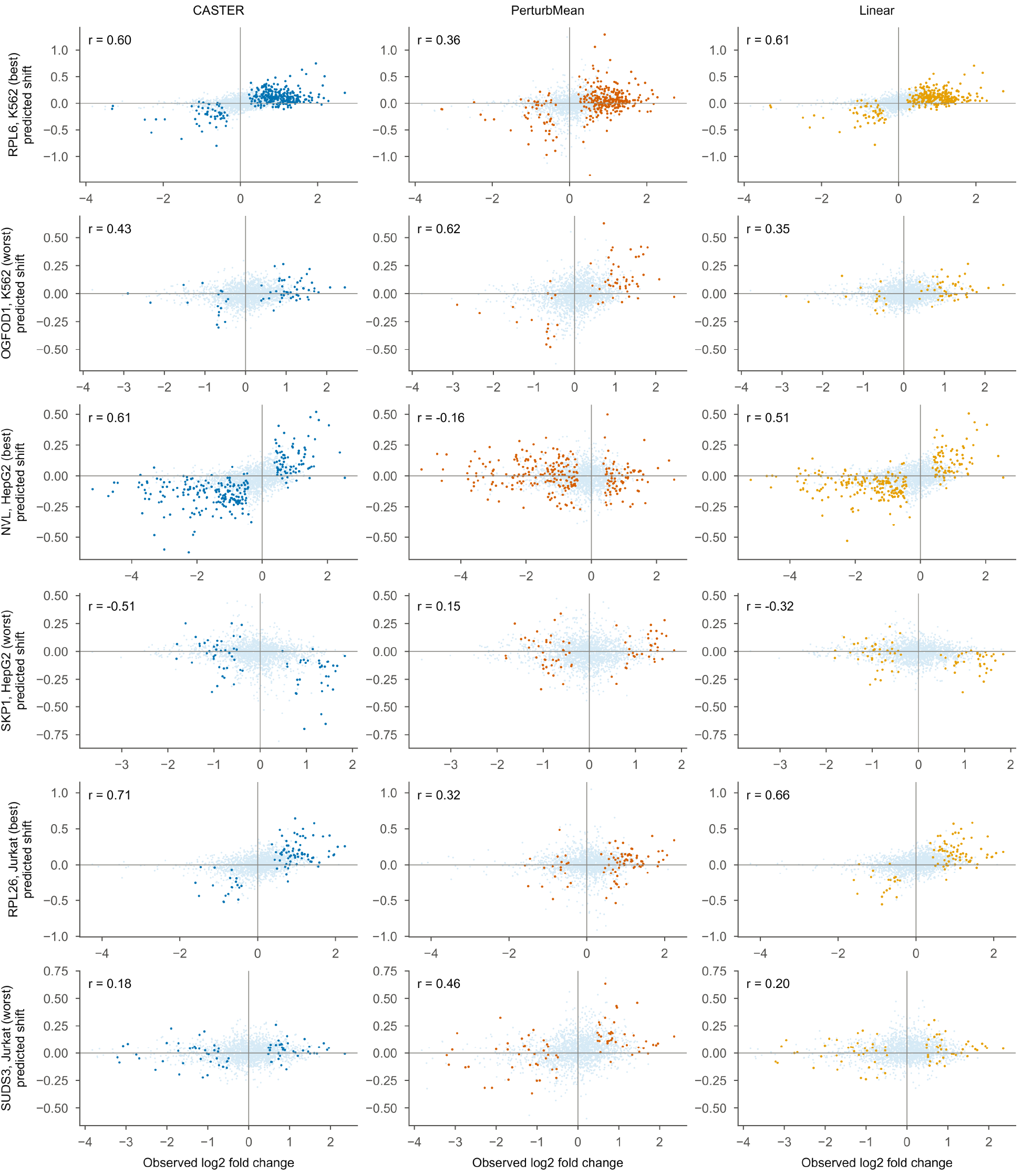
Line-specific example knockdowns, related to Figure 5. For each cancer line, the knockdown (≥ 30 DEGs) with the largest CASTER gain over PerturbMean (best) and the one with the largest loss (worst): observed log₂ fold change versus predicted pseudobulk shift; colored, observed DEGs.

**Figure S12.**
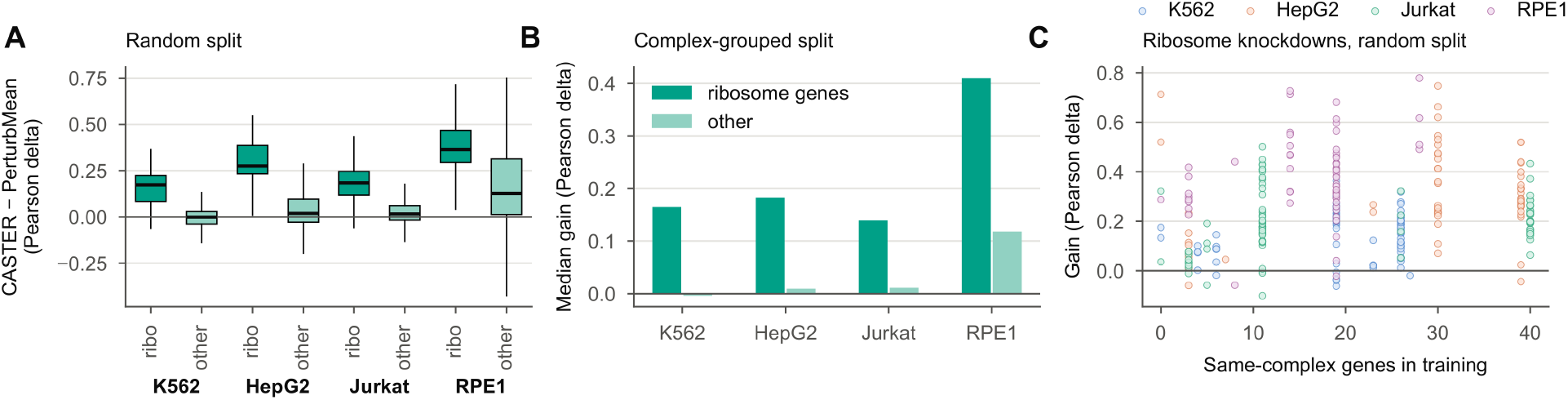
Ribosome knockdowns and related genes in training, related to Figure 5. (A) CASTER gain over PerturbMean for ribosome-biogenesis and ribosomal-protein knockdowns versus others (random split, first seed). (B) Median gain under complex-grouped splits, where no same-complex gene is in training; the RPE1 ribosome bar has no ribosomal-protein knockdowns, because that group went to training in RPE1. (C) Gain versus the number of same-complex genes in training (random split).

**Figure S13.**
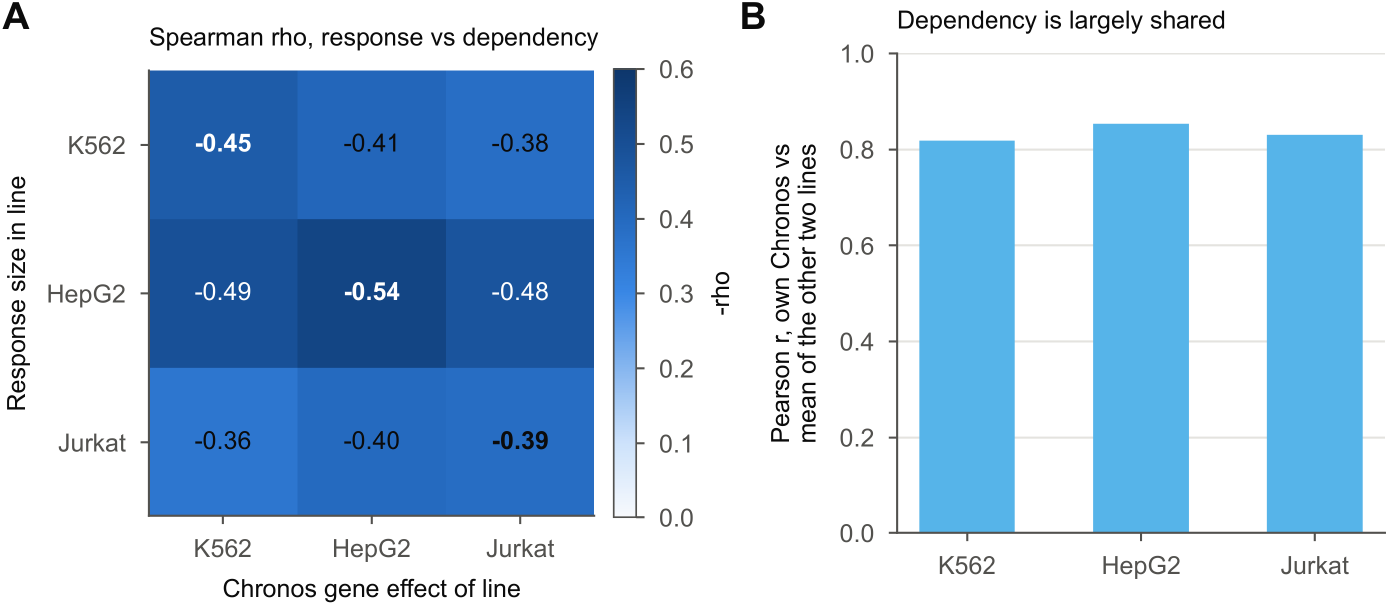
The dependency-response link is not line-specific, related to Figure 6. (A) Spearman rho between response size in one line (rows) and Chronos score in each line (columns). (B) Pearson r between each line’s Chronos scores and the mean of the other two lines.

**Figure S14.**
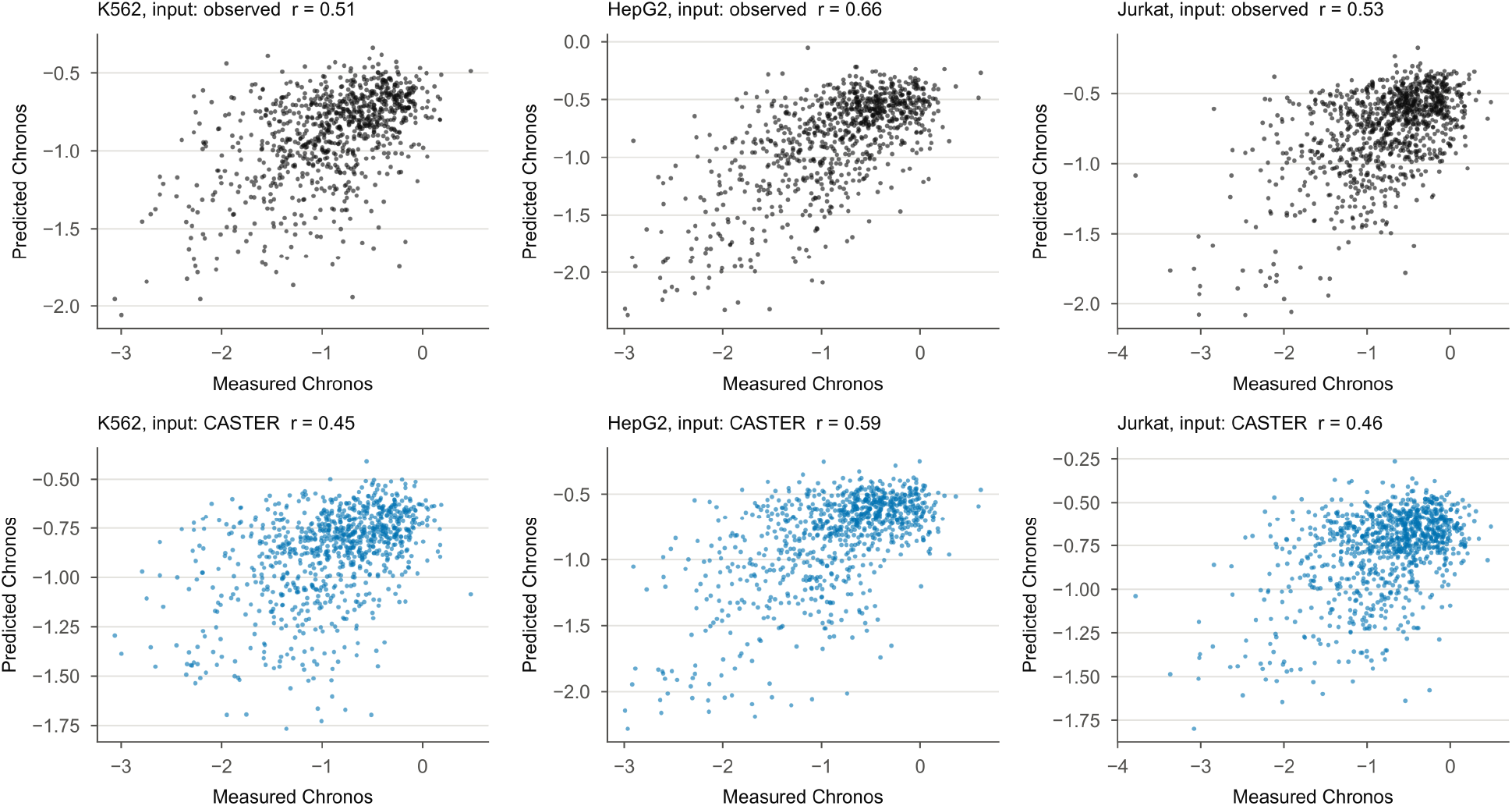
Dependency prediction from transcriptional shifts, related to Figure 6. Measured versus predicted Chronos score per cancer line, from observed shifts (top) and from CASTER-predicted shifts (bottom).

**Figure S15.**
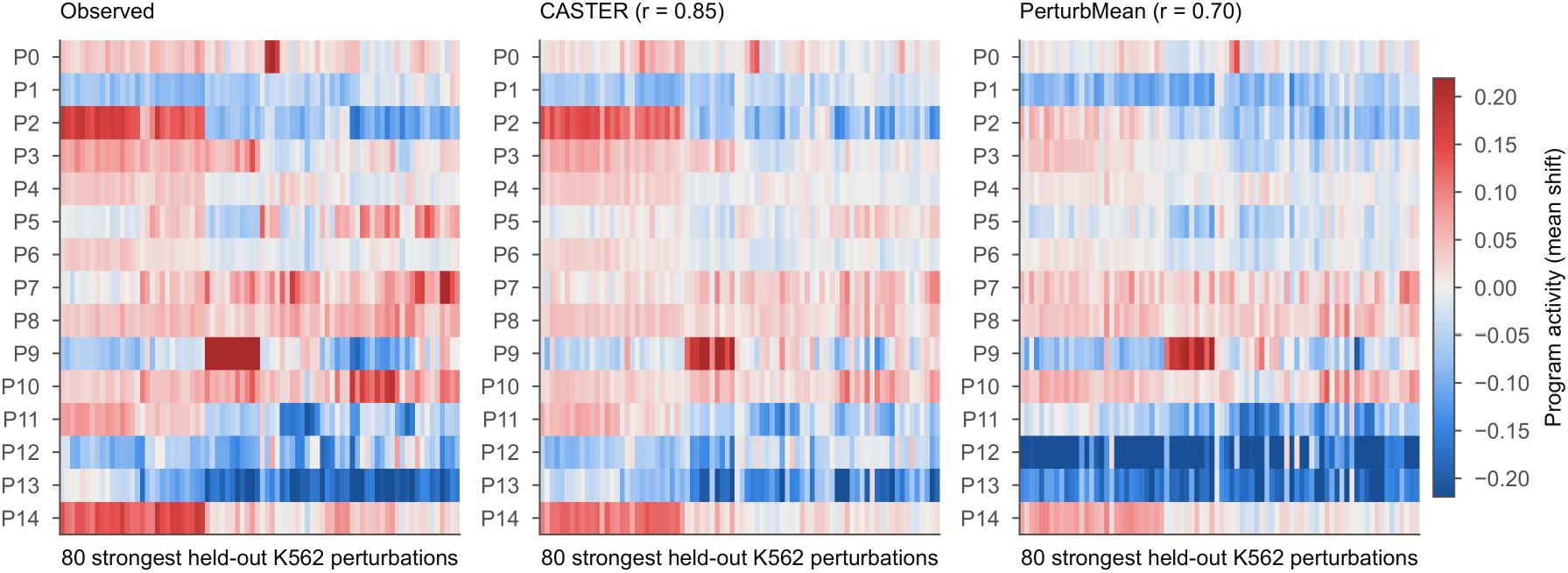
Gene-program activity of held-out knockdowns, related to Figure 6. Activity of the 15 K562 gene programs (rows) for the 80 strongest held-out K562 knockdowns (columns, clustered on the observed matrix): observed, CASTER and PerturbMean. r, Pearson correlation with the observed matrix. Values below the color scale are shown at its lower end.

**Figure S16.**
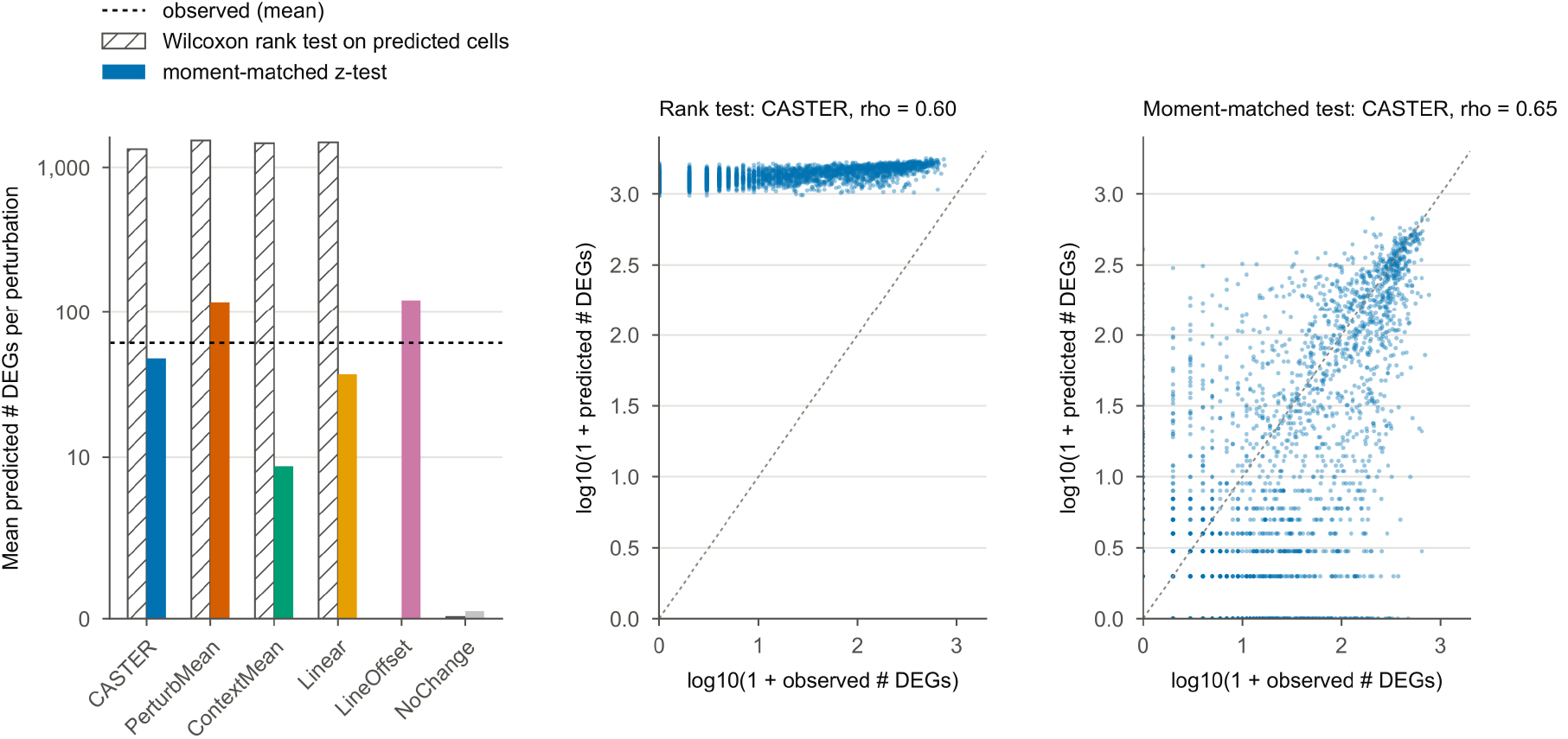
Predicted-side differential expression: rank test versus moment-matched test, related to STAR Methods. (Left) Mean number of predicted DEGs per knockdown under the Wilcoxon rank test on predicted cells and the moment-matched z test (method colors); dashed line, observed mean. LineOffset was not run with the rank test. Shifted predictions turn exact zeros into small positive values, so the rank test calls most genes significant; unshifted control cells (NoChange) are unaffected. (Middle, right) Observed versus CASTER-predicted DEG counts per knockdown under the two tests.

**Figure S17.**
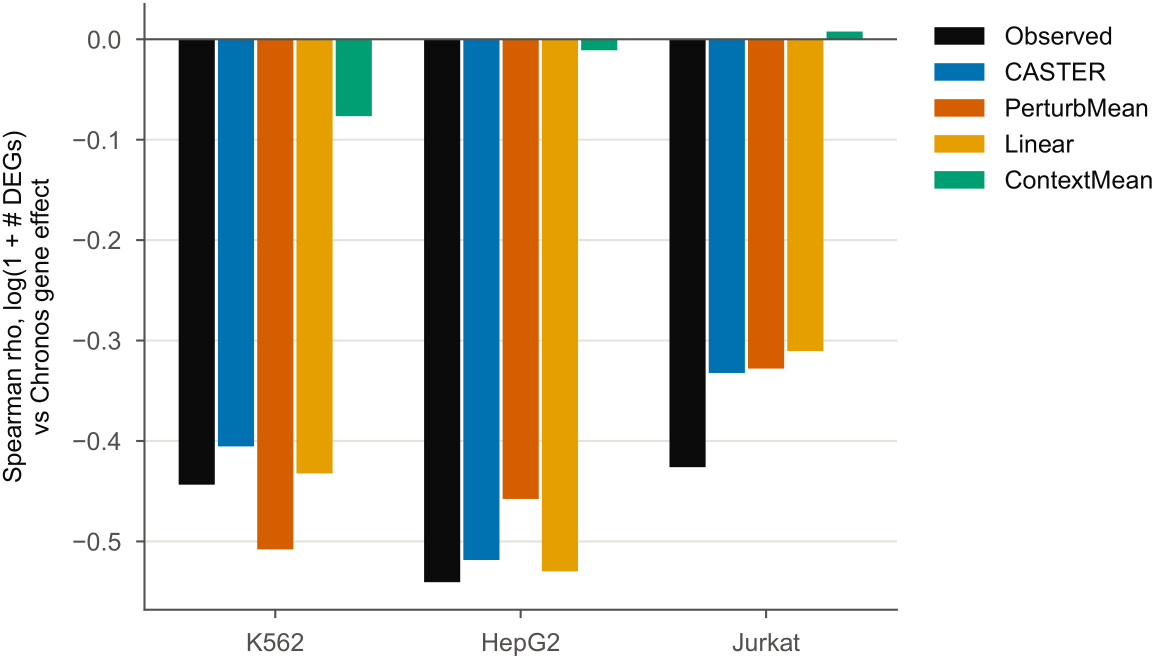
Dependency-response coupling in predicted responses, related to Figure 6. Spearman correlation between response size (log(1 + number of DEGs)) and Chronos score for observed responses and for the responses predicted by each method (held-out knockdowns, partial-screen task).

**Table S1.** Dataset after subsampling and filtering, related to Figure 1.

| **Cell line** | **Cells (knockdown plus control)** | **Control cells** | **Knockdowns** | **Median cells per knockdown** | **Median number of DEGs** | **Knockdowns with ≥ 10 DEGs (%)** |
|---|---|---|---|---|---|---|
| K562 | 47,236 | 3,000 | 1,383 | 32 | 2 | 32 |
| HepG2 | 45,783 | 3,000 | 1,340 | 32 | 5 | 46 |
| Jurkat | 52,151 | 3,000 | 1,537 | 32 | 2 | 32 |
| RPE1 | 50,923 | 3,000 | 1,499 | 32 | 29 | 60 |

**Table S2.**
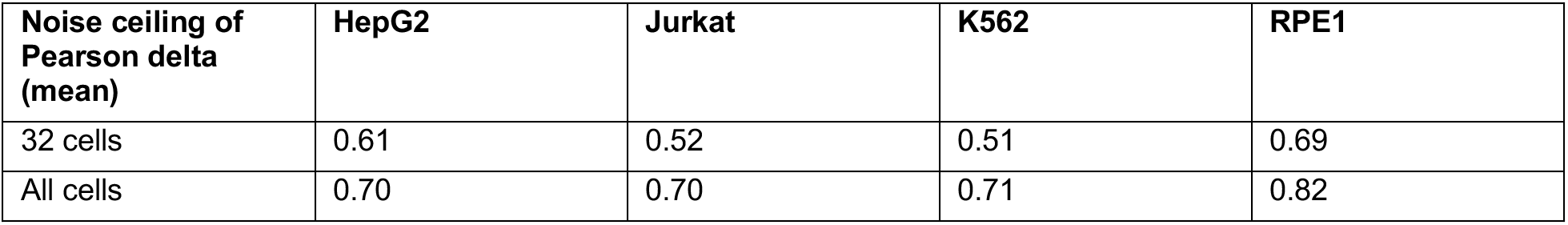
Noise ceilings of Pearson delta per line, related to Figure 3. Mean over held-out knockdowns of the square root of the Spearman-Brown-corrected split-half reliability, at 32 cells and against all cells.

**Table S3.** CASTER hyperparameters and training settings, related to STAR Methods.

| Component | Setting |
|---|---|
| Cell encoder | 4-layer MLP, GELU, 2,000 → 256 → 256 → 256 → 256 |
| Prior encoder | 2-layer MLP, GELU, 2,000 → 256 → 256 |
| Line summary and FiLM | mean of cell embeddings → 2-layer MLP → (γ, β), each of size 256 |
| Set transformer | 4 layers, 8 heads, width 256, feed-forward 512, pre-norm, GELU, dropout 0, no positional encoding |
| Output | LayerNorm + linear 256 → 2,000 (zero-initialized); g(c): linear 256 → 2,000 (zero-initialized) |
| Cell-set size S | 32 (ablation: 1, 8) |
| Loss | energy-distance MMD² + 1.0 × pseudobulk squared error |
| Optimizer | AdamW, learning rate 3 × 10⁻⁴, weight decay 10⁻⁴, batch 32 sets, gradient-norm clipping at 1.0 |
| Schedule | 5% linear warm-up, cosine decay to 5% of peak |
| Epochs | 20 (partial-screen and unseen-line tasks, ablations, learning curve); 10 (single-source transfer) |
| Model selection | lowest validation loss after each epoch (partial screen: the target line's validation knockdowns; unseen line and single-source transfer: 5% of source items) |
| Hardware | CPU only (4 cores, 8 GB RAM), PyTorch 2.13.0 |
| Parameters | 4.6 million (excluding the one-hot embedding, used only in ablations) |

**Table S4.** Ablations, related to Figure 1. Partial-screen task, K562 target line, 20 epochs; mean over seeds.

| **Variant** | **Seeds** | **Pearson delta** | **Perturbation discrimination** | **DE AUPRC** | **DE overlap** | **Fold-change Spearman** |
|---|---|---|---|---|---|---|
| Full model (S = 32) | 2 | 0.201 | 0.392 | 0.141 | 0.113 | 0.424 |
| S = 8 | 1 | 0.179 | 0.291 | 0.107 | 0.078 | 0.414 |
| Single cells (S = 1) | 1 | 0.177 | 0.436 | 0.117 | 0.151 | 0.420 |
| No attention | 2 | 0.189 | 0.329 | 0.127 | 0.093 | 0.409 |
| No FiLM | 2 | 0.201 | 0.389 | 0.139 | 0.111 | 0.421 |
| No transfer prior | 2 | 0.100 | 0.004 | 0.033 | 0.002 | 0.246 |
| Prior plus ID embedding | 1 | 0.199 | 0.389 | 0.137 | 0.114 | 0.421 |

**Table S5.** Learning curve and small-screen controls, related to Figure 2. Scaled top-100 Pearson delta against all cells (0 = NoChange, 1 = technical duplicate) for every method, screened fraction and line (mean over three seeds; per-seed values with the raw scores of each run, the technical duplicate and NoChange in Data S1), with the numbers of screened and test knockdowns. The 30% rows are the partial-screen task. Small-screen controls (2.5% to 10%): anchored ridge, PerturbMean plus a ridge on the residual from PerturbMean; scaled PerturbMean, PerturbMean with a gene-wise scale fitted on the screened knockdowns; Linear with 8 or 16 PCA components; mean-profile CASTER (2.5% only). NA, not run.

| Screened (%) | Cell line | Screened knockdowns | Test knockdowns | Linear | CASTER | LineOffset | PerturbMean | ContextMean | Anchored ridge | Scaled PerturbMean | Linear (8 PCs) | Linear (16 PCs) | Mean-profile CASTER |
|---|---|---|---|---|---|---|---|---|---|---|---|---|---|
| 2.5 | HepG2 | 32 | 894 | 0.56 | 0.67 | 0.39 | 0.47 | −0.49 | 0.54 | 0.52 | 0.54 | 0.54 | 0.70 |
| 2.5 | Jurkat | 37 | 1027 | 0.32 | 0.62 | 0.49 | 0.52 | −0.22 | 0.55 | 0.56 | 0.24 | 0.31 | 0.63 |
| 2.5 | K562 | 32 | 917 | 0.49 | 0.69 | 0.49 | 0.37 | −0.54 | 0.57 | 0.54 | 0.46 | 0.48 | 0.63 |
| 2.5 | RPE1 | 36 | 1003 | 0.53 | 0.50 | 0.47 | 0.35 | −0.04 | 0.57 | 0.39 | 0.49 | 0.51 | 0.56 |
| 5 | HepG2 | 64 | 894 | 0.70 | 0.71 | 0.44 | 0.49 | −0.44 | 0.62 | 0.55 | 0.64 | 0.66 | NA |
| 5 | Jurkat | 73 | 1027 | 0.58 | 0.62 | 0.50 | 0.52 | −0.26 | 0.60 | 0.59 | 0.52 | 0.56 | NA |
| 5 | K562 | 66 | 917 | 0.63 | 0.70 | 0.52 | 0.39 | −0.46 | 0.60 | 0.56 | 0.57 | 0.60 | NA |
| 5 | RPE1 | 72 | 1003 | 0.59 | 0.53 | 0.47 | 0.34 | −0.05 | 0.59 | 0.39 | 0.54 | 0.58 | NA |
| 10 | HepG2 | 127 | 894 | 0.78 | 0.78 | 0.46 | 0.50 | −0.37 | 0.67 | 0.57 | 0.69 | 0.73 | NA |
| 10 | Jurkat | 146 | 1027 | 0.72 | 0.68 | 0.50 | 0.52 | −0.25 | 0.65 | 0.60 | 0.63 | 0.68 | NA |
| 10 | K562 | 131 | 917 | 0.75 | 0.71 | 0.53 | 0.39 | −0.45 | 0.64 | 0.57 | 0.67 | 0.71 | NA |
| 10 | RPE1 | 143 | 1003 | 0.63 | 0.62 | 0.47 | 0.34 | −0.06 | 0.63 | 0.39 | 0.56 | 0.60 | NA |
| 20 | HepG2 | 256 | 894 | 0.88 | 0.86 | 0.45 | 0.50 | −0.37 | NA | NA | NA | NA | NA |
| 20 | Jurkat | 293 | 1027 | 0.81 | 0.79 | 0.51 | 0.52 | −0.23 | NA | NA | NA | NA | NA |
| 20 | K562 | 262 | 917 | 0.85 | 0.81 | 0.53 | 0.39 | −0.44 | NA | NA | NA | NA | NA |
| 20 | RPE1 | 287 | 1003 | 0.68 | 0.65 | 0.47 | 0.35 | −0.04 | NA | NA | NA | NA | NA |
| 30 | HepG2 | 384 | 894 | 0.91 | 0.88 | 0.45 | 0.49 | −0.40 | NA | NA | NA | NA | NA |
| 30 | Jurkat | 439 | 1027 | 0.85 | 0.81 | 0.51 | 0.52 | −0.21 | NA | NA | NA | NA | NA |
| 30 | K562 | 392 | 917 | 0.89 | 0.84 | 0.54 | 0.39 | −0.44 | NA | NA | NA | NA | NA |
| 30 | RPE1 | 430 | 1003 | 0.71 | 0.67 | 0.48 | 0.35 | −0.02 | NA | NA | NA | NA | NA |

**Table S6.** Per-line controls on the scaled top-100 Pearson delta against all cells, related to Figure 3. First seed, noise-free predictions; 0 = NoChange, 1 = technical duplicate. The mean-profile and line-label models are scored as output minus the line’s mean control profile. NIR and PDS use the perturbed mean as the negative control. The wrong-line CASTER (another line’s control cells, mean over two seeds) scored 0.36 on the mean top-100 row, versus 0.80 with the correct cells.

| **Line** | **Metric** | **CASTER** | **Mean-profile CASTER** | **Line-label CASTER** | **LineOffset** | **Linear** | **PerturbMean** |
|---|---|---|---|---|---|---|---|
| K562 | Top-100 Pearson delta† | 0.84 | 0.89 | 0.85 | 0.55 | 0.87 | 0.42 |
| K562 | NIR | 0.52 | 0.52 | 0.50 | 0.68 | 0.53 | 0.69 |
| K562 | PDS | 0.50 | 0.50 | 0.48 | 0.63 | 0.51 | 0.63 |
| HepG2 | Top-100 Pearson delta† | 0.88 | 0.91 | 0.88 | 0.44 | 0.91 | 0.48 |
| HepG2 | NIR | 0.57 | 0.55 | 0.58 | 0.53 | 0.57 | 0.49 |
| HepG2 | PDS | 0.55 | 0.54 | 0.57 | 0.52 | 0.56 | 0.48 |
| Jurkat | Top-100 Pearson delta† | 0.82 | 0.88 | 0.85 | 0.53 | 0.84 | 0.54 |
| Jurkat | NIR | 0.42 | 0.43 | 0.42 | 0.46 | 0.39 | 0.44 |
| Jurkat | PDS | 0.39 | 0.39 | 0.38 | 0.40 | 0.36 | 0.38 |
| RPE1 | Top-100 Pearson delta† | 0.66 | 0.67 | 0.65 | 0.48 | 0.71 | 0.35 |
| RPE1 | NIR | 0.49 | 0.50 | 0.46 | 0.39 | 0.46 | 0.16 |
| RPE1 | PDS | 0.47 | 0.48 | 0.44 | 0.35 | 0.43 | 0.15 |
| Mean | Top-100 Pearson delta† | 0.80 | 0.84 | 0.81 | 0.50 | 0.83 | 0.45 |
| Mean | NIR | 0.50 | 0.50 | 0.49 | 0.52 | 0.49 | 0.45 |
| Mean | PDS | 0.48 | 0.48 | 0.47 | 0.48 | 0.47 | 0.41 |

**Table S7.** Robustness checks, related to Figure 3.

| **Check** | **As rendered** | **Noise-free** | **CASTER as rendered** |
|---|---|---|---|
| PerturbMean: Pearson delta rendered on sampled versus noise-free control cells (first seed) | 0.198 | 0.252 | 0.278 |
| Linear: Pearson delta rendered on sampled versus noise-free control cells (first seed) | 0.282 | 0.437 | 0.278 |
| LineOffset: Pearson delta rendered on sampled versus noise-free control cells (first seed) | 0.251 | 0.314 | 0.278 |
| Cross-line response similarity on all 6,546 genes, cancer-cancer (median r; n = 813) | 0.36 | NA | NA |
| Cross-line response similarity on all 6,546 genes, with RPE1 (median r; n = 988) | 0.40 | NA | NA |

**Table S8.** Calibrated metrics against one half of each knockdown’s cells, partial-screen task, related to Figure 3. Mean over lines and seeds (32 cells, as rendered). The technical and interpolated duplicates are positive controls; the perturbed mean (the mean of the screened knockdowns) is the negative control, and NoChange (an exact zero shift) for the metrics relative to the perturbed mean (†). Raw metric means; scaled scores and both DRF definitions are in Table S9.

| **Method** | **Pearson delta** | **Top-100 Pearson delta†** | **Weighted R²†** | **WMSE** | **MSE** | **MAE** | **NIR** | **PDS** |
|---|---|---|---|---|---|---|---|---|
| Technical duplicate | 0.291 | 0.457 | −0.920 | 0.0139 | 0.0180 | 0.101 | 0.900 | 0.884 |
| Interpolated duplicate | 0.419 | 0.455 | −0.920 | 0.0139 | 0.0103 | 0.074 | 0.843 | 0.858 |
| CASTER | 0.236 | 0.308 | −0.953 | 0.0143 | 0.0157 | 0.093 | 0.696 | 0.684 |
| Linear | 0.241 | 0.286 | −0.957 | 0.0146 | 0.0155 | 0.092 | 0.691 | 0.681 |
| LineOffset | 0.211 | 0.242 | −0.977 | 0.0166 | 0.0197 | 0.104 | 0.697 | 0.682 |
| PerturbMean | 0.167 | 0.242 | −0.985 | 0.0178 | 0.0212 | 0.106 | 0.670 | 0.656 |
| ContextMean | 0.175 | 0.000 | −0.996 | 0.0168 | 0.0177 | 0.096 | 0.500 | 0.501 |
| CASTER, wrong-line controls | 0.127 | 0.181 | −0.993 | 0.0191 | 0.0208 | 0.102 | 0.589 | 0.577 |
| Perturbed mean | 0.304 | NA | −0.997 | 0.0150 | 0.0136 | 0.083 | 0.500 | 0.500 |
| NoChange | 0.039 | 0.127 | −0.997 | 0.0190 | 0.0196 | 0.099 | 0.500 | 0.500 |

**Table S9.** Calibrated metrics: DRF, scaled scores and CASTER versus Linear, related to Figure 3. At 32 cells (as rendered and noise-free) and against all cells (noise-free). DRF, mean of clipped per-knockdown values, and the ratio of means (which scaling the means in Table S8 gives). Scaled scores: 0 = negative control (NoChange for † metrics, the perturbed mean otherwise), 1 = technical duplicate, computed per line and seed and averaged; NA where the DRF is below 0.2, because the scale then has no meaning. Last columns: share of knockdowns for which CASTER, Linear or neither scored better, and the two-sided paired Wilcoxon p over knockdowns and seeds (approximate, see STAR Methods). Scores compare means and the test compares knockdowns, so they can disagree when a few knockdowns differ strongly (weighted R² is clipped at −1 and tied for many knockdowns). MSE, MAE and WMSE are lower-is-better.

| **Metric** | **Data** | **DRF (mean of per-knockdown values)** | **DRF (ratio of means)** | **Linear** | **CASTER** | **LineOffset** | **Perturb Mean** | **Context Mean** | **CASTER better (%)** | **Linear better (%)** | **Tied (%)** | **p** |
|---|---|---|---|---|---|---|---|---|---|---|---|---|
| Top-100 Pearson delta† | 32 cells, rendered | 0.11 | 0.38 | NA | NA | NA | NA | NA | 57 | 43 | 0 | 4.0 × 10<sup>−86</sup> |
| Top-100 Pearson delta† | 32 cells, noise-free | 0.11 | 0.38 | NA | NA | NA | NA | NA | 44 | 56 | 0 | 3.0 × 10<sup>−28</sup> |
| Top-100 Pearson delta† | All cells, noise-free | 0.38 | 0.66 | 0.84 | 0.80 | 0.49 | 0.44 | −0.27 | 41 | 59 | 0 | 5.5 × 10<sup>−61</sup> |
| Weighted R²† | 32 cells, rendered | 0.04 | 0.04 | NA | NA | NA | NA | NA | 3 | 2 | 94 | 2.7 × 10<sup>−8</sup> |
| Weighted R²† | 32 cells, noise-free | 0.04 | 0.04 | NA | NA | NA | NA | NA | 4 | 3 | 93 | 5.1 × 10<sup>−12</sup> |
| Weighted R²† | All cells, noise-free | 0.25 | 0.24 | 0.48 | 0.52 | 0.19 | 0.15 | −0.12 | 31 | 24 | 44 | 1.7 × 10<sup>−31</sup> |
| NIR | 32 cells, rendered | 0.67 | 0.80 | 0.48 | 0.49 | 0.50 | 0.43 | 0.00 | 50 | 48 | 2 | 5.4 × 10<sup>−4</sup> |
| NIR | 32 cells, noise-free | 0.67 | 0.80 | 0.49 | 0.49 | 0.51 | 0.44 | 0.00 | 48 | 50 | 2 | 0.214 |
| NIR | All cells, noise-free | 0.73 | 0.78 | 0.49 | 0.49 | 0.51 | 0.44 | 0.00 | 49 | 48 | 3 | 0.467 |
| PDS | 32 cells, rendered | 0.64 | 0.77 | 0.47 | 0.48 | 0.48 | 0.42 | 0.00 | 49 | 49 | 2 | 0.127 |
| PDS | 32 cells, noise-free | 0.64 | 0.77 | 0.48 | 0.48 | 0.49 | 0.42 | 0.00 | 47 | 51 | 2 | 1.5 × 10<sup>−4</sup> |
| PDS | All cells, noise-free | 0.68 | 0.71 | 0.47 | 0.47 | 0.47 | 0.41 | 0.00 | 47 | 50 | 3 | 0.008 |
| Pearson delta | 32 cells, rendered | −0.04 | −0.02 | NA | NA | NA | NA | NA | 47 | 53 | 0 | 3.0 × 10<sup>−17</sup> |
| Pearson delta | 32 cells, noise-free | −0.04 | −0.02 | NA | NA | NA | NA | NA | 40 | 60 | 0 | 3.1 × 10<sup>−110</sup> |
| Pearson delta | All cells, noise-free | 0.11 | 0.11 | NA | NA | NA | NA | NA | 39 | 61 | 0 | 8.2 × 10<sup>−145</sup> |
| Weighted R² | 32 cells, rendered | 0.06 | 0.04 | NA | NA | NA | NA | NA | 6 | 4 | 90 | 3.0 × 10<sup>−20</sup> |
| Weighted R² | 32 cells, noise-free | 0.06 | 0.04 | NA | NA | NA | NA | NA | 7 | 4 | 89 | 1.6 × 10<sup>−22</sup> |
| Weighted R² | All cells, noise-free | 0.32 | 0.26 | 0.51 | 0.56 | 0.26 | 0.20 | 0.00 | 32 | 24 | 44 | 2.0 × 10<sup>−54</sup> |
| MSE | 32 cells, rendered | −0.47 | −0.33 | NA | NA | NA | NA | NA | 50 | 50 | 0 | 6.6 × 10<sup>−5</sup> |
| MSE | 32 cells, noise-free | −0.47 | −0.33 | NA | NA | NA | NA | NA | 42 | 58 | 0 | 1.4 × 10<sup>−103</sup> |
| MSE | All cells, noise-free | −0.12 | 0.07 | NA | NA | NA | NA | NA | 40 | 60 | 0 | 2.1 × 10<sup>−126</sup> |
| MAE | 32 cells, rendered | −0.24 | −0.22 | NA | NA | NA | NA | NA | 48 | 52 | 0 | 3.8 × 10<sup>−10</sup> |
| MAE | 32 cells, noise-free | −0.24 | −0.22 | NA | NA | NA | NA | NA | 41 | 59 | 0 | 1.6 × 10<sup>−126</sup> |
| MAE | All cells, noise-free | −0.09 | −0.06 | NA | NA | NA | NA | NA | 39 | 61 | 0 | 3.9 × 10<sup>−171</sup> |
| WMSE | 32 cells, rendered | −0.08 | 0.07 | NA | NA | NA | NA | NA | 52 | 48 | 0 | 2.1 × 10<sup>−6</sup> |
| WMSE | 32 cells, noise-free | −0.08 | 0.07 | NA | NA | NA | NA | NA | 51 | 49 | 0 | 0.019 |
| WMSE | All cells, noise-free | 0.32 | 0.67 | 0.54 | 0.61 | 0.37 | 0.29 | 0.00 | 53 | 47 | 0 | 7.8 × 10<sup>−23</sup> |

**Table S10.** All metrics per target line, unseen-line task, related to Figure 4. One seed; mean over test knockdowns.

| **Cell line** | **Method** | **Pearson delta** | **Perturbation discrimination** | **MAE** | **DE AUPRC** | **DE overlap** | **Fold-change Spearman** | **Direction agreement** | **Effect-size Spearman** |
|---|---|---|---|---|---|---|---|---|---|
| HepG2 | CASTER | 0.169 | 0.191 | 0.081 | 0.128 | 0.109 | 0.331 | 0.715 | 0.672 |
| HepG2 | CASTER (noise-free) | 0.227 | 0.199 | 0.068 | 0.172 | 0.083 | 0.386 | 0.799 | 0.699 |
| HepG2 | NoChange | 0.060 | −0.008 | 0.079 | 0.081 | 0.000 | 0.099 | 0.549 | 0.010 |
| HepG2 | PerturbMean | 0.247 | 0.414 | 0.087 | 0.165 | 0.204 | 0.458 | 0.784 | 0.666 |
| HepG2 | PerturbMean (noise-free) | 0.301 | 0.438 | 0.074 | 0.191 | 0.186 | 0.506 | 0.826 | 0.684 |
| Jurkat | CASTER | 0.138 | 0.179 | 0.089 | 0.077 | 0.071 | 0.305 | 0.717 | 0.581 |
| Jurkat | CASTER (noise-free) | 0.252 | 0.176 | 0.070 | 0.148 | 0.058 | 0.395 | 0.822 | 0.610 |
| Jurkat | NoChange | 0.039 | 0.001 | 0.089 | 0.028 | 0.000 | 0.054 | 0.531 | −0.010 |
| Jurkat | PerturbMean | 0.155 | 0.335 | 0.097 | 0.107 | 0.144 | 0.352 | 0.756 | 0.635 |
| Jurkat | PerturbMean (noise-free) | 0.209 | 0.336 | 0.080 | 0.138 | 0.135 | 0.405 | 0.807 | 0.652 |
| K562 | CASTER | 0.187 | 0.401 | 0.081 | 0.093 | 0.105 | 0.409 | 0.769 | 0.635 |
| K562 | CASTER (noise-free) | 0.262 | 0.409 | 0.066 | 0.118 | 0.099 | 0.443 | 0.832 | 0.635 |
| K562 | NoChange | 0.051 | −0.001 | 0.075 | 0.029 | 0.000 | 0.067 | 0.552 | −0.009 |
| K562 | PerturbMean | 0.186 | 0.538 | 0.089 | 0.122 | 0.159 | 0.416 | 0.784 | 0.643 |
| K562 | PerturbMean (noise-free) | 0.228 | 0.559 | 0.075 | 0.146 | 0.175 | 0.458 | 0.808 | 0.657 |
| RPE1 | CASTER | 0.191 | 0.100 | 0.091 | 0.126 | 0.053 | 0.316 | 0.683 | 0.556 |
| RPE1 | CASTER (noise-free) | 0.308 | 0.095 | 0.075 | 0.214 | 0.039 | 0.439 | 0.778 | 0.579 |
| RPE1 | NoChange | 0.048 | 0.001 | 0.095 | 0.091 | 0.000 | 0.063 | 0.541 | 0.002 |
| RPE1 | PerturbMean | 0.205 | 0.185 | 0.099 | 0.146 | 0.118 | 0.363 | 0.715 | 0.533 |
| RPE1 | PerturbMean (noise-free) | 0.271 | 0.175 | 0.084 | 0.196 | 0.092 | 0.446 | 0.776 | 0.570 |

**Table S11.** Single-source transfer, related to Figure 4. Mean over the knockdowns shared by source and target line; one seed.

| **Source line** | **Target line** | **Method** | **Pearson delta** | **Perturbation discrimination** | **DE AUPRC** | **DE overlap** |
|---|---|---|---|---|---|---|
| HepG2 | Jurkat | CASTER | 0.127 | 0.153 | 0.054 | 0.066 |
| HepG2 | Jurkat | PerturbMean | 0.132 | 0.286 | 0.064 | 0.112 |
| HepG2 | K562 | CASTER | 0.160 | 0.287 | 0.077 | 0.096 |
| HepG2 | K562 | PerturbMean | 0.162 | 0.475 | 0.087 | 0.134 |
| HepG2 | RPE1 | CASTER | 0.246 | 0.138 | 0.154 | 0.089 |
| HepG2 | RPE1 | PerturbMean | 0.235 | 0.270 | 0.150 | 0.159 |
| Jurkat | HepG2 | CASTER | 0.157 | 0.117 | 0.098 | 0.090 |
| Jurkat | HepG2 | PerturbMean | 0.150 | 0.262 | 0.096 | 0.166 |
| Jurkat | K562 | CASTER | 0.152 | 0.313 | 0.090 | 0.104 |
| Jurkat | K562 | PerturbMean | 0.148 | 0.521 | 0.096 | 0.139 |
| Jurkat | RPE1 | CASTER | 0.143 | 0.068 | 0.107 | 0.047 |
| Jurkat | RPE1 | PerturbMean | 0.144 | 0.190 | 0.107 | 0.136 |
| K562 | HepG2 | CASTER | 0.164 | 0.066 | 0.107 | 0.059 |
| K562 | HepG2 | PerturbMean | 0.161 | 0.206 | 0.104 | 0.145 |
| K562 | Jurkat | CASTER | 0.119 | 0.096 | 0.073 | 0.057 |
| K562 | Jurkat | PerturbMean | 0.132 | 0.274 | 0.085 | 0.127 |
| K562 | RPE1 | CASTER | 0.135 | 0.027 | 0.118 | 0.033 |
| K562 | RPE1 | PerturbMean | 0.135 | 0.103 | 0.118 | 0.106 |
| RPE1 | HepG2 | CASTER | 0.272 | 0.411 | 0.180 | 0.201 |
| RPE1 | HepG2 | PerturbMean | 0.262 | 0.558 | 0.177 | 0.230 |
| RPE1 | Jurkat | CASTER | 0.130 | 0.211 | 0.084 | 0.100 |
| RPE1 | Jurkat | PerturbMean | 0.136 | 0.363 | 0.100 | 0.140 |
| RPE1 | K562 | CASTER | 0.149 | 0.283 | 0.087 | 0.101 |
| RPE1 | K562 | PerturbMean | 0.150 | 0.430 | 0.102 | 0.140 |

**Table S12.** Complex-grouped versus random splits, related to Figure 5. Scaled and raw top-100 Pearson delta against all cells (noise-free predictions) per line, seed, split and test set; matched = knockdowns held out under both splits (same line and seed). The perturbed mean depends on the screened set, so the zero of the scaled score differs between splits. Change in CASTER minus Linear (grouped minus random, matched knockdowns): scaled 0.04 (bootstrap 95% interval 0.03 to 0.05), raw 0.02 (0.02 to 0.03).

| **Line** | **Seed** | **Split** | **Test set** | **Knockdowns** | **CASTER scaled (raw)** | **Linear scaled (raw)** | **LineOffset scaled (raw)** | **PerturbMean scaled (raw)** |
|---|---|---|---|---|---|---|---|---|
| HepG2 | 0 | grouped | full | 894 | 0.82 (0.58) | 0.83 (0.58) | 0.53 (0.40) | 0.55 (0.41) |
| HepG2 | 0 | random | full | 894 | 0.88 (0.64) | 0.91 (0.66) | 0.44 (0.42) | 0.48 (0.45) |
| HepG2 | 0 | grouped | matched | 629 | 0.82 (0.58) | 0.83 (0.58) | 0.51 (0.39) | 0.53 (0.41) |
| HepG2 | 0 | random | matched | 629 | 0.89 (0.65) | 0.92 (0.67) | 0.48 (0.42) | 0.50 (0.43) |
| HepG2 | 1 | grouped | full | 845 | 0.88 (0.64) | 0.86 (0.62) | 0.50 (0.42) | 0.54 (0.44) |
| HepG2 | 1 | random | full | 894 | 0.88 (0.63) | 0.90 (0.64) | 0.46 (0.40) | 0.50 (0.43) |
| HepG2 | 1 | grouped | matched | 595 | 0.88 (0.64) | 0.84 (0.62) | 0.49 (0.42) | 0.53 (0.45) |
| HepG2 | 1 | random | matched | 595 | 0.89 (0.65) | 0.91 (0.66) | 0.49 (0.42) | 0.53 (0.44) |
| HepG2 | 2 | grouped | full | 821 | 0.82 (0.61) | 0.84 (0.62) | 0.40 (0.41) | 0.48 (0.44) |
| HepG2 | 2 | random | full | 894 | 0.88 (0.63) | 0.92 (0.65) | 0.45 (0.41) | 0.50 (0.43) |
| HepG2 | 2 | grouped | matched | 589 | 0.83 (0.61) | 0.86 (0.63) | 0.40 (0.40) | 0.47 (0.44) |
| HepG2 | 2 | random | matched | 589 | 0.91 (0.64) | 0.93 (0.65) | 0.44 (0.40) | 0.48 (0.42) |
| Jurkat | 0 | grouped | full | 987 | 0.71 (0.50) | 0.73 (0.51) | 0.41 (0.37) | 0.44 (0.38) |
| Jurkat | 0 | random | full | 1,027 | 0.82 (0.53) | 0.84 (0.54) | 0.53 (0.38) | 0.54 (0.38) |
| Jurkat | 0 | grouped | matched | 713 | 0.72 (0.51) | 0.75 (0.52) | 0.45 (0.38) | 0.46 (0.39) |
| Jurkat | 0 | random | matched | 713 | 0.80 (0.53) | 0.82 (0.54) | 0.49 (0.37) | 0.49 (0.37) |
| Jurkat | 1 | grouped | full | 1,023 | 0.73 (0.54) | 0.69 (0.52) | 0.36 (0.40) | 0.47 (0.44) |
| Jurkat | 1 | random | full | 1,027 | 0.81 (0.52) | 0.85 (0.54) | 0.52 (0.37) | 0.53 (0.37) |
| Jurkat | 1 | grouped | matched | 717 | 0.75 (0.54) | 0.69 (0.52) | 0.40 (0.40) | 0.49 (0.44) |
| Jurkat | 1 | random | matched | 717 | 0.75 (0.48) | 0.80 (0.51) | 0.52 (0.37) | 0.54 (0.38) |
| Jurkat | 2 | grouped | full | 930 | 0.80 (0.52) | 0.79 (0.52) | 0.52 (0.38) | 0.55 (0.40) |
| Jurkat | 2 | random | full | 1,027 | 0.82 (0.54) | 0.87 (0.56) | 0.48 (0.38) | 0.50 (0.39) |
| Jurkat | 2 | grouped | matched | 658 | 0.81 (0.53) | 0.80 (0.53) | 0.52 (0.39) | 0.55 (0.40) |
| Jurkat | 2 | random | matched | 658 | 0.81 (0.54) | 0.84 (0.56) | 0.53 (0.40) | 0.55 (0.41) |
| K562 | 0 | grouped | full | 890 | 0.77 (0.56) | 0.74 (0.54) | 0.54 (0.45) | 0.39 (0.39) |
| K562 | 0 | random | full | 917 | 0.84 (0.58) | 0.87 (0.59) | 0.55 (0.43) | 0.42 (0.37) |
| K562 | 0 | grouped | matched | 627 | 0.77 (0.56) | 0.74 (0.54) | 0.56 (0.45) | 0.43 (0.39) |
| K562 | 0 | random | matched | 627 | 0.83 (0.58) | 0.83 (0.58) | 0.56 (0.44) | 0.44 (0.38) |
| K562 | 1 | grouped | full | 903 | 0.76 (0.60) | 0.75 (0.60) | 0.31 (0.47) | 0.13 (0.41) |
| K562 | 1 | random | full | 917 | 0.87 (0.59) | 0.92 (0.62) | 0.53 (0.45) | 0.38 (0.38) |
| K562 | 1 | grouped | matched | 646 | 0.79 (0.60) | 0.79 (0.60) | 0.30 (0.47) | 0.12 (0.42) |
| K562 | 1 | random | matched | 646 | 0.84 (0.57) | 0.91 (0.59) | 0.43 (0.43) | 0.23 (0.35) |
| K562 | 2 | grouped | full | 871 | 0.73 (0.54) | 0.74 (0.54) | 0.57 (0.46) | 0.44 (0.39) |
| K562 | 2 | random | full | 917 | 0.82 (0.58) | 0.88 (0.61) | 0.53 (0.44) | 0.37 (0.37) |
| K562 | 2 | grouped | matched | 602 | 0.73 (0.54) | 0.73 (0.54) | 0.57 (0.46) | 0.44 (0.39) |
| K562 | 2 | random | matched | 602 | 0.79 (0.58) | 0.85 (0.62) | 0.57 (0.47) | 0.44 (0.40) |
| RPE1 | 0 | grouped | full | 960 | 0.62 (0.53) | 0.62 (0.54) | 0.47 (0.42) | 0.33 (0.31) |
| RPE1 | 0 | random | full | 1,003 | 0.66 (0.55) | 0.71 (0.59) | 0.48 (0.41) | 0.35 (0.31) |
| RPE1 | 0 | grouped | matched | 677 | 0.61 (0.53) | 0.62 (0.54) | 0.47 (0.42) | 0.32 (0.31) |
| RPE1 | 0 | random | matched | 677 | 0.65 (0.55) | 0.70 (0.58) | 0.48 (0.42) | 0.33 (0.30) |
| RPE1 | 1 | grouped | full | 930 | 0.55 (0.49) | 0.61 (0.54) | 0.44 (0.41) | 0.32 (0.32) |
| RPE1 | 1 | random | full | 1,003 | 0.68 (0.57) | 0.72 (0.60) | 0.48 (0.41) | 0.35 (0.31) |
| RPE1 | 1 | grouped | matched | 646 | 0.58 (0.52) | 0.62 (0.55) | 0.44 (0.42) | 0.33 (0.33) |
| RPE1 | 1 | random | matched | 646 | 0.66 (0.56) | 0.70 (0.60) | 0.45 (0.41) | 0.35 (0.33) |
| RPE1 | 2 | grouped | full | 991 | 0.58 (0.54) | 0.55 (0.53) | 0.39 (0.42) | 0.30 (0.36) |
| RPE1 | 2 | random | full | 1,003 | 0.66 (0.55) | 0.68 (0.57) | 0.47 (0.40) | 0.35 (0.30) |
| RPE1 | 2 | grouped | matched | 701 | 0.58 (0.55) | 0.54 (0.52) | 0.38 (0.42) | 0.30 (0.36) |
| RPE1 | 2 | random | matched | 701 | 0.63 (0.55) | 0.66 (0.58) | 0.41 (0.40) | 0.33 (0.34) |

**Table S13.** Gene programs and their top Reactome enrichment, related to Figure 6. Programs P0 to P14 per line; BH-adjusted p of the top Reactome term (hypergeometric test); NA, no term at FDR < 0.05.

| **Cell line** | **Program** | **Number of genes** | **Top Reactome pathway** | **BH-adjusted p** |
|---|---|---|---|---|
| HepG2 | P0 | 199 | Cell Cycle | 5.0 × 10<sup>−23</sup> |
| HepG2 | P1 | 85 | Metabolism | 4.6 × 10<sup>−5</sup> |
| HepG2 | P2 | 127 | Metabolism of RNA | 4.6 × 10<sup>−5</sup> |
| HepG2 | P3 | 78 | Generic Transcription Pathway | 6.0 × 10<sup>−5</sup> |
| HepG2 | P4 | 193 | NA |  |
| HepG2 | P5 | 180 | Anchoring of the basal body to the plasma membrane | 5.9 × 10<sup>−5</sup> |
| HepG2 | P6 | 83 | Metabolism | 9.0 × 10<sup>−6</sup> |
| HepG2 | P7 | 161 | NA |  |
| HepG2 | P8 | 133 | NA |  |
| HepG2 | P9 | 142 | NA |  |
| HepG2 | P10 | 68 | DNA Double Strand Break Response | 4.5 × 10<sup>−5</sup> |
| HepG2 | P11 | 170 | NA |  |
| HepG2 | P12 | 186 | NA |  |
| HepG2 | P13 | 86 | NA |  |
| HepG2 | P14 | 109 | Cholesterol biosynthesis | 7.5 × 10<sup>−7</sup> |
| Jurkat | P0 | 13 | NA |  |
| Jurkat | P1 | 183 | NA |  |
| Jurkat | P2 | 62 | Cell Cycle, Mitotic | 1.1 × 10<sup>−17</sup> |
| Jurkat | P3 | 271 | Gene expression (Transcription) | 6.8 × 10<sup>−5</sup> |
| Jurkat | P4 | 229 | NA |  |
| Jurkat | P5 | 106 | Cell Cycle | 9.4 × 10<sup>−12</sup> |
| Jurkat | P6 | 151 | NA |  |
| Jurkat | P7 | 122 | Metabolism of RNA | 3.0 × 10<sup>−8</sup> |
| Jurkat | P8 | 100 | Cholesterol biosynthesis | 1.2 × 10<sup>−8</sup> |
| Jurkat | P9 | 37 | NA |  |
| Jurkat | P10 | 113 | Muscle contraction | 4.4 × 10<sup>−5</sup> |
| Jurkat | P11 | 183 | FCGR3A-mediated phagocytosis | 8.3 × 10<sup>−5</sup> |
| Jurkat | P12 | 246 | NA |  |
| Jurkat | P13 | 59 | NA |  |
| Jurkat | P14 | 125 | NA |  |
| K562 | P0 | 29 | NA |  |
| K562 | P1 | 129 | Cholesterol biosynthesis | 4.3 × 10<sup>−5</sup> |
| K562 | P2 | 152 | NA |  |
| K562 | P3 | 152 | NA |  |
| K562 | P4 | 264 | NA |  |
| K562 | P5 | 108 | NA |  |
| K562 | P6 | 256 | NA |  |
| K562 | P7 | 79 | Autophagy | 4.1 × 10<sup>−5</sup> |
| K562 | P8 | 172 | NA |  |
| K562 | P9 | 48 | NA |  |
| K562 | P10 | 128 | Immune System | 1.6 × 10<sup>−7</sup> |
| K562 | P11 | 104 | NA |  |
| K562 | P12 | 55 | Cell Cycle | 1.4 × 10<sup>−14</sup> |
| K562 | P13 | 132 | Metabolism of RNA | 1.6 × 10<sup>−5</sup> |
| K562 | P14 | 192 | NA |  |
| RPE1 | P0 | 38 | Cellular response to chemical stress | 7.5 × 10<sup>−7</sup> |
| RPE1 | P1 | 285 | NA |  |
| RPE1 | P2 | 234 | NA |  |
| RPE1 | P3 | 109 | NA |  |
| RPE1 | P4 | 29 | NA |  |
| RPE1 | P5 | 155 | NA |  |
| RPE1 | P6 | 165 | NA |  |
| RPE1 | P7 | 70 | Asparagine N-linked glycosylation | 3.3 × 10<sup>−8</sup> |
| RPE1 | P8 | 283 | Cell Cycle | 7.2 × 10<sup>−28</sup> |
| RPE1 | P9 | 47 | NA |  |
| RPE1 | P10 | 149 | NA |  |
| RPE1 | P11 | 93 | NA |  |
| RPE1 | P12 | 133 | Intraflagellar transport | 2.7 × 10<sup>−5</sup> |
| RPE1 | P13 | 186 | NA |  |
| RPE1 | P14 | 24 | NA |  |

**Table S14.**
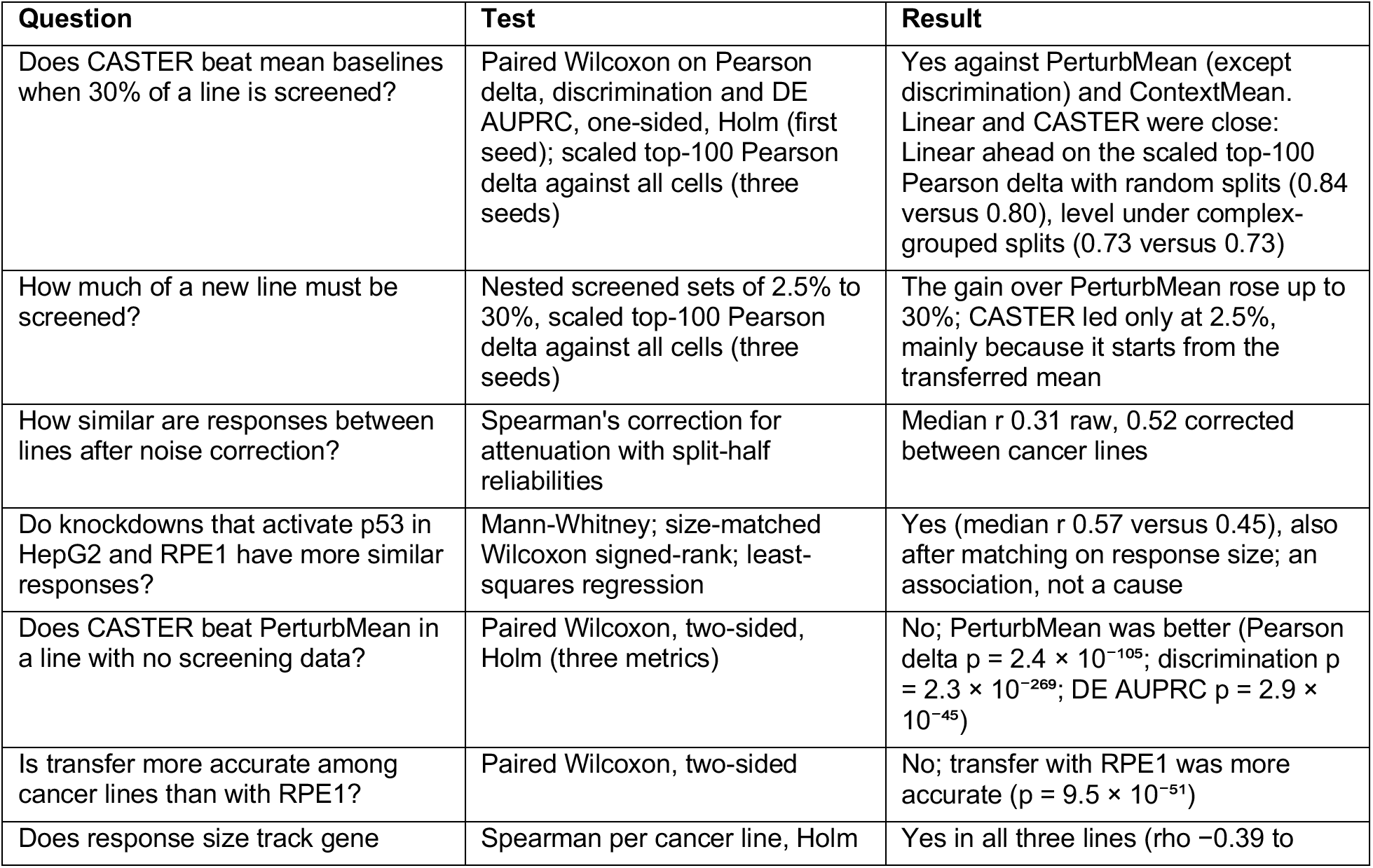
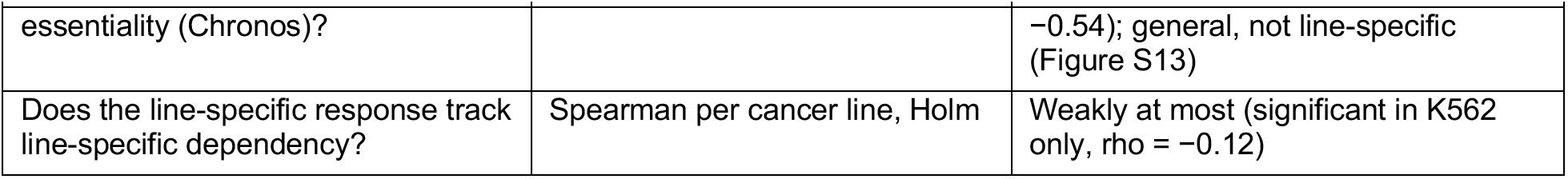
Main questions, tests and outcomes, related to Figures 2 to 6.

**Table S15.** Cross-line response similarity before and after correction for measurement noise, related to Figure 1. Knockdowns with ≥ 10 DEGs in both lines; medians over knockdowns.

| **Line pair** | **n** | **Raw r (32 cells)** | **Reliability, line 1** | **Reliability, line 2** | **Corrected r (32 cells)** | **Raw r (all cells)** | **Corrected r (all cells)** |
|---|---|---|---|---|---|---|---|
| HepG2-Jurkat | 289 | 0.28 | 0.77 | 0.57 | 0.48 | 0.38 | 0.49 |
| HepG2-RPE1 | 400 | 0.52 | 0.75 | 0.85 | 0.66 | 0.57 | 0.66 |
| Jurkat-RPE1 | 311 | 0.29 | 0.54 | 0.83 | 0.47 | 0.37 | 0.48 |
| K562-HepG2 | 287 | 0.32 | 0.59 | 0.79 | 0.50 | 0.42 | 0.50 |
| K562-Jurkat | 237 | 0.33 | 0.57 | 0.58 | 0.59 | 0.47 | 0.59 |
| K562-RPE1 | 277 | 0.31 | 0.59 | 0.86 | 0.47 | 0.39 | 0.47 |

**Table S16.** All metrics per target line, partial-screen task, related to Figure 2. First seed; mean over held-out knockdowns.

| Cell line | Method | Pearson delta | Perturbation discrimination | MAE | DE AUPRC | DE overlap | Fold-change Spearman | Direction agreement | Effect-size Spearman |
|---|---|---|---|---|---|---|---|---|---|
| HepG2 | CASTER | 0.315 | 0.470 | 0.070 | 0.227 | 0.177 | 0.506 | 0.813 | 0.703 |
| HepG2 | CASTER (noise-free) | 0.455 | 0.489 | 0.055 | 0.333 | 0.151 | 0.546 | 0.898 | 0.736 |
| HepG2 | ContextMean | 0.254 | −0.007 | 0.076 | 0.114 | 0.016 | 0.428 | 0.730 | −0.003 |
| HepG2 | ContextMean (noise-free) | 0.393 | 0.000 | 0.061 | 0.251 | 0.000 | 0.519 | 0.840 | 0.000 |
| HepG2 | LineOffset | 0.273 | 0.461 | 0.085 | 0.174 | 0.207 | 0.501 | 0.808 | 0.651 |
| HepG2 | LineOffset (noise-free) | 0.330 | 0.496 | 0.072 | 0.209 | 0.195 | 0.536 | 0.850 | 0.675 |
| HepG2 | Linear | 0.322 | 0.478 | 0.069 | 0.210 | 0.151 | 0.488 | 0.814 | 0.703 |
| HepG2 | Linear (noise-free) | 0.468 | 0.506 | 0.054 | 0.324 | 0.118 | 0.540 | 0.895 | 0.733 |
| HepG2 | NoChange | 0.060 | −0.012 | 0.078 | 0.078 | 0.000 | 0.103 | 0.549 | 0.017 |
| HepG2 | PerturbMean | 0.243 | 0.423 | 0.086 | 0.162 | 0.201 | 0.445 | 0.785 | 0.671 |
| HepG2 | PerturbMean (noise-free) | 0.297 | 0.452 | 0.074 | 0.188 | 0.186 | 0.490 | 0.824 | 0.686 |
| Jurkat | CASTER | 0.205 | 0.378 | 0.085 | 0.123 | 0.099 | 0.386 | 0.777 | 0.575 |
| Jurkat | CASTER (noise-free) | 0.364 | 0.379 | 0.066 | 0.234 | 0.076 | 0.446 | 0.904 | 0.597 |
| Jurkat | ContextMean | 0.113 | 0.002 | 0.088 | 0.039 | 0.002 | 0.221 | 0.627 | −0.040 |
| Jurkat | ContextMean (noise-free) | 0.296 | 0.000 | 0.068 | 0.130 | 0.000 | 0.391 | 0.793 | 0.000 |
| Jurkat | LineOffset | 0.181 | 0.388 | 0.096 | 0.121 | 0.157 | 0.403 | 0.783 | 0.620 |
| Jurkat | LineOffset (noise-free) | 0.247 | 0.393 | 0.078 | 0.159 | 0.140 | 0.440 | 0.835 | 0.647 |
| Jurkat | Linear | 0.197 | 0.349 | 0.085 | 0.101 | 0.073 | 0.367 | 0.766 | 0.529 |
| Jurkat | Linear (noise-free) | 0.370 | 0.361 | 0.065 | 0.216 | 0.047 | 0.444 | 0.887 | 0.520 |
| Jurkat | NoChange | 0.039 | 0.001 | 0.090 | 0.029 | 0.000 | 0.060 | 0.516 | −0.050 |
| Jurkat | PerturbMean | 0.160 | 0.348 | 0.098 | 0.112 | 0.148 | 0.354 | 0.757 | 0.650 |
| Jurkat | PerturbMean (noise-free) | 0.216 | 0.350 | 0.080 | 0.143 | 0.139 | 0.404 | 0.813 | 0.667 |
| K562 | CASTER | 0.205 | 0.408 | 0.072 | 0.145 | 0.115 | 0.426 | 0.815 | 0.617 |
| K562 | CASTER (noise-free) | 0.370 | 0.422 | 0.055 | 0.261 | 0.093 | 0.506 | 0.923 | 0.638 |
| K562 | ContextMean | 0.101 | −0.003 | 0.075 | 0.036 | 0.001 | 0.276 | 0.655 | 0.043 |
| K562 | ContextMean (noise-free) | 0.283 | 0.000 | 0.058 | 0.107 | 0.000 | 0.508 | 0.808 | 0.000 |
| K562 | LineOffset | 0.183 | 0.543 | 0.086 | 0.133 | 0.172 | 0.447 | 0.808 | 0.626 |
| K562 | LineOffset (noise-free) | 0.231 | 0.560 | 0.072 | 0.159 | 0.179 | 0.494 | 0.833 | 0.646 |
| K562 | Linear | 0.209 | 0.429 | 0.072 | 0.133 | 0.101 | 0.450 | 0.805 | 0.626 |
| K562 | Linear (noise-free) | 0.378 | 0.450 | 0.055 | 0.271 | 0.065 | 0.503 | 0.916 | 0.616 |
| K562 | NoChange | 0.049 | −0.002 | 0.076 | 0.029 | 0.000 | 0.069 | 0.553 | −0.005 |
| K562 | PerturbMean | 0.189 | 0.537 | 0.089 | 0.121 | 0.158 | 0.436 | 0.789 | 0.643 |
| K562 | PerturbMean (noise-free) | 0.232 | 0.558 | 0.075 | 0.147 | 0.176 | 0.475 | 0.814 | 0.656 |
| RPE1 | CASTER | 0.386 | 0.468 | 0.082 | 0.276 | 0.205 | 0.501 | 0.812 | 0.668 |
| RPE1 | CASTER (noise-free) | 0.507 | 0.460 | 0.065 | 0.388 | 0.183 | 0.563 | 0.891 | 0.694 |
| RPE1 | ContextMean | 0.361 | 0.006 | 0.088 | 0.198 | 0.065 | 0.480 | 0.773 | −0.009 |
| RPE1 | ContextMean (noise-free) | 0.488 | 0.000 | 0.072 | 0.344 | 0.027 | 0.549 | 0.856 | −0.133 |
| RPE1 | LineOffset | 0.365 | 0.379 | 0.094 | 0.219 | 0.198 | 0.533 | 0.828 | 0.543 |
| RPE1 | LineOffset (noise-free) | 0.442 | 0.393 | 0.079 | 0.292 | 0.164 | 0.585 | 0.871 | 0.622 |
| RPE1 | Linear | 0.399 | 0.438 | 0.081 | 0.259 | 0.171 | 0.503 | 0.823 | 0.701 |
| RPE1 | Linear (noise-free) | 0.531 | 0.446 | 0.063 | 0.386 | 0.139 | 0.559 | 0.896 | 0.736 |
| RPE1 | NoChange | 0.050 | 0.002 | 0.095 | 0.092 | 0.000 | 0.071 | 0.541 | −0.010 |
| RPE1 | PerturbMean | 0.204 | 0.190 | 0.098 | 0.148 | 0.121 | 0.364 | 0.710 | 0.544 |
| RPE1 | PerturbMean (noise-free) | 0.268 | 0.176 | 0.084 | 0.199 | 0.094 | 0.441 | 0.771 | 0.580 |

**Table S17.** All paired tests, partial-screen task, related to Figure 2. One-sided paired Wilcoxon signed-rank tests (CASTER better than the baseline); Holm correction within each family: the three main metrics against the three baselines (first seed, n = 3,841 knockdowns over four lines), the secondary metrics against each baseline (three seeds pooled, n = 11,523), and the predicted-side DE test.

| Baseline | Metric | Predicted-side DE test | n | CASTER mean | Baseline mean | Knockdowns with CASTER better (%) | p | Holm-adjusted p |
|---|---|---|---|---|---|---|---|---|
| PerturbMean | Pearson delta | Moment-matched | 3,841 | 0.278 | 0.198 | 66 | 5.8 × 10<sup>−153</sup> | 4.0 × 10<sup>−152</sup> |
| PerturbMean | Perturbation discrimination | Moment-matched | 3,841 | 0.430 | 0.369 | 46 | 0.648 | 1.000 |
| PerturbMean | DE AUPRC | Moment-matched | 2,666 | 0.198 | 0.137 | 67 | 1.2 × 10<sup>−103</sup> | 6.1 × 10<sup>−103</sup> |
| ContextMean | Pearson delta | Moment-matched | 3,841 | 0.278 | 0.208 | 67 | 3.3 × 10<sup>−150</sup> | 2.0 × 10<sup>−149</sup> |
| ContextMean | Perturbation discrimination | Moment-matched | 3,841 | 0.430 | 0.000 | 80 | < 10<sup>−300</sup> | < 10<sup>−300</sup> |
| ContextMean | DE AUPRC | Moment-matched | 2,666 | 0.198 | 0.104 | 77 | 2.3 × 10<sup>−223</sup> | 1.8 × 10<sup>−222</sup> |
| Linear | Pearson delta | Moment-matched | 3,841 | 0.278 | 0.282 | 50 | 0.960 | 1.000 |
| Linear | Perturbation discrimination | Moment-matched | 3,841 | 0.430 | 0.422 | 50 | 0.039 | 0.117 |
| Linear | DE AUPRC | Moment-matched | 2,666 | 0.198 | 0.181 | 56 | 4.0 × 10<sup>−18</sup> | 1.6 × 10<sup>−17</sup> |
| PerturbMean | MAE | Moment-matched | 11,523 | 0.078 | 0.093 | 98 | < 10<sup>−300</sup> | < 10<sup>−300</sup> |
| PerturbMean | DE overlap | Moment-matched | 7,977 | 0.149 | 0.150 | 30 | 0.577 | 1.000 |
| PerturbMean | Fold-change Spearman | Moment-matched | 6,267 | 0.462 | 0.396 | 59 | 8.1 × 10<sup>−120</sup> | 4.9 × 10<sup>−119</sup> |
| PerturbMean | Direction agreement | Moment-matched | 7,977 | 0.808 | 0.755 | 52 | 1.8 × 10<sup>−250</sup> | 1.3 × 10<sup>−249</sup> |
| ContextMean | MAE | Moment-matched | 11,523 | 0.078 | 0.082 | 75 | < 10<sup>−300</sup> | < 10<sup>−300</sup> |
| ContextMean | DE overlap | Moment-matched | 7,977 | 0.149 | 0.023 | 47 | < 10<sup>−300</sup> | < 10<sup>−300</sup> |
| ContextMean | Fold-change Spearman | Moment-matched | 6,267 | 0.462 | 0.362 | 70 | < 10<sup>−300</sup> | < 10<sup>−300</sup> |
| ContextMean | Direction agreement | Moment-matched | 7,977 | 0.808 | 0.702 | 60 | < 10<sup>−300</sup> | < 10<sup>−300</sup> |
| Linear | MAE | Moment-matched | 11,523 | 0.078 | 0.077 | 48 | 1.000 | 1.000 |
| Linear | DE overlap | Moment-matched | 7,977 | 0.149 | 0.126 | 30 | 1.1 × 10<sup>−50</sup> | 5.3 × 10<sup>−50</sup> |
| Linear | Fold-change Spearman | Moment-matched | 6,267 | 0.462 | 0.459 | 46 | 0.052 | 0.208 |
| Linear | Direction agreement | Moment-matched | 7,977 | 0.808 | 0.803 | 28 | 1.000 | 1.000 |
| PerturbMean | DE AUPRC | Wilcoxon rank test | 2,666 | 0.039 | 0.035 | 73 | 1.6 × 10<sup>−152</sup> | 1.6 × 10<sup>−152</sup> |
| ContextMean | DE AUPRC | Wilcoxon rank test | 2,666 | 0.039 | 0.036 | 85 | 5.8 × 10<sup>−246</sup> | 1.7 × 10<sup>−245</sup> |
| Linear | DE AUPRC | Wilcoxon rank test | 2,666 | 0.039 | 0.037 | 85 | 5.1 × 10<sup>−244</sup> | 1.0 × 10<sup>−243</sup> |
| PerturbMean | DE overlap | Wilcoxon rank test | 2,666 | 0.078 | 0.055 | 28 | 5.4 × 10<sup>−25</sup> | 5.4 × 10<sup>−25</sup> |
| ContextMean | DE overlap | Wilcoxon rank test | 2,666 | 0.078 | 0.034 | 35 | 6.2 × 10<sup>−120</sup> | 1.9 × 10<sup>−119</sup> |
| Linear | DE overlap | Wilcoxon rank test | 2,666 | 0.078 | 0.066 | 28 | 1.2 × 10<sup>−48</sup> | 2.3 × 10<sup>−48</sup> |
| PerturbMean | Fold-change Spearman | Wilcoxon rank test | 2,094 | 0.509 | 0.435 | 62 | 2.7 × 10<sup>−50</sup> | 5.4 × 10<sup>−50</sup> |
| ContextMean | Fold-change Spearman | Wilcoxon rank test | 2,094 | 0.509 | 0.342 | 76 | 2.1 × 10<sup>−182</sup> | 6.2 × 10<sup>−182</sup> |
| Linear | Fold-change Spearman | Wilcoxon rank test | 2,094 | 0.509 | 0.496 | 53 | 5.4 × 10<sup>−14</sup> | 5.4 × 10<sup>−14</sup> |
| PerturbMean | Direction agreement | Wilcoxon rank test | 2,666 | 0.796 | 0.756 | 49 | 4.6 × 10<sup>−65</sup> | 9.2 × 10<sup>−65</sup> |
| ContextMean | Direction agreement | Wilcoxon rank test | 2,666 | 0.796 | 0.691 | 60 | 4.0 × 10<sup>−192</sup> | 1.2 × 10<sup>−191</sup> |
| Linear | Direction agreement | Wilcoxon rank test | 2,666 | 0.796 | 0.793 | 30 | 0.073 | 0.073 |

**Table S18.** Partial-screen task scored against all cells of each held-out knockdown, related to Figure 3. Noise-free predictions; mean over lines and seeds. With all cells, 5% to 11% of genes are differentially expressed per knockdown; DE metrics use the moment-matched test with each knockdown’s full cell count. NoChange is rendered on sampled control cells, so sampling noise gives it a few predicted DEGs and a small nonzero DE overlap and fold-change Spearman.

| **Method** | **Pearson delta** | **Perturbation discrimination** | **MAE** | **DE AUPRC** | **DE overlap** | **Fold-change Spearman** |
|---|---|---|---|---|---|---|
| CASTER | 0.511 | 0.401 | 0.045 | 0.344 | 0.170 | 0.507 |
| Linear | 0.528 | 0.408 | 0.044 | 0.349 | 0.150 | 0.500 |
| LineOffset | 0.370 | 0.418 | 0.064 | 0.230 | 0.233 | 0.491 |
| PerturbMean | 0.305 | 0.350 | 0.067 | 0.195 | 0.201 | 0.438 |
| ContextMean | 0.438 | 0.000 | 0.050 | 0.272 | 0.035 | 0.442 |
| NoChange | 0.065 | −0.002 | 0.075 | 0.094 | 0.045 | 0.081 |

**Table S19.** Information available to each method, related to STAR Methods. Control cells of each line are split once (first seed) into disjoint model-input and reference control cells. Gene selection uses control cells only. Every knockdown in the held-out test set is excluded from all training and model-selection steps for that line. The code checks these rules at run time.

| **Method** | **Target-line control cells** | **Screened knockdowns of the target line** | **Source lines** | **Test knockdowns of the target line** |
|---|---|---|---|---|
| NoChange | model-input control cells | no | no | never |
| ContextMean | model-input control cells | yes (mean) | no | never |
| PerturbMean | model-input control cells | no | yes | never |
| Linear | model-input control cells | yes (fit) | yes | never |
| CASTER (partial screen) | model-input control cells | yes (training and validation) | yes | never |
| CASTER (unseen line, single source) | model-input control cells | no | yes (5% used for validation) | never |
| Evaluation | reference control cells | no | no | observed cells |

**Table S20.** Split sizes and training summary, related to STAR Methods. Training and validation counts are knockdowns of the target line (zero in the unseen-line task). Run time is the time to train and evaluate; one grouped run spans a suspension of the computer, and that epoch is counted at the median epoch duration. Learning-curve, single-source and ablation runs are not listed (learning-curve sizes in Table S5).

| Task | Target line | Seed | Training knockdowns | Validation knockdowns | Test knockdowns | Selected epoch | Run time (min) |
|---|---|---|---|---|---|---|---|
| Partial screen | HepG2 | 0 | 320 | 64 | 894 | 18 | 21.6 |
| Partial screen | HepG2 | 1 | 320 | 64 | 894 | 16 | 46.2 |
| Partial screen | HepG2 | 2 | 320 | 64 | 894 | 15 | 40.3 |
| Partial screen | Jurkat | 0 | 366 | 73 | 1,027 | 18 | 36.3 |
| Partial screen | Jurkat | 1 | 366 | 73 | 1,027 | 11 | 44.0 |
| Partial screen | Jurkat | 2 | 366 | 73 | 1,027 | 14 | 31.7 |
| Partial screen | K562 | 0 | 327 | 65 | 917 | 18 | 36.0 |
| Partial screen | K562 | 1 | 327 | 65 | 917 | 16 | 62.1 |
| Partial screen | K562 | 2 | 327 | 65 | 917 | 16 | 32.6 |
| Partial screen | RPE1 | 0 | 358 | 72 | 1,003 | 14 | 31.3 |
| Partial screen | RPE1 | 1 | 358 | 72 | 1,003 | 9 | 47.8 |
| Partial screen | RPE1 | 2 | 358 | 72 | 1,003 | 13 | 31.8 |
| Unseen line | HepG2 | 0 | 0 | 0 | 1,278 | 15 | 29.7 |
| Unseen line | Jurkat | 0 | 0 | 0 | 1,466 | 6 | 35.3 |
| Unseen line | K562 | 0 | 0 | 0 | 1,309 | 17 | 33.7 |
| Unseen line | RPE1 | 0 | 0 | 0 | 1,433 | 18 | 19.3 |
| Partial screen, mean-profile CASTER | HepG2 | 0 | 320 | 64 | 894 | 14 | 33.1 |
| Partial screen, mean-profile CASTER | Jurkat | 0 | 366 | 73 | 1,027 | 16 | 31.9 |
| Partial screen, mean-profile CASTER | K562 | 0 | 327 | 65 | 917 | 20 | 32.3 |
| Partial screen, mean-profile CASTER | RPE1 | 0 | 358 | 72 | 1,003 | 8 | 32.8 |
| Partial screen, line-label CASTER | HepG2 | 0 | 320 | 64 | 894 | 19 | 33.8 |
| Partial screen, line-label CASTER | Jurkat | 0 | 366 | 73 | 1,027 | 20 | 41.3 |
| Partial screen, line-label CASTER | K562 | 0 | 327 | 65 | 917 | 20 | 32.1 |
| Partial screen, line-label CASTER | RPE1 | 0 | 358 | 72 | 1,003 | 16 | 29.1 |
| Partial screen, grouped split | HepG2 | 0 | 320 | 64 | 894 | 15 | 33.0 |
| Partial screen, grouped split | HepG2 | 1 | 343 | 90 | 845 | 12 | 34.1 |
| Partial screen, grouped split | HepG2 | 2 | 393 | 64 | 821 | 16 | 42.8 |
| Partial screen, grouped split | Jurkat | 0 | 396 | 83 | 987 | 19 | 32.3 |
| Partial screen, grouped split | Jurkat | 1 | 369 | 74 | 1,023 | 14 | 32.6 |
| Partial screen, grouped split | Jurkat | 2 | 462 | 74 | 930 | 17 | 34.4 |
| Partial screen, grouped split | K562 | 0 | 337 | 82 | 890 | 18 | 32.9 |
| Partial screen, grouped split | K562 | 1 | 335 | 71 | 903 | 18 | 33.6 |
| Partial screen, grouped split | K562 | 2 | 351 | 87 | 871 | 19 | 40.6 |
| Partial screen, grouped split | RPE1 | 0 | 377 | 96 | 960 | 12 | 48.9 |
| Partial screen, grouped split | RPE1 | 1 | 409 | 94 | 930 | 10 | 33.4 |
| Partial screen, grouped split | RPE1 | 2 | 369 | 73 | 991 | 8 | 39.6 |

## Notes

### Competing Interest Statement

The authors have declared no competing interest.

